# Contingency and determinism in rapid adaptation shape evolutionary rescue and extinction in *Drosophila*

**DOI:** 10.64898/2026.11.17.767432

**Authors:** René S. Shahmohamadloo, Joshua J. Faber-Hammond, Amir R. Gabidulin, Catherine I. Clare, Seth M. Rudman

## Abstract

Populations declining under anthropogenic stressors, including rising temperatures and novel chemicals, may depend on rapid adaptation to persist. Yet fundamental questions remain about what promotes and constrains such evolutionary rescue. We tracked demographic, phenotypic, and genomic dynamics in a replicated outdoor evolution experiment using *Drosophila* under insecticide stress, and also manipulated initial maladaptation by pre-adapting a subset of populations. Half of populations without pre-adaptation went extinct and, unexpectedly, extinction was associated with a greater magnitude of phenotypic and parallel genomic adaptation. Rescue was driven by polygenic adaptation that was parallel across replicates and not predictable from genes previously implicated in insecticide resistance, with notable genomic convergence towards pre-adapted populations. Targeted intervention through pre-adaptation prevented extinction entirely, despite evidence of pleiotropic costs. Without intervention, evolutionary rescue remains contingent even with exceptionally rapid adaptation.

## Main Text

Environmental change and overexploitation are driving accelerated loss of biodiversity, with projections suggesting a mass extinction is already underway (*1*–*3*). Whether biodiversity will persist beyond projected rates of extinction depends, in part, on whether adaptive evolution can reduce the mismatch between organismal phenotypes and the conditions of changing environments rapidly enough to bolster population growth and enable persistence through evolutionary rescue (*4*, *5*). There is considerable optimism that evolutionary rescue could facilitate biodiversity preservation (*5*–*9*), and it is plausible that the likelihood of successful rescue could be increased through proactive evolutionary or genetic manipulations (*10, 11*). Understanding whether evolutionary rescue can be reliably predicted (*8*), and whether genetic manipulations can decrease extinction risk under rapid environmental change, is critical to proactive conservation (*1*).

One factor predicted to strongly influence the probability of evolutionary rescue in response to environmental change is the degree of initial maladaptation (*12*, *13*). Maladaptation lowers mean absolute fitness and can depress it below replacement and intensify selection, reducing the probability evolution can sufficiently shift vital rates to restore positive population growth (*14*). Interventions that increase the frequency of beneficial alleles in a new environment (e.g. assisted gene flow) can decrease the phenotype-environment mismatch in anticipation of environmental change and potentially increase the probability of rescue (*10*, *11*, *15*). Microbial evolution experiments provide empirical support for such interventions (*16*–*20*). Yet these manipulations may fail if environmental variation causes the direction and strength of selection, and therefore the fitness benefits of introduced alleles, to change through time (*21*, *22*), or if introduced alleles impose fitness costs in other environments through antagonistic pleiotropy or negative epistatic interactions (*10*, *23*–*25*). These complexities and resultant uncertainties are likely to be compounded in sexually reproducing natural populations, where environmental heterogeneity, genetic linkage, complex traits, and multifaceted phenotype-fitness relationships may make the ultimate demographic benefits of reducing initial maladaptation difficult to predict.

Understanding the demographic consequences of rapid evolution is particularly important in cases of evolutionary rescue, and such cases are powerful for understanding how genomic adaptation interacts with demography in declining populations more broadly. Adaptation that underlies rescue may occur through rapid shifts at large-effect loci or concurrent shifts at many small-effect loci, with differences in genomic architecture potentially influencing the rate, repeatability, and genomic signatures of adaptation (*13*, *26*–*28*). Genomic architecture may also shape the costs of rescue: large-effect adaptive alleles are frequently associated with antagonistic pleiotropy or fitness trade-offs (*29*, *30*), but whether costs are reduced when adaptation involves many loci of small-effect rather than few of large-effect has rarely been tested directly. Repeated sampling through time provides a resolution to assess costs of adaptation, since alleles that carry them should tend to decline in frequency once conditions change or the stressor is removed (*24*). Failed rescue, however, is rarely observed alongside successful rescue under shared conditions (*9*), leaving unclear whether populations that go extinct show distinct, and potentially diagnostic, patterns of evolution and demography from those that persist through evolutionary rescue. Temporal genomic sequencing of replicate populations undergoing both outcomes, would allow these predictions to be tested directly by linking allele frequency change to phenotypic adaptation and demography. These comparisons could reveal whether adaptation proceeds via repeatable or contingent genomic trajectories, and identify patterns of allele frequency change unique to populations that fail to persist.

The interaction between demography and the efficacy of selection could further complicate genomic adaptation in cases of evolutionary rescue. As populations decline, reduced effective population size may amplify genetic drift (*31*, *32*), lowering the efficacy of selection on moderate-effect loci and limiting the extent to which adaptation can contribute to demographic recovery (*33*). In practice, this drift-selection balance may depend on the rate of demographic decline, as well as the strength of selection and the genomic architecture of the adaptive response (*34*, *35*). Determining whether selection remains efficacious in driving molecular evolution, even as populations decline towards extinction, is key to extending theory largely developed for equilibrium conditions to the dynamics of evolutionary rescue in rapidly declining natural populations (*14*, *36*).

Here, we use a field evolution experiment with *Drosophila* populations exposed to insecticide stress to test whether demographic, evolutionary, or genomic factors predict evolutionary rescue, including whether reducing initial maladaptation is effective as a means of ‘pre-rescue’ (Fig. 1). Insecticide resistance continually threatens global food security and reduces the effectiveness of disease vector control (*37*–*39*). Resistance evolution also provides some of the clearest examples of evolutionary rescue in nature, offering tractable systems for directly observing the pace and limits of adaptation (*27*, *40*, *41*). Spinosad was used as the focal insecticide; it is widely applied to control drosophilid pests, and resistance has evolved with damaging consequences for control (*42*). We exposed replicate outdoor populations of *Drosophila melanogaster* (*41*, *43*, *44*) to sustained spinosad stress at an LC50 concentration (fig. S1) in two treatments that differed in prior selection history: (i) outbred *Drosophila* reared indoors for nine generations on standard medium, then used to establish outdoor populations reared on a spinosad-treated medium (hereafter ‘exposed’)—testing whether rescue and extinction can be predicted; and (ii) outbred *Drosophila* reared indoors for nine generations on a spinosad-treated medium, then used to establish field populations exposed to the same diet (hereafter ‘pre-adapted’)—testing whether prior selection reduces maladaptation and lowers extinction risk. A third treatment never received spinosad (hereafter ‘unexposed’) and provided a reference for demographic, phenotypic, and genomic comparisons. This design allows for direct comparison of rescue and extinction outcomes—within the exposed treatment, where populations diverged in fate despite identical founding conditions, and between treatments differing in initial maladaptation—while characterizing genomic responses through repeated genome-wide pooled sequencing in a naturally complex outdoor environment. We use these data to test whether successful and failed evolutionary rescue are predictable, and whether rescue and pre-adapted populations achieve persistence through similar mechanisms.

**Fig. 1.**
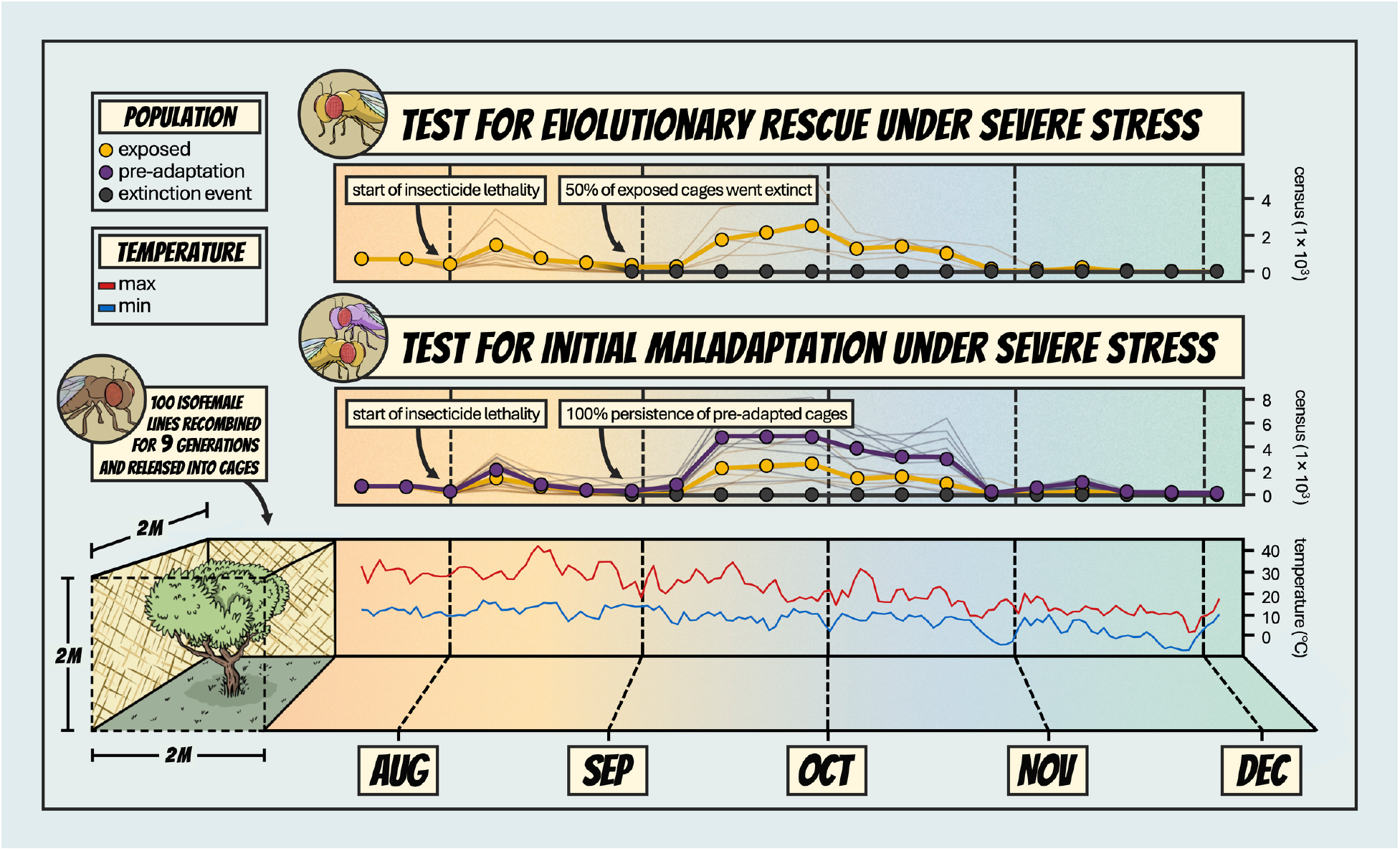
Experimental design and demographic outcomes of an evolutionary rescue experiment comparing exposed and pre-adapted populations under severe stress. All treatments originated from a single genetically diverse base population created by recombining 100 inbred lines of *Drosophila melanogaster* (Drosophila Genetic Reference Panel, (*45*)) (*44*), but differed in selection history before field exposure: founders of exposed populations evolved without the insecticide, whereas founders of ‘pre-adapted’ populations underwent selection for resistance to the insecticide in the lab. Ten outdoor mesocosms (2 × 2 × 2 m) per treatment were each founded with 2,500 flies; exposed and pre-adapted populations were fed insecticide-treated medium, and unexposed populations were fed untreated medium. Adult census size was estimated weekly using a custom-trained machine learning algorithm (*46*) across 5 months of seasonal change (thick line, treatment mean; thin lines, replicates), alongside daily minimum and maximum temperatures (blue and red lines). Exposed populations collapsed sharply in August (“extinction event”), with 50% of replicates going extinct. Phenotypic and genomic evolution were assessed on the founder population and at four subsequent time points (August to November): eggs from each replicate were reared under common garden laboratory conditions for two generations and scored for six fitness-associated phenotypes, and of 200 flies per replicate were sequenced to quantify changes in allele frequencies.

## Results

### Polygenic resistance evolves rapidly from standing genetic variation

Exposed populations evolved resistance rapidly, diverging from unexposed populations (*F*_4,56_ = 14.40, *P* = 3.71 × 10^−8^; Fig. 2A) and, over the growing season, in fecundity and starvation tolerance (*F*_4,58_ = 2.90, *P* = 0.029, Fig 2B; *F*_4,70_ = 3.27, *P* = 0.016, Fig 2C; see fig. S2 for other traits measured) (*47*, *48*). Laboratory pre-adaptation was nonetheless more effective: after nine generations of selection at the same insecticide concentration, pre-adapted populations had substantially higher resistance than exposed populations achieved in the field (*F*_4,68_ = 28.49, *P* = 6.48 × 10^−14^; Fig. 2A). The greater magnitude of resistance evolution in pre-adapted populations may reflect the simpler and more consistent selective environment of the laboratory, which can translate to stronger responses to selection (*49*, *50*). In the field, by contrast, adaptation may be constrained by multidimensional and temporally varying selection, and shifting optima combined with genetic correlations and pleiotropy constrain the multivariate response to selection (*26*, *51*–*53*).

**Fig. 2.**
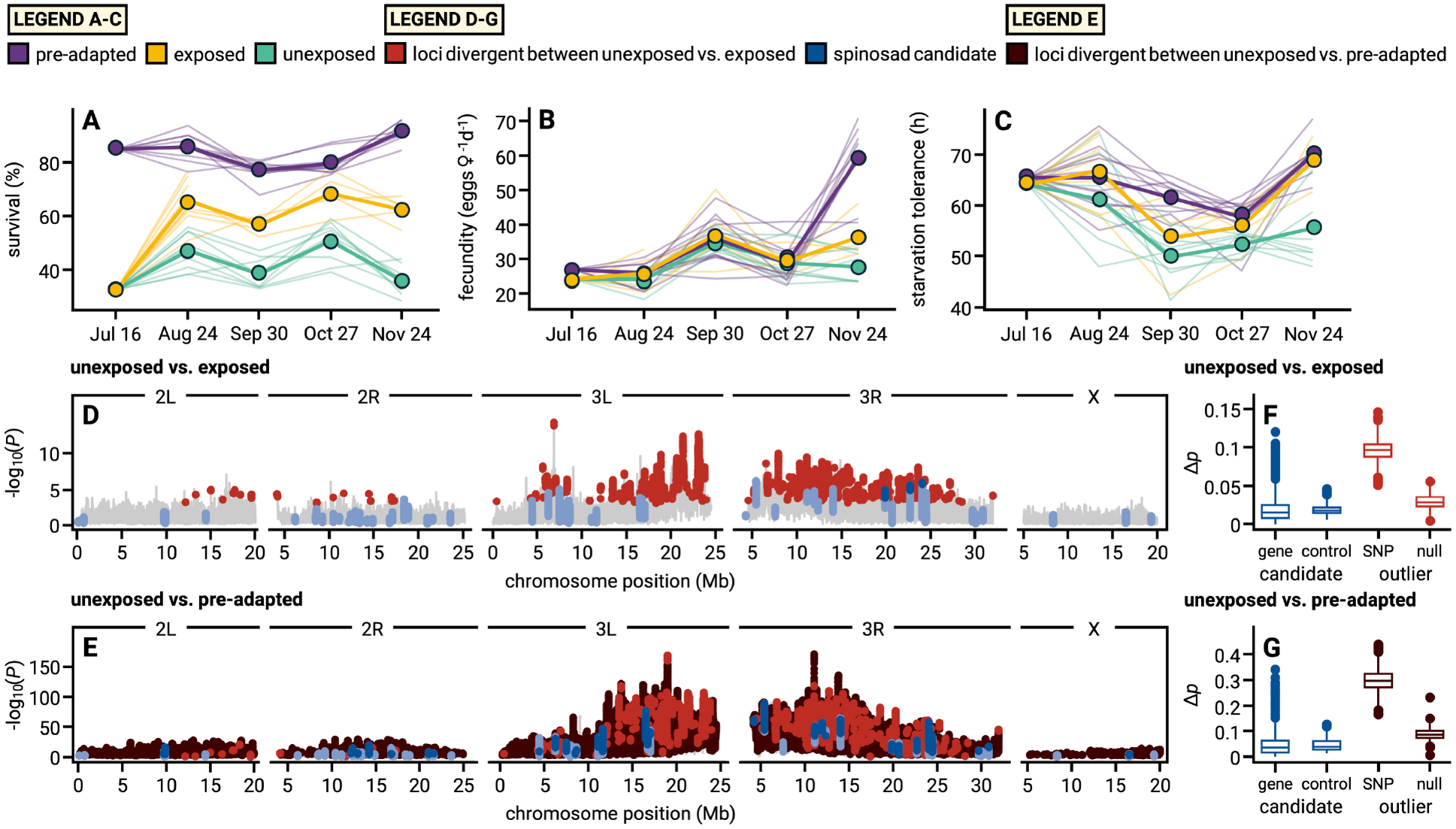
Insecticide resistance evolution is rapid, polygenic, and not driven by selection on candidate genes. (**A** to **C**) Phenotypic evolution over time. Eggs from outdoor populations were reared for two generations in a laboratory common garden to remove environmental effects. (**A**) Resistance, as percent survival from egg to adult on spinosad-treated medium. (**B**) Fecundity, as eggs per female per day. (**C**) Starvation tolerance, as time to death without food. (**D** and **E**) Allele frequency divergence between unexposed populations and (**D**) exposed or (**E**) pre-adapted populations, shown as −log_10_(*P*) from GLMs of 21-SNP sliding windows; each point is the locus at the window center. Red: loci divergent between unexposed and exposed populations (FDR < 0.01); dark blue: significant loci in previously characterized spinosad resistance genes; light blue: non-significant candidate SNPs; dark brown (**E** only), loci divergent between unexposed and pre-adapted populations (FDR < 0.01). (**F** and **G**) Absolute allele frequency differences (Δ*p*) for candidate resistance genes versus matched control genes, and for outlier SNPs from the “unexposed vs. exposed” contrast versus matched null SNPs, in (**F**) “unexposed vs. exposed” and (**G**) “unexposed vs. pre-adapted” comparisons. Overlapping distributions between candidate and matched null SNPs indicates no enrichment; outlier SNP differences show effect sizes at loci with parallel allele frequency change compared to matched sites.

The genomic architecture underlying the evolutionary divergence between exposed and unexposed populations is consistent with polygenic adaptation from standing genetic variation rather than a small number of large-effect resistance variants or mutations (*54*, *55*). Both exposed (4,189 SNPs, FDR < 0.01) and, to a greater extent, pre-adapted (134,215 SNPs, FDR < 0.01) populations exhibited genome-wide allele frequency divergence from unexposed populations (Fig. 2D-E; unsmoothed results in fig. S3). The magnitude of allele frequency change at outlier loci was substantial (Fig. 2F-G): exposed populations exhibited average allele frequency shifts of 9.8% and pre-adapted populations 25.6% at outlier loci by the end of the experiment. Despite uniform insecticide dosage across treatments, genomic signatures of selection were considerably less strong in exposed populations relative to pre-adapted populations. This could stem from the temporally variable fitness landscape (i.e., seascape; see (*49*)) in the field environment, with environmental fluctuations altering the efficacy of selection (*56*). Resistance alleles may have carried costs in unexposed populations; alleles at sites with greatest initial divergence between pre-adapted and unexposed populations showed subsequent temporal evolution away from the putative resistance alleles in unexposed populations (hypergeometric overlap test, *P* = 9.8 × 10^−18^), indicating selection against resistance variants in the absence of the insecticide stressor (see *Materials and Methods*, table S1).

Although resistance evolved rapidly and repeatedly, allele frequency divergence from unexposed populations at loci in genes previously characterized as underlying spinosad resistance (*57*–*59*) was no greater than at matched control loci in either exposed (Welch’s *t*-test, *P* = 0.78; Fig. 2F; fig. S3; fig. S4; table S2) or pre-adapted (Welch’s *t*-test, *P* > 0.99; Fig. 2G; table S2) populations. Instead, the genomic architecture of divergence suggests a polygenic basis of adaptation with substantial genetic draft (fig. S3), which itself complicates tests for enrichment. The lack of elevated divergence at candidate genes amongst the highly polygenic response in exposed populations, including at the top outliers within clusters of divergent SNPs to account for draft, suggests that previously characterized spinosad resistance mechanisms spanning target-site, detoxification, transport, and cuticular pathways need not contribute to the genetic architecture of rapid field-evolved resistance (table S2). This fits with emerging data on the limited predictive power of genetic mapping for forecasting the genomic architecture of polygenic adaptation (*60*), but is surprising given the noted repeatability of molecular mechanisms conferring resistance across taxa (*61*–*63*).

### Extinction occurs despite strong adaptive responses

Insecticide exposure immediately and significantly slowed demographic expansion relative to unexposed populations (Welch’s *t*-test, *t*_16.0_ = 2.25, *P* = 0.039). The magnitude of demographic suppression was amplified by a naturally occurring heatwave that drove demographic declines across all treatments (fig. S5): peak temperatures during August 14-16, 2023 averaged 40.57 ± 1.48 °C, which was 3.48 SDs above the 2013-2022 historical baseline (27.72 ± 3.69 °C)—with a maximum of 42.2 °C on August 14, 2023. These temperatures exceeded the short-term critical thermal maximum of adult *D. melanogaster* (∼39–41°C; (*64*)) and the chronic developmental thermal maximum (∼29°C; (*65*, *66*)), causing substantial mortality of larvae and adults across all treatments and reducing census sizes by a median decline of 86% (fig. S5). Recovery to pre-heatwave abundances, however, varied considerably across treatments despite comparable declines: unexposed populations recovered within 5 weeks, and all pre-adapted populations recovered within 3 weeks despite continued insecticide exposure. In contrast, 50% of the exposed populations failed to recover and went extinct (hereafter ‘extinct’). The remaining 50% underwent evolutionary rescue (hereafter ‘rescue’), recovering within 4 weeks, as demonstrated by the U-shaped demographic trajectory (Fig. 3A) and concurrent phenotypic resistance evolution (Fig. 2A). Multistressor environments shaped both population dynamics and evolutionary trajectories (*67*–*69*), with the combination of insecticide and heat stress causing high rates of extinction in the absence of pre-adaptation.

**Fig. 3.**
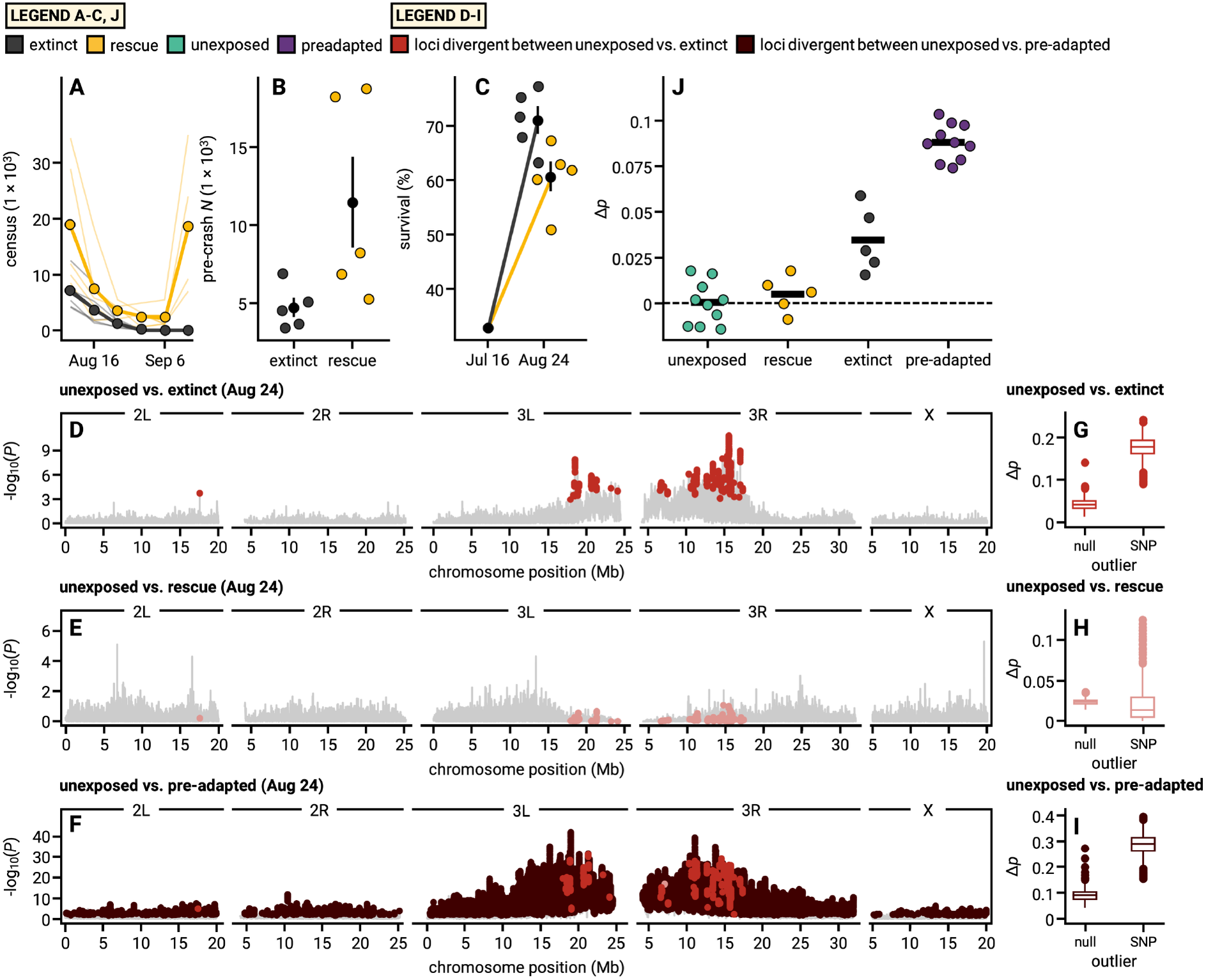
Demographic, phenotypic, and genomic factors associated with failed and successful evolutionary rescue. (**A**) Adult census estimates spanning the period of collapse and subsequent recovery or extinction; lines represent individual mesocosms. (**B**) Pre-crash adult abundance by fate. (**C**) Magnitude of phenotypic resistance evolution by fate. (B to C) Points represent individual mesocosms; symbols with vertical lines denote group means ± SE. (**D** to **F**) Allele frequency divergence in August between unexposed populations and extinct (**D**), rescue (**E**), or pre-adapted (**F**) populations, shown as −log_10_(*P*) from GLMs in 21-SNP sliding windows. Red, loci divergent between unexposed and extinct populations (FDR < 0.05 in **D**), shown in all three panels for comparison but denoted as light red in (**E**), where they are not significant. Brown (**F**), loci divergent between unexposed and pre-adapted populations (FDR < 0.05). (**G** to **I**) Absolute allele frequency differences (Δ*p*) of GLM outlier SNPs in August compared with matched background SNPs for the unexposed versus extinct (**G**), rescue (**H**), and pre-adapted (**I**) contrasts. Extinct-contrast outliers are displayed in (**G**) and (**H**), as no loci were significant in the rescue contrast. (**J**) Mean allele frequency shift (Δ*p*) of each population in August, relative to the August unexposed mean, in the direction of pre-adapted allelic change at SNPs differentiating pre-adapted and unexposed populations in November (FDR < 0.01; 428,205 SNPs). Points are individual mesocosm populations; bars are group means.

Next, we asked what facets of the demographic and evolutionary trajectories distinguished extinct and rescue populations. Populations that went extinct had lower pre-crash adult abundances than those that underwent rescue despite identical founding and exposure conditions (mean *N*: 4,742 in extinct vs. 11,452 in rescue populations; linear model, *F*_1,8_ = 7.46, *P* = 0.026, R^2^ = 0.48; Fig. 3B). Extinct populations exhibited higher rates of phenotypic adaptation than rescue populations, including greater resistance (Fig. 3C; 71% vs. 61%; Wilcoxon rank-sum test, *W* = 24, *P* = 0.016) and greater starvation tolerance (fig. S6; 71.1 vs. 62.9 h; Wilcoxon rank-sum test, *W* = 23, *P* = 0.032). Moreover, extinct populations differed in the direction of overall phenotypic evolution (PC2: β = -1.77, *P* = 0.024, R^2^ = 0.49; fig. S7).

Fast rates of adaptation are often considered a key factor in evolutionary rescue outcomes (*5*, *72*), but in extinct populations we observed greater phenotypic evolution. That extinction occurred despite greater phenotypic adaptation is consistent with a demographic limit that adaptation could not overcome (*70*, *71*). The lower census size of these populations may have magnified both the rate of evolution, as genetic storage effects would be muted, and the cost of selection as populations approached extinction (*73*, *74*). These results indicate that rates of adaptive evolution may be unreliable predictors of evolutionary rescue (*75*, *76*), as the genetic and demographic costs of strong selection, including drift load, trade-offs, and storage effects, may decouple adaptive rates from demographic outcomes (*73*, *77*, *78*).

Genomic divergence was faster, of greater magnitude, and more parallel among populations that went extinct. By August, extinct populations exhibited significant parallel allele frequency divergence from the unexposed populations (619 significant SNPs, FDR < 0.05), indicative of adaptive divergence associated with insecticide resistance (Fig. 3D; unsmoothed results in fig. S8). Divergent SNPs were grouped into clusters, defined as genomic regions in which divergence exceeded a permuted null expectation (adapted from (*43*); see *Materials and Methods*), providing an estimate of the number of distinct genomic regions that exhibited patterns of parallel genomic evolution indicative of natural selection. The strength of selection, pace of adaptation, and movement of founding haplotypes produced substantial genomic draft; the 619 SNPs were distributed across 15 clusters, concentrated on chromosome arms 3L and 3R (fig. S9A). The magnitude of allele frequency divergence between extinct and unexposed populations at outlier loci was well beyond divergence at matched background loci (mean Δ*p* = 0.18 vs. 0.04, *P* < 2.2 × 10^−16^; Fig. 3G), and allele frequency changes at top outlier loci within each divergence cluster were largely parallel across extinct populations (one-tailed *t*-tests, median *t* = 7.16, median *P* = 8.05 × 10^−11^; table S3), implicating directional selection as a driving force in the divergence from unexposed populations. In contrast, no SNPs in rescue populations passed the significance threshold for divergence by August (Fig. 3E), and only three clusters of divergence were detected (fig. S9B). At the loci that distinguished extinct populations, rescue populations showed no elevated divergence relative to background (mean Δ*p* = 0.02 vs. 0.02, *P* = 1.0; Fig. 3H), and allele frequency change at the few clusters detected in rescue populations was not detectably parallel (one-tailed *t*-tests, median *t* = 0.91, median *P* = 0.20; table S4), mirroring their weaker phenotypic responses to selection. Founder effects could, in principle, generate genomic differences between extinct and rescue populations. Two lines of evidence indicate selection is the more likely mechanism: allele frequency divergence was concordant across independent populations, and patterns of genomic divergence between extinct and rescue populations exceeded those observed among unexposed replicates (fig. S10). Parallel allele frequency change during demographic decline to extinction, despite the increasing potential for drift (*79*, *80*), likely reflects both the strength of selection and a rapid demographic collapse which was too fast for genetic diversity loss via drift (i.e. time lag, (*81*)) (table S5).

If resistance evolves through repeatable genomic mechanisms, populations independently responding strongly to insecticide selection may show patterns of convergence. By August, pre-adapted populations had diverged strongly from unexposed populations, with outlier loci concentrated on chromosome arms 3L and 3R (Fig. 3F) but clusters of divergence spanning all chromosome arms (fig. S9C), and elevated divergence relative to matched background loci (Fig. 3I), providing a reference against which to assess convergence. To test for convergence, we asked how far each exposed population had shifted toward the pre-adapted state by August, measured at SNPs that had diverged between pre-adapted and unexposed populations by the end of the experiment (GLM FDR < 0.01; 428,205 SNPs; see *Materials and Methods*). Just prior to extinction, extinct populations had converged more strongly with pre-adapted populations than rescue populations (mean Δ*p* = 0.035 vs. 0.005; exact permutation test, *P* = 0.016; Fig. 3J), compared with a mean shift of 0.088 in pre-adapted populations themselves. In contrast, convergence observed in rescue populations fell within what was observed among unexposed populations. A genome-wide analysis across all loci corroborated this pattern: Spearman’s rank correlations in the direction and significance of allele frequency change were greater between extinct and pre-adapted populations (ρ = 0.56) than between rescue and pre-adapted populations (ρ = 0.14), each relative to unexposed populations (fig. S11). Together, these results indicate that populations that went extinct exhibited genomic convergence toward pre-adapted populations prior to extinction, but that a shared genomic response to the insecticide, and the larger magnitude of phenotypic resistance evolution, were insufficient for persistence. The pace of phenotypic and genomic adaptation, combined with lower pre-crash adult abundance, suggests that a reduced storage effect and lower demographic buffering may explain the divergence in fates.

### Pre-adaptation is costly but effective, and shares a genomic basis with de novo rescue

Pre-adaptation was uniformly successful at preventing extinction (fig. S5), even amid the heatwave when multistressor interactions were most consequential (Fig. 4A), whereas only half of exposed populations persisted through rescue without prior laboratory selection (hereafter ‘*de novo* rescue’, from standing genetic variation). Vector angle analysis of multivariate phenotypic evolution (*82*) showed that pre-adapted populations evolved along closely aligned trajectories during the outdoor experiment (mean angle = 22°, significantly < 90°; *t*_9_ = -45.10, *P* = 6.48 × 10^−12^), rescue populations also evolved in parallel but showed greater across-replicate phenotypic dispersion (mean angle = 57°, significantly < 90°; *t*_4_ = -4.14, *P* = 0.014; Fig. 4B). Greater parallelism among pre-adapted populations (Welch’s *t*-test, *t*_4.3_ = -4.30, *P* = 0.011) is unlikely to reflect their slightly lower genetic diversity (table S5), and more likely reflects how prior evolution altered genetic covariances among traits or constrained subsequent evolutionary trajectories through epistatic interactions (*83*, *84*).

**Fig. 4.**
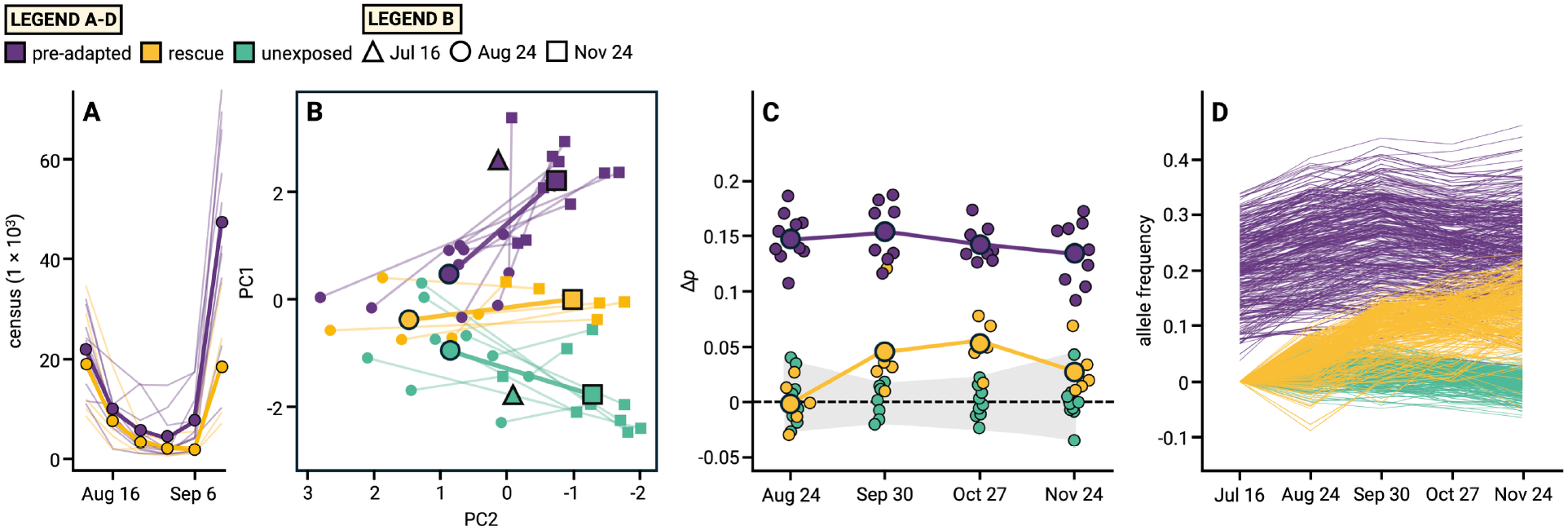
Demographic, phenotypic, and genomic convergence between pre-adapted and rescue populations suggests a shared basis for evolutionary rescue. **(A)** Census size for pre-adapted and rescue populations during the critical extinction period. Bold lines show treatment means; thin lines show replicates. **(B)** Multivariate phenotypic trajectories across treatments and time. Bold vectors connect population-level means from August 24 (●) to November 24 (▪); founder populations (July 16, ▴) are additionally plotted. Small symbols and thin lines denote individual replicate populations. **(C)** Mean allele frequency shift (Δ*p*) in the direction of pre-adapted allelic change, relative to concurrent unexposed populations, at SNPs differentiating the pre-adapted and unexposed founder populations on July 16 (222,983 SNPs). Small symbols are individual mesocosm populations and large symbols are group means, with bold lines connecting rescue and pre-adapted means. The shaded band spans the values of unexposed populations, each scored against the mean of the other unexposed populations. **(D)** Allele frequency trajectories across the full time series (July 16 to November 24) for the 441 SNPs showing divergence from unexposed populations in both rescue and pre-adapted populations by November 24, expressed relative to the July 16 unexposed population allele frequencies (see *Materials and Methods* for full filtering criteria and fig. S12). Each line shows the allele frequency trajectory of one SNP within one treatment.

By November, rescue populations had also evolved in parallel at the genomic level: at top divergent sites, allele frequencies shifted in the same direction in all five replicates relative to random matched loci (one-tailed *t*-tests; median *t* = 4.25, median *P* = 1.34 × 10^−5^; table S6), indicating that continued insecticide selection and temporal adaptation produced increasingly parallel genomic adaptation. Genetic interventions to reduce maladaptation have been recognized as promising to promote rescue (*10*, *11*), but whether these interventions occur via the same genetic mechanisms as *de novo* rescue is largely unknown. Shared molecular constraints and standing genetic variants could generate parallel or convergent responses across treatments, whereas epistasis, genotype-by-environment interactions, pleiotropy, and a highly polygenic basis for fitness variation could all produce largely treatment-specific patterns of genomic evolution. We tested whether pre-adaptation and *de novo* rescue occurred via selection on similar genomic variants by examining whether allele frequency divergence in pre-adapted populations occurred at the same loci—and in the same direction—as selection on rescue populations in field conditions. Rescue populations underwent convergent genomic evolution toward alleles favored during laboratory pre-adaptation. Along an axis of SNPs differentiating the pre-adapted and unexposed founder populations before field exposure (222,983 SNPs; see *Materials and Methods*), rescue populations were indistinguishable from unexposed populations in August (Δp = −0.001; exact permutation test, *P* = 0.92) but increasingly showed a pattern of convergence towards pre-adapted populations in September, October, and November (Δ*p* = 0.045, 0.054, and 0.029; *P* = 0.003, 0.001, and 0.030; Fig. 4C). By October, rescue populations had converged to a similar extent as extinct populations in August (Δ*p* = 0.055; relative to unexposed populations, *P* = 0.003), approximately 38% of the total divergence between unexposed and pre-adapted populations.

Rescue populations diverged from unexposed populations at 4,429 SNPs (found across 157 clusters across all chromosomal arms; fig. S9D) by November, of which 2,456 also diverged in parallel in pre-adapted populations. Applying more stringent filtering criteria, we identified 441 convergent outlier SNPs in both pre-adapted and rescue populations in November (Fig. 4D), almost entirely located on chromosome arms 3L and 3R (see *Materials and Methods*; fig. S12, table S7). Of these, 81 had high predicted functional impacts based on a machine learning model trained on *Drosophila* variants (table S7) (*85*). The overall pattern of adaptation and convergence we observed in these loci is consistent with resistance arising through polygenic mechanisms across detoxification and redox pathways (table S8) rather than single large-effect alleles contributing to target site resistance (*62*, *86*). The design of the experiment and sampling here gives reasonable power to detect putative resistance variants that contribute in natural populations, which is challenging in field contexts (*58*).

Loci identified as initially most divergent between pre-adapted and unexposed populations subsequently declined in divergence magnitude over the following months, with pre-adapted populations shifting back toward unexposed allele frequencies. These initial outliers were substantially enriched among loci showing convergence in pre-adapted populations toward unexposed populations between September and November (59.2% of 8,761 loci; hypergeometric overlap *P* = 1.49 × 10^−242^), but not among loci diverging in pre-adapted populations (25.3% of 3,586 loci; hypergeometric overlap *P* = 1) (see *Materials and Methods*, table S9). We extended this test for convergence across exposed treatments: loci distinguishing pre-adapted or extinct populations from unexposed populations in August shifted predominantly pre-adapted populations in the direction of the allele that was more common in unexposed populations in the vast majority of time intervals (fig. S13A–B). In contrast, loci distinguishing rescue populations, in either August or November, showed no such reversal: in pre-adapted populations, the majority of these loci shifted in the direction of rescue allelic change in all time intervals, consistent with monotonic divergence (fig. S13C–D).

Patterns of reversal occurring despite continued exposure to the insecticide suggest that some alleles favored in the laboratory environment carried costs that only became apparent under the multifarious and fluctuating selective environment in the field. The costs of pre-adapted alleles were more visible than any associated with *de novo* rescue. These patterns may reflect genotype-by-environment interactions between lab and field selection, including pleiotropic costs at loci originally under positive selection in the laboratory—a concern for using assisted evolution to rescue wild populations (*9*, *11*, *87*). Despite this genomic signature of cost, the phenotypic and demographic benefit of pre-adaptation was evident throughout our experiment, and represents a promising avenue for reducing extinction risk under environmental change (*15*).

## Discussion

The extinction we observed did not reflect a slower rate of adaptation, but a mismatch between the tempo of evolutionary change and the co-occurring demographic decline (*8*, *12*, *32*, *73*, *88*). Strong selection in small, declining populations simultaneously increases the probability of negative hitchhiking (*89*), negative epistatic interactions (*90*), and fitness trade-offs (*91*). These processes collectively erode demographic recovery and may contribute to Allee effect dynamics in which per-capita growth rates decline at low density (*92*). In this experiment, differences in the magnitude of genetic storage effects (*93*), spurred by initial demographic stochasticity (*14*, *94*–*96*), may explain why the response to selection was stronger in populations that ultimately went extinct. The insecticide’s mode of action may have also contributed to this pattern. Spinosad causes mortality during larval development; in stage-structured populations, surviving adults continue to reproduce while selection acts on their offspring. This could allow populations with higher pre-crash adult abundances to sustain a greater demographic cost of selection, but exhibit a slower rate of evolution, and still ultimately persist. Whereas prior work has emphasized that the pace of adaptation is often too slow to permit evolutionary rescue across many species (*97*), our results show a complementary limitation: even exceptionally rapid, parallel adaptation did not reliably translate to an increase in the probability of evolutionary rescue. Differences in population size, rather than rates of adaptive evolution, are among the most likely explanations for rescue versus extinction, suggesting caution when projecting conservation outcomes based primarily on rates of evolution.

Notably, strong selection here drove rapid adaptation, but at demographic costs that reduced the likelihood of rescue, consistent with recent outdoor evolution experiments that identified eco-evolutionary tipping points where extreme selection overwhelms adaptive potential (*98*). However, the populations that went extinct in the present work showed no sign of lost adaptive potential: they evolved greater resistance than rescue populations, their allele frequency change was highly parallel across replicates, and their genetic diversity did not differ from that of rescue populations. Instead, persistence depended on whether populations could sustain the demographic cost of strong selection. Incorporating the full complexity of selection—with stage structuring, population dynamics, and density dependence—into projections of future rates of adaptation is an underappreciated challenge for evolutionary predictions (*75*, *99*–*101*), particularly when forecasting demography and persistence. Progress will require studies that measure demographic rates and adaptive capacity in the same populations, since projections of persistence depend on both. Parameterizing adaptive seascape models offers a potential path forward (*49*).

Reducing initial maladaptation through pre-adaptation (or ‘pre-rescue’) proved highly effective at preventing extinction. The magnitude of the benefit exceeded what prior experiments (*16*) would lead one to anticipate: all pre-adapted populations persisted and evolved along tightly parallel phenotypic trajectories in a dynamic, multistressor environment. Pre-adaptation partly shared a genomic basis with rescue in the field, as rescue populations later shifted toward alleles favored during pre-adaptation, yet many alleles that increased in frequency under laboratory selection were subsequently selected against in the field—a pattern consistent with the prediction that trade-offs are more likely to evolve under directional than fluctuating selection (*24*). The costs implied by this reversal evidently did not offset the advantage of reducing initial maladaptation via pre-adaptation.

Here, pre-adaptation was wholly effective at preventing extinction despite the insecticide being imposed within a complex, ecologically realistic setting. This is consistent with evidence that introducing pre-adapted individuals, whether through assisted gene flow (*102*–*104*) or reintroductions (*105*, *106*), can improve tolerance and, in some cases, demographic performance under rapid environmental change. Whether pre-adaptation to stressors with more diffuse or pleiotropic targets of selection than insecticides would prove similarly effective at spurring evolutionary rescue is worthy of future investigation. The broader challenge, therefore, lies in translating evolutionary rescue from a conceptual framework into a practical conservation tool—one our results suggest should prioritize proactively reducing maladaptation and closely tracking demography, given that rapid adaptation alone may be insufficient to prevent extinction.

Materials and methods are available in the supplementary materials.

## Acknowledgments

We thank D. Schluter and S. Otto for feedback and discussions on pre-adaptation; M. Osmond for feedback and discussions on evolutionary rescue theory; K. Lotterhos for discussion on gene flow; and J. Mallet for discussion of carrying capacity and the links between demography and Darwinian evolution. We are grateful to the late R. Gomulkiewicz for discussions on doom and persistence that motivated this study. We thank members of the Rudman, Osmond, and Lotterhos labs for feedback on the research and manuscript. We are grateful to E. Andrews, B. Campagnari, and E. Prileson for assistance with experimental maintenance and data collection, and to T. Gabidulin for creating the artwork illustrated in Fig. 1. 

## Author contributions

Conceptualization: SMR

Methodology: RSS, SMR

Formal analysis: RSS, JJFH

Investigation: RSS, ARG, CIC, SMR

Writing – original draft: RSS, SMR

Writing – review & editing: RSS, JJFH, ARG, CIC, SMR

## Data, code, and materials availability

All code necessary to replicate the analyses presented in this manuscript is available at https://github.com/jfaberha/drosophila_evolutionary_rescue. Pooled sequencing reads generated for this study are available at the NCBI under BioProject accession PRJNA1468879. Upon manuscript acceptance, demographic, phenotypic, environmental, and allele frequency data, together with the input files required to run the genomic analyses, will be archived in Dryad.

## Supplementary Materials

### Materials and Methods

#### 1.1 Founding populations

The founding population for all experimental treatments was derived from a recombined population created by crossing 100 inbred lines from the *Drosophila melanogaster* Genetic Reference Panel (DGRP) (*45*), following established protocols (*43*, *44*, *47*). Three experimental treatments were established from this base population: (i) unexposed populations were reared indoors for nine generations on standard food media without insecticide exposure, then released into outdoor mesocosms where they continued on standard food media throughout the experiment; (ii) exposed populations were reared indoors for nine generations on standard food media, then released into outdoor mesocosms where they were exposed to spinosad-amended food media at an LC50 concentration (see *1.2 Media preparation*) for the first time; and (iii) pre-adapted populations were reared indoors for nine generations on spinosad-amended food media at an LC50 concentration, then released into outdoor mesocosms where they continued on spinosad-amended food media at the same concentration.

#### 1.2 Media preparation

Standard fly food media (modified Bloomington recipe - per liter: 68 g cornmeal, 16 g yeast, 9 g soy flour, 65 mL molasses, 5 g agar, and 2 g methylparaben (Tegosept) dissolved in ethanol) was prepared for unexposed mesocosms and served as the base recipe for all treatments. Spinosad (Entrust® SC, 22.5% active ingredient; Corteva Agriscience, Indianapolis, IN, United States) was diluted in sterile distilled water and incorporated directly into fly food media during preparation. To ensure homogeneous distribution, the insecticide was added after the media cooled below 60°C and immediately before dispensing into vials. To quantify toxicity, concentration-response assays were conducted by rearing flies from egg to adult on media containing a range of spinosad concentrations (0, 0.0188, 0.0375, 0.075, 0.1125, 0.15, and 0.225 µg L⁻¹). For each concentration, 30 eggs were placed per vial (*n* = 6 vials per concentration) and resistance, measured as percent survival from egg to adulthood, was recorded. The LC50 was estimated using logistic regression in R (fig. S1), yielding a treatment concentration of 0.0375 µg L⁻¹ used for both laboratory pre-adaptation and field exposure of the exposed and pre-adapted treatments. Food for unexposed mesocosms was always prepared first using separate utensils and containers, and insecticide-treated food was prepared in a dedicated area with specially labeled equipment stored separately from unexposed food.

#### 1.3 Outdoor experimental design

The replicated outdoor mesocosm experiment was conducted in Vancouver, WA, United States, from July to November 2023 using 2 × 2 × 2 m mesh enclosures (BioQuip PO 1406C) containing a non-fruiting dwarf apple tree and vegetative ground cover to provide shade and a semi-natural ecological environment (*43*, *44*, *47*). On July 16, 2023, each mesocosm was seeded with 2,500 F9 adults (ninth-generation descendants of the base population) of a single age cohort. The experiment included ten replicate populations per treatment (30 total), with treatments randomly assigned to mesocosms. 350 mL of *Drosophila* media (modified Bloomington recipe) via food pans was provided three times each week as the sole food source and oviposition substrate. Food was sheltered from direct rain and sunlight. After two days of egg-laying, each pan was covered with a mesh lid to prevent further oviposition and monitored daily until adult eclosion, at which point emerging flies were released into the mesocosm (*43*, *47*, *53*). Mesh enclosures were inspected weekly for integrity to prevent migration between mesocosms.

#### 1.4 Phenotypic assays

To assess the evolutionary response to insecticide exposure, approximately 750 eggs were collected from each replicate mesocosm at four time points (August 24, September 30, October 27, November 24, 2023) and reared indoors on standard medium under common garden conditions (25°C, 12 h light : 12 h dark photoperiod) for two generations before assaying, maintaining population sizes of approximately 750 individuals per generation. Offspring of the second common garden generation were then assayed on either standard medium or spinosad-amended medium at the LC50 concentration; results from the spinosad assays are presented in the main text and fig. S2, with control medium results provided in the supplementary data files. Common garden rearing standardizes environmental conditions, removing phenotypic plasticity induced by outdoor environmental variation and maternal effects to allow estimation of genetic contributions to phenotypic evolution (*107*).

Six phenotypes were assayed per replicate population, with measurements conducted following established protocols (*43*, *44*, *46*). Resistance was quantified as the percent survival from egg to adulthood (30 eggs per vial). Development rate was calculated as the inverse of development time from egg to adult. Fecundity was measured as eggs per female per day, averaged across three consecutive days of egg laying by five females. Starvation tolerance was measured as the time to death for 10 males per vial on media containing 1% agar in water, with deaths recorded three times daily (three replicate vials per population). Heat tolerance was measured as the time to death for 15 females per enclosed petri dish in a water bath at 53°C (adapted from (*108*)). Adult dry mass was measured as the mean weight of three pools of five females dried at 55°C for 24 hours.

#### 1.5 Population census dynamics and extinction

Census size was estimated weekly in each replicate mesocosm by photographing eight 0.3 m² transects on the ceiling of each enclosure near sunset, when flies congregated on the upper portions of enclosures during the evening activity peak. Flies in each photograph were counted using a custom-trained machine learning algorithm (*46*). Total adult census estimates were obtained by summing counts across all transect photographs and multiplying by 35 to account for the surface area sampled relative to all cage substrates (*43*, *46*). Census counts continued through the end of the season or until populations reached extinction, defined as no adults detected in any transect photograph on consecutive census dates with no evidence of recruitment.

A heatwave affecting all treatments was identified based on daily maximum air temperature exceeding the 2013−2022 historical baseline for the study site by more than 3 SDs (see *Results* for full temperature statistics; daily temperature records are available in the Dryad repository). Pre-crash abundance (*N*) was quantified as the mean census abundance across the two sampling dates immediately preceding collapse (August 1 and August 9).

#### 1.6 DNA extraction and sequencing

Founder samples were collected on July 16, 2023, prior to release into mesocosms. At four subsequent time points (August 24, September 30, October 27, November 24, 2023), 200 adult flies were collected from each replicate mesocosm and flash frozen at −80°C until further processing. Individuals from each replicate mesocosm were pooled and DNA was extracted using the Monarch® Spin gDNA Extraction Kit (New England Biolabs, Ipswich, MA, United States) optimized for our sample input. Libraries were prepared using a Nextera library preparation kit (Illumina), normalized, and sequenced across two lanes of Illumina NovaSeq S4 PE150.

#### 1.7 Genomic sequencing and allele frequency estimation

Raw FASTQ files were processed using *grenepipe* to automate variant calling (*109*). Within *grenepipe*, reads were quality filtered and adapter trimmed using *fastp* (*110*), mapped to the *D. melanogaster* genome (BDGP6.46) using *bwa-mem2* (*111*), and deduplicated with Picard Tools (*112*). Variant calling and filtering were performed with *BCFtools* (*113*). Haplotype-derived allele frequencies (HAFs) were then calculated via local inference with HAF-pipe (*114*) using haplotype information from the *Drosophila melanogaster* Genetic Reference Panel 2 (DGRP2) founder lines (*45*). The resulting HAF-pipe allele frequency table was used for all downstream genomic analyses.

HAF-pipe greatly increases confidence in allele frequency calls (*115*). HAF-pipe-imputed frequencies showed stronger concordance with DGRP2 founder frequencies (Pearson’s R = 0.90) than frequencies derived from raw read counts with *BCFtools* (R = 0.86). This difference was further evident in density plots, where HAF-pipe estimates more closely followed the linear relationship observed for founder frequencies (fig. S14).

Preliminary analysis revealed that labels for six consecutive October 27 DNA samples had been transposed during processing. Corrected assignments were consistent with sample clustering by treatment via principal component and hierarchical clustering analyses of allele frequencies.

Functional annotation of detected variants was performed using Variant Effect Predictor (VEP), implemented within *grenepipe*. Sequencing quality metrics and mapping statistics were summarized using *multiqc* (*116*). Details regarding *grenepipe* parameter settings are provided in the configuration file available at https://github.com/jfaberha/drosophila_evolutionary_rescue.

#### 1.8 Data analysis

##### 1.8.1 Phenotypic evolution

To test for the effects of insecticide exposure on the evolution of fitness-related traits, we fitted linear mixed-effects models for each phenotype using the *lme4* package (*117*). Models included unexposed, exposed, and pre-adapted replicate mesocosms, sampled at up to four time points across the growing season, with treatment, sampling time point, and their interaction included as fixed effects, and mesocosm included as a random effect to account for repeated measures within experimental replicates. Significance was assessed using Type III ANOVA with Satterthwaite’s method for denominator degrees of freedom via the *lmerTest* package (*118*). Pairwise contrasts between treatments at each time point were performed using the *emmeans* package (*119*), with Benjamini-Hochberg correction applied for multiple comparisons (*120*).

##### 1.8.2 Early phenotypic divergence among exposed populations

To test for phenotypic divergence between exposed populations that were later rescued versus those that went extinct, we compared population-level mean phenotypes of the ten exposed populations measured on August 24, 2023, six days before extinction on August 30, 2023. For each trait, a linear model with persistence status (i.e., rescue vs. extinct) as a fixed effect was fitted to population means. Multivariate phenotypic divergence was assessed using principal component analysis (PCA) on all six traits (resistance, development rate, fecundity, starvation tolerance, heat tolerance, and adult dry mass). Prior to PCA, each trait was centered and scaled to unit variance to account for differences in measurement scale across traits. A PERMANOVA found no overall difference between persistence classes (*F*_1,8_ = 1.93, R^2^ = 0.19, *P* = 0.113, 9,999 permutations). Because PC2 was the axis that separated extinct from rescue populations, we tested for differences in PC2 scores between persistence classes using a linear model, reporting the coefficient (β) and R^2^ for that model.

##### 1.8.3 Multivariate phenotypic trajectories

A second PCA was used to quantify multivariate phenotypic evolution across the full experimental period. Traits were centered and scaled to unit variance (*z*-score standardized) prior to analysis, and PCA axes were constructed from the first (August 24) and last (November 24) field time points. Founder phenotypes (July 16) were subsequently projected into this PCA space using the fitted loadings. Evolutionary trajectories for each mesocosm were defined as PCA score vectors between August 24 and November 24. The magnitude of vectors was calculated as Euclidean distance, with direction and parallelism quantified using the *multivarvector* package (*82*). Under random orientation, the expected mean angle between vectors is 90°; angles significantly below this indicate more parallelism than would be expected by chance (*82*). Mean angles among trajectories were therefore compared to a null expectation of 90° using one-sample *t*-tests, and differences in parallelism between treatments were assessed using two-sample *t*-tests.

##### 1.8.4 Modeling allele frequency divergence

Generalized linear models (GLMs) were implemented in R using HAF-pipe-imputed allele frequency tables to identify genomic divergence over time and across treatments. Prior to analysis, SNPs were filtered to a minimum allele frequency variance >0.001 across samples. GLMs were fit for a defined set of treatment contrasts, applied either across the full time series or within individual time points depending on the biological question addressed. Models included treatment as a fixed effect and, where appropriate, time point and treatment × time point interaction terms, with a quasi-Poisson error distribution. Scripts used for these analyses are available at: <u>github.com/jfaberha/drosophila_evolutionary_rescue/</u>.

For visualization and most downstream functional analyses, both raw GLM *P*-values and false discovery rate (FDR) corrected *P*-values were summarized using a rolling-window average across 21 consecutive SNPs using slide_mean in the *slider* R package (*121*), with mean values assigned to center SNPs. This window-based smoothing approach was done for noise-reduction between highly divergent contrasts, particularly in regions that are not selective targets but rather appeared divergent due to linkage to true spinosad-adaptive loci. SNPs with FDR values less than 0.01 were designated significant outliers in contrasts involving samples from all time points, and below 0.05 in contrasts within a single time point. Two separate thresholds were used due to differences in sample sizes and power to detect parallel allele frequency changes, in addition to the observed magnitude of differentiation. While smoothed *P*-values and FDR values revealed strong biological signals in the majority of focal GLM contrasts, several contrasts showed surprisingly little signal of selection, therefore we additionally examined non-window-based (per-SNP) *P*-values and FDR values to assess whether minor signals of selection had been masked.

##### 1.8.5 Genetic diversity estimates

Genetic diversity was estimated using Watterson’s θ per sample and per chromosome, to assess variation associated with demographic change or bottlenecks resulting from strong selection. Estimates were generated using the *diversity* tool in *grenedalf* (*122*) with HAF-pipe-imputed allele frequency tables as input. Because Watterson’s θ requires sequencing depth and HAF-pipe-imputed allele frequencies do not correspond to raw counts, effective coverage was estimated for each sample using the HAF-pipe effective coverage calculator (https://ec-calculator.shinyapps.io/shinyapp/). To evaluate sensitivity to this parameter, diversity estimates were calculated at both the lower (500×) and upper (3,500×) bounds of estimated effective coverage, and results were examined for systematic differences across depth assumptions (table S5).

Diversity estimates from August samples were analyzed with quasi-Poisson GLMs (θ × 10^5^, rounded) with treatment as a fixed effect, fitted separately for each chromosome and genome-wide, and pairwise contrasts among treatments were tested using the *emmeans* package (*119*).

Resulting diversity estimates were analyzed using GLMs with treatment and chromosome as fixed effects.

##### 1.8.6 Functional enrichment tests

To assess whether loci associated with spinosad resistance were overrepresented among genomic signals of adaptation, we curated a set of 69 *a priori* candidate genes previously implicated in spinosad response through knockout and RNAi screens, differential expression under insecticide exposure, heterologous expression assays, or known involvement in spinosad’s mode of action (table S2). These genes span target-site, detoxification, transport, and cuticular pathways. We tested whether SNPs associated with these candidates showed stronger evidence of association in focal GLM contrasts than expected by chance using two complementary enrichment approaches: hypergeometric overlap tests and Wilcoxon rank-sum analyses. Post-filtering, the dataset comprised 10,705 VEP-annotated SNPs in candidate gene regions, 4,982 SNPs excluding upstream and downstream flanking regions, and 120 missense SNPs predicted to alter amino acid sequence; enrichment analyses were conducted using both the full SNP set and the missense subset alone.

Hypergeometric overlap tests were performed using the *phyper* function in R. Each VEP-annotated feature (gene body ± 5 kb) was assigned the minimum GLM FDR across its SNPs, and we tested whether candidate genes were over-represented among features below the significance threshold. Tests were performed at two GLM FDR thresholds (FDR < 0.05 and FDR < 0.01) and for two SNP classes (all SNPs and missense SNPs only), giving four tests per contrast, using FDRs from 21-SNP sliding windows and, separately, from individual SNPs. Each set of *P*-values from the hypergeometric tests were adjusted across contrasts using the Benjamini–Hochberg procedure.

Wilcoxon rank-sum tests were used as a non-parametric enrichment approach to compare distributions of minimum GLM *P*-values between candidate genes and matched background gene sets, with the expectation that candidate genes would exhibit lower *P*-values in contrasts of adapted versus non-adapted populations. For each VEP-annotated feature (gene or genomic feature), the minimum GLM *P*-value across all SNPs assigned to that feature was used to represent it in the ranking, rather than the FDR-corrected value, to retain a broader distribution for ranking. Background gene sets were generated by matching each candidate gene to a random non-candidate gene of the same type (protein-coding, lncRNA, etc.), on the same chromosome arm, with similar gene length (±25%), and with similar associated SNP count (±25%). This matching procedure was repeated for 1,000 iterations, and Wilcoxon tests were conducted for each GLM contrast in every iteration. To account for genetic draft, we also tested whether the SNP with the lowest GLM *P*-value in each cluster of divergent SNPs (fig. S9) fell in candidate gene regions more often than the genome-wide proportion of SNPs (one-sided exact binomial tests).

##### 1.8.7 Population-level shift toward the pre-adapted genomic state

To test whether extinct and rescue populations differed in the extent to which they converged with pre-adapted populations, we defined a pre-adapted axis and projected each population sampled in August onto it. The axis comprised SNPs that differed between pre-adapted and unexposed populations at the end of the experiment in November (GLM FDR < 0.01; 428,205 SNPs). At each SNP, the direction of pre-adapted allelic change was defined as the direction in which pre-adapted populations differed from unexposed populations across September–November. For each exposed population sampled in August, we calculated the difference between its allele frequency and the mean frequency of unexposed populations in August, signed so that positive values indicate change in the direction of pre-adapted allelic change, and averaged this across all axis SNPs to obtain one value (Δ*p*) per population; each unexposed population was scored against the mean of the other unexposed populations. Because SNPs are not independent (linkage and genetic draft cause nearby SNPs to change together), Δ*p* largely reflects shifts across a limited number of linked regions rather than many independent loci; populations, not SNPs, were therefore treated as the unit of replication. Extinct and rescue populations (*n* = 5 each) were compared using an exact permutation test over all 252 assignments of the 10 exposed populations to two groups of 5, along with Welch’s *t*-test and a Wilcoxon rank-sum test.

##### 1.8.8 Temporal shift in the direction of pre-adapted allelic change

To test whether rescue populations shifted toward alleles that increased during laboratory selection for insecticide resistance (i.e., pre-adaptation) over the course of the experiment, we used pre-adapted and unexposed founder populations to create an axis of divergence. The axis comprised SNPs whose allele frequencies differed by more than 0.1 between the pre-adapted and unexposed founder populations sampled on July 16 (222,983 SNPs). At each SNP, the direction of pre-adapted allelic change was defined as the direction in which the pre-adapted founder population differed from the unexposed founder population. We projected every population sampled from August to November onto this axis. Because each treatment was represented by a single founder population, technical replicates of these founder populations (3 per treatment) were used only to exclude poorly measured SNPs and not as biological replication, so axis SNPs were selected by this threshold rather than by a test of significance. The SNPs used to score populations were chosen using only the founder samples, so the field samples being scored played no part in choosing them.

Populations were scored as stated previously (see *1.8.7 Population-level shift toward the pre-adapted genomic state*), except that at each time point, each population’s allele frequency was compared with the mean of the unexposed populations sampled at the same time point, with differences signed so that positive values indicate change in the direction of pre-adapted allelic change. Unexposed populations were again each scored against the mean of the remaining unexposed populations. At each time point, rescue and unexposed populations were compared using an exact two-sided permutation test over all assignments of populations.

##### 1.8.9 Non-parametric concordance analysis

To test whether extinct and rescue populations converged toward the genomic profiles of pre-adapted populations over time, we assessed overlap and genome-wide concordance among GLM treatment contrasts using two complementary approaches. Hypergeometric overlap tests (see *1.8.6 Functional enrichment tests*) were used to identify shared sets of significant outlier SNPs between ‘unexposed vs. pre-adapted’, ‘unexposed vs. extinct’, and ‘unexposed vs. rescue’ contrasts. Because such overlap tests are insensitive to genome-wide signals distributed across many loci of small effect, we additionally evaluated concordance across all loci using non-parametric Spearman rank correlations of GLM −log_10_(*P*) values (*123*), each assigned a positive or negative sign according to the direction of allele frequency differentiation in pre-adapted or exposed populations relative to unexposed populations. These signed values capture concordance in both the magnitude and direction of allele frequency change, independent of any significance threshold.

For August, Spearman correlations were calculated between ‘unexposed vs. pre-adapted’ and ‘unexposed vs. extinct’ contrasts, and between ‘unexposed vs. pre-adapted’ and ‘unexposed vs. rescue’ contrasts, to test whether extinct populations more closely resembled pre-adapted populations than rescue populations did at the same time point. Additionally, ‘unexposed vs. pre-adapted’ and ‘rescue vs. extinct’ contrasts were compared to assess divergence between rescue and extinct populations despite identical treatment conditions and a shared founding population. Each of these GLM-result comparisons was visualized using Rank-Rank Hypergeometric Overlap (RRHO) plots, a non-parametric rank-based analysis that assesses genome-wide concordance across two contrasts while incorporating all loci rather than only significant outliers (*124*), implemented using the *rrho2* package (*125*). RRHO plots were constructed on a random sample of 25,000 SNPs to limit computation time, and random sampling was performed 10 times per correlation of contrasts to confirm that all subsets showed similar patterns of concordance.

##### 1.8.10 Identification of convergent candidate loci across insecticide-exposed treatments

To identify loci putatively under convergent selection across insecticide-exposed treatments, we applied a sequential filtering cascade to GLM contrast results: (i) ‘unexposed vs. pre-adapted’ GLM contrasts were significant at FDR < 0.01 in both August and November, confirming that the signal persisted throughout the experiment; (ii) mean allele frequencies in unexposed populations in August were between 0.1 and 0.9, with a minimum allele frequency difference of 0.1 between unexposed and pre-adapted populations, excluding variants near fixation; (iii) for extinct population candidates in August, the ‘unexposed vs. extinct’ contrast was significant at FDR < 0.05; (iv) for rescue population candidates in November, the ‘unexposed vs. rescue’ contrast was significant at FDR < 0.05 and allele frequency change was in the same direction as in the ‘unexposed vs. pre-adapted’ contrast; and (v) candidate SNPs with FlyCADD scores > 0.6, from a deep learning model trained to predict the functional impact of *Drosophila melanogaster* variants (*85*), were classified as putatively functional, to prioritize likely direct targets of selection over variants closely linked to selected loci. Candidate SNPs and their associated genomic features are listed in table S10 (extinct populations, August), table S7 (rescue populations, November) and table S11 (three-way overlap). Allele frequency trajectories for all candidate loci within a set were plotted together, relative to the mean allele frequencies in the unexposed founder populations for each SNP.

This framework was applied at two experimental endpoints: August for extinct populations, representing their last sampling point before extinction, and November for rescue populations, representing the final sampling time point of the growing season. A three-way overlap analysis was subsequently conducted to identify SNPs and genes trending toward adaptive convergence across the two spinosad-exposed population endpoints and in pre-adapted populations; a relaxed threshold (FDR < 0.1 in all relevant GLM contrasts) was used for this three-way intersection to reduce the compounded stringency of requiring simultaneous significance across three contrasts.

To visualize convergent candidates and cross-check the GLM-based filtering cascade, rather than to identify candidates, we used a difference-of-differences residual approach (fig. S12). The core logic of this approach is that loci that are shared targets of selection should show significant allele frequency divergence from unexposed populations in all populations consistently exposed to the insecticide (i.e., pre-adapted and exposed populations), while showing limited divergence between the two exposed treatments.

For each 21-SNP window, residuals were calculated by subtracting the −log_10_(*P*) of the ‘pre-adapted vs. exposed’ GLM contrast from the −log_10_(*P*) of the ‘unexposed vs. pre-adapted’ contrast and, separately, from that of the ‘unexposed vs. exposed’ contrast, for August (extinct populations), November (rescue populations) and all time points combined. Before subtraction, the ‘pre-adapted vs. exposed’ values were linearly rescaled so that their maximum equaled the maximum of the contrast from which they were subtracted, and negative residuals were set to zero. Subtracting the ‘pre-adapted vs. exposed’ signal removes divergence specific to either treatment, so that high residuals mark regions in which pre-adapted and exposed populations both diverged from unexposed populations but not from each other. Such regions are the most likely to contain shared targets of selection from insecticide exposure, rather than loci that diverged through genetic draft in only one treatment.

Gene Ontology (GO) enrichment analyses were performed in R on genes overlapping, or within 5 kb of, convergent candidate SNPs at each time point and in the three-way overlap. Annotations were taken from FlyBase (GO annotation file, release 28 May 2026; annotations qualified with ‘NOT’ excluded) and the GO ontology (go-basic.obo, release 26 July 2026), with each gene’s annotations propagated to all ancestor terms through is_a and part_of relationships. Over-representation of each GO term among annotated candidate genes was tested with a one-sided hypergeometric test against a background of all annotated *D. melanogaster* genes (13,622 genes), following the approach implemented in BiNGO (*126*). *P*-values were corrected across all terms annotated in each candidate set using the Benjamini–Hochberg procedure, and terms with corrected *P* < 0.1 were considered enriched (table S8). All candidate SNP positions were annotated using Ensembl genome release BDGP6.46.

##### 1.8.11 Leave-one-out tests of parallel evolution in exposed populations

To test whether allele frequency change at outlier loci was parallel across replicate populations, rather than driven by a subset of them, we used a leave-one-out approach. For each exposed group for which replicate-level parallelism was tested (extinct populations in August, rescued populations in August, pre-adapted populations in August, and rescued populations in November), we fitted five additional per-SNP GLMs contrasting that group with unexposed populations, each omitting one of the five replicate populations. Outlier clusters were then defined from the remaining four populations, and the held-out population was tested for allele frequency change in the same direction at those clusters. As the held-out population’s data were not used to identify the clusters, this test avoids the circularity of evaluating a population at loci selected partly on its own allele frequency change.

To define clusters, each SNP was first assigned a score in each contrast, adapted from a previous clustering protocol (*43*): score 1, FDR < 0.2 and |Δ*p*| > 0.02; score 2, FDR < 0.05 and |Δ*p*| > 0.04; score 3, FDR < 0.01 and |Δ*p*| > 0.06; and score 0, all other SNPs. Here |Δ*p*| is the absolute difference in mean allele frequency between the exposed and unexposed populations included in the model, and each SNP received the highest score whose conditions it met.

To generate a null expectation for the spatial distribution of scores, we permuted scores among SNP positions within each chromosome. Observed and permuted scores were then compared using one-tailed Welch two-sample t-tests within windows of 501 consecutive SNPs (250 SNPs on either side of a central SNP), at 100-SNP steps. Windows with an FDR-corrected *P* < 0.05 that contained at least one SNP with an observed score of 3 were considered significant, and overlapping or adjacent significant windows were merged into clusters. Clusters were also defined in the same way from GLMs including all replicate populations, for extinct, rescue, and pre-adapted populations in August and rescue populations in November (fig. S9). Clusters for pre-adapted populations were used only to assess their overlap with clusters for extinct and rescue populations (see *2.4 Parallel evolution in exposed populations*).

Within each cluster, the SNP with the lowest FDR was selected as the focal SNP, and focal SNPs with FDR ≥ 0.05 were excluded. Each focal SNP was matched to a randomly drawn background SNP that lay outside all clusters and had FDR > 0.05. Matches were on the same chromosome arm, more than 50 kb from the focal SNP, with the same predicted variant consequence (the most severe VEP consequence per SNP, e.g. missense, synonymous, intronic or intergenic) and an unexposed founder allele frequency within 25% of that of the focal SNP. Focal SNPs with fewer than five possible matches were excluded. For the held-out population, we calculated a sign-corrected allele frequency difference at each SNP: its difference from the unexposed mean, multiplied by the sign of the difference between the remaining four populations and the unexposed mean. Positive values therefore indicate change in the same direction as the other replicates. Focal and matched background SNPs were compared with one-tailed Welch two-sample *t*-tests (focal > matched). Matched sets were drawn 100 times per held-out population, and we report the median *t*, the median *P* and the percentage of iterations with *P* < 0.05 (fig. S9; tables S3, S4, and S6). For completeness, each held-out population was also tested at clusters defined from all populations of the other exposed group or time point (fig. S9; tables S3, S4, and S6).

##### 1.8.12 Pleiotropic cost of resistance evolution

Alleles under directional selection during laboratory pre-adaptation may carry costs in the field environment (i.e., pleiotropic costs of resistance). We tested for such costs by asking whether, at loci that differed between pre-adapted and unexposed populations in August, allele frequencies within populations subsequently shifted in the direction of unexposed allelic change. At each SNP, the direction of pre-adapted allelic change was defined from the August ‘unexposed vs. pre-adapted’ GLM contrast as the direction in which pre-adapted populations differed from unexposed populations, and the direction of unexposed allelic change as the opposite direction. We tested temporal change in both pre-adapted and unexposed populations. Pre-adapted populations remained under insecticide exposure throughout, so continued selection for resistance would be expected to maintain or increase allele frequency change in the direction of pre-adapted allelic change. Shifts in the direction of unexposed allelic change in these populations therefore provides the strongest evidence of a cost. Unexposed populations, which never encountered the insecticide, test whether the same alleles are disfavored in the field environment without ongoing selection for resistance. We used three complementary approaches: (i) hypergeometric overlap tests, (ii) rank correlations across all loci, and (iii) comparisons of top outlier loci with matched background loci.

First, we identified loci with consistent temporal allele frequency change separately within unexposed and within pre-adapted populations, and tested whether each set overlapped with SNPs differentiated between unexposed and pre-adapted populations in August (GLM FDR < 0.05). Because GLMs revealed few significant temporal shifts in unexposed populations, loci were selected as those changing by more than 3% in the same direction in all, or all but one, populations of a group. Shifts in the direction of pre-adapted allelic change and in the direction of unexposed allelic change were tested separately using the hypergeometric distribution (*phyper* function in R). Results for unexposed populations are reported in table S1 and for pre-adapted populations in table S9; the main text reports the September-to-November interval, the longest interval that excludes the time point used to define the direction of allelic change.

Second, because overlap tests and outlier-based tests depend on filtering thresholds and miss weaker, genome-wide patterns, we also tested across all loci using Spearman rank correlations. For each temporal GLM contrast in unexposed and pre-adapted populations, −log_10_(*P*) values were assigned a positive sign when allele frequencies shifted in the direction of pre-adapted allelic change and a negative sign when they shifted in the direction of unexposed allelic change, and correlated with values from the August ‘unexposed vs. pre-adapted’ contrast (table S12). Negative correlations indicate that loci most strongly differentiated in August tended to shift in the direction of unexposed allelic change over time, consistent with a cost of resistance.

Third, we tested whether top outlier loci changed in the direction of unexposed allelic change more than comparable background loci. For each of four focal contrasts (the ‘unexposed vs. pre-adapted’, ‘unexposed vs. extinct’ and ‘unexposed vs. rescue’ contrasts in August, and the ‘unexposed vs. rescue’ contrast in November), we selected the 10,000 SNPs with the lowest GLM FDR values. Each was matched to a randomly drawn SNP outside the top 10,000 on the same chromosome arm, with the same VEP consequence, a founder allele frequency within 25% and a position more than 50 kb away; SNPs without VEP annotation, and SNPs with fewer than five candidate matches, were excluded. For each temporal contrast in unexposed and pre-adapted populations, signed −log_10_(*P*) values at focal and matched SNPs were compared using one-tailed Welch *t*-tests, with 100 iterations of matched-SNP selection (table S13). Significantly lower values at focal SNPs indicate change in the direction of unexposed allelic change beyond that of the genomic background. To show the direction of change, we also counted, for each focal contrast and temporal interval in pre-adapted populations, the number of top SNPs (restricted to mean allele frequency 0.15–0.85) changing in the direction of pre-adapted versus unexposed allelic change (fig. S13).

### Supplementary Results

#### 2.1 Polygenic resistance evolves rapidly from standing genetic variation

##### 2.1.1 Inversion polymorphisms are rare and do not underlie adaptive signals

The *D. melanogaster* genome contains several well-characterized chromosomal inversions that can influence linkage structure and allele frequency dynamics. To evaluate whether inversions contributed to the genomic signals observed in spinosad-exposed treatments, we screened the founder VCF and HAF-pipe allele frequency tables for diagnostic markers of 54 known inversions (473 SNPs). Only 27 inversion-associated markers were detected across all samples, of which one marker from each of two inversions (In(3L)P and In(3R)C) exceeded a mean allele frequency of 0.5 (i.e., 50%). The remaining inversion markers were either absent or present at frequencies below 0.1 (i.e., 10%), indicating that inversions were rare in the founding population. No inversion marker differed significantly between treatments or over time (per-SNP GLMs, all FDR > 0.05). These results suggest that known inversions do not underlie the strong allele frequency shifts observed in spinosad-exposed treatments and were not the driver of adaptive genomic responses.

##### 2.1.2 Sequencing and variant calling statistics

Read mapping and variant calling using *BCFtools* resulted in a mean per-sample sequencing coverage of 6.85× (SD = 2.99). Following HAF-pipe processing, estimated effective coverage increased substantially, to approximately 500–3,500× per sample (see *1.8.5 Genetic diversity estimates*). All downstream genomic analyses were conducted using HAF-pipe–imputed allele frequencies. HAF-pipe generated allele frequency estimates for 3.23 million variants genome-wide. We removed SNPs whose allele frequency variance across the samples was <0.001, which also filtered alleles with very low founder frequencies, and all sites on chromosome 4 (dot chromosome). We retained 1,095,839 SNPs for analyses.

##### 2.1.3 Genomic responses to spinosad exposure

Genome-wide allele frequency divergence in pre-adapted and exposed populations was strongly concentrated on chromosome arms 3L and 3R, consistent with patterns described in the main text. Of the 1,095,839 SNPs retained after filtering, 650,663 (59.4%) SNPs showed significant divergence between pre-adapted and unexposed populations (FDR < 0.01). Of these divergent sites, 60.1% were located on chromosome arms 3L and 3R. In the ‘unexposed vs. exposed’ contrast, 128,642 (11.7%) of filtered SNPs were significant at the same threshold, 83.7% of which were located on 3L and 3R (fig. S3).

To reduce noise in SNP-level estimates, we also averaged *P*-values and FDRs over 21-SNP sliding windows, identifying genomic regions in which neighboring SNPs were consistently differentiated rather than isolated SNPs with spurious signals. After smoothing, 134,215 (12.2%) of filtered SNPs were significant (FDR < 0.01) in the ‘unexposed vs. pre-adapted’ contrast and 4,189 (0.4%) in the ‘unexposed vs. exposed’ contrast, with 88.7% and 97.2%, respectively, located on chromosome arms 3L and 3R, suggesting that these two chromosome arms contain the majority of adaptive loci in the current study (Fig. 2). Significant SNPs in the two contrasts overlapped strongly: 3,727 of the 4,189 ‘unexposed vs. exposed’ outliers (89.0%) were also outliers in the ‘unexposed vs. pre-adapted’ contrast, 7.3-fold more than expected by chance (hypergeometric test at FDR < 0.01, *P* < 10⁻^300^), consistent with parallel genomic responses to spinosad exposure.

##### 2.1.4 Enrichment tests for spinosad candidate genes

Previously identified spinosad resistance genes (table S2) were not enriched among outlier SNPs. Across all-time point contrasts, no test using all SNPs was significant prior to correction. The lowest uncorrected *P* values were obtained for the ‘unexposed vs. pre-adapted’ contrast at FDR < 0.01 (36 candidate genes, uncorrected *P* = 0.11). Considering missense SNPs only, the lowest uncorrected *P* value was obtained for the ‘unexposed vs. exposed’ contrast at FDR < 0.05 (3 candidate genes, uncorrected *P* = 0.17). At the ‘GLM FDR < 0.01’ cutoff, 3 candidate genes reached genome-wide significance in the ‘unexposed vs. exposed’ all-time point contrast with all SNPs; in individual time point contrasts, none were significant from August through October and one was significant in November (Fig. 2D–E, table S2). At ‘GLM FDR < 0.05’ cutoff, 14 candidate genes were significant in the all-time point contrast, none were significant in within time point contrasts for August through October, and 6 candidates were significant in November.

Because enrichment tests based on discrete FDR thresholds can be sensitive to cutoff choice, we additionally performed rank-based Wilcoxon tests. Each candidate gene was represented by the 21-SNP window minimum GLM *P*-value across its SNPs and compared with 1,000 sets of matched background genes (see *1.8.6 Functional enrichment tests*). Across iterations, mean Wilcoxon test *P*-values were 0.43 for ‘unexposed vs. pre-adapted’ and 0.40 for ‘unexposed vs. exposed’. Only 0.2% of ‘unexposed vs. pre-adapted’ iterations and 0.7% of ‘unexposed vs. exposed’ iterations yielded *P* < 0.05.

To confirm that the lack of enrichment among candidate genes was not a consequence of smoothing over 21-SNP windows, we repeated the hypergeometric tests using SNP-level FDRs. More candidate genes contained significant SNPs at FDR < 0.01 (45 genes in the ‘unexposed vs. exposed’ contrast and 57 in the ‘unexposed vs. pre-adapted’ contrast), but overlap remained non-significant (*P* = 0.40 and *P* = 0.68, respectively; fig. S3). The presence of significant SNPs in candidate genes without corresponding enrichment is consistent with genetic draft from nearby selected loci, given the broad selective sweeps observed across chromosome arms 3L and 3R, rather than with direct selection on previously characterized resistance variants. Restricting the analysis to the top SNP within each cluster of divergent SNPs likewise revealed no enrichment of candidate genes in any contrast (all binomial *P* ≥ 0.49).

Finally, we tested whether spinosad candidate SNPs exhibited larger absolute allele frequency changes than matched background SNPs. Matched sets were generated by pairing each candidate or outlier locus with up to 100 randomly selected loci on the same chromosome arm with similar founder allele frequencies within unexposed populations (±25%) and the same VEP-predicted consequence. Mean allele frequency changes across matched loci were compared per locus with observed allele frequency changes using Welch two-sample *t*-tests. No significant differences were detected in either contrast (‘unexposed vs. exposed’: *P* = 0.78; ‘unexposed vs. pre-adapted’: *P* > 0.99).

We repeated these analyses for the August contrasts to test whether candidate genes were targets of selection during early adaptation (Fig. 3D–F). Neither hypergeometric overlap tests nor Wilcoxon rank-sum tests detected enrichment of candidate genes in the ‘unexposed vs. extinct’, ‘unexposed vs. rescue’ or ‘unexposed vs. pre-adapted’ contrasts (hypergeometric tests, all uncorrected *P* ≥ 0.30; Wilcoxon rank-sum tests, mean *P* = 0.42, 0.46 and 0.63, respectively, across 1,000 matched-SNP iterations). To confirm that this lack of enrichment was not a consequence of window-based smoothing, we repeated the hypergeometric tests using SNP-level FDRs (FDR < 0.05). Only the ‘unexposed vs. pre-adapted’ contrast showed nominal enrichment (54 of 57 candidate genes; uncorrected *P* = 0.041), and this did not remain significant after Bonferroni correction for the three August contrasts (adjusted *P* = 0.12). Neither the ‘unexposed vs. extinct’ (*P* = 0.98) nor the ‘unexposed vs. rescue’ (*P* = 0.37) contrast was enriched (fig. S4).

Together, these analyses indicate that although genomic responses to spinosad exposure were strong and highly parallel across treatments, previously identified spinosad resistance genes did not show stronger divergence than comparable genes in the same genomic regions. Because selection signals extended across large portions of chromosome arms 3L and 3R, these analyses cannot exclude a contribution of individual candidate genes, but they provide no evidence that adaptation was disproportionately concentrated in them.

#### 2.2 Extinction occurs despite strong adaptive responses

##### 2.2.1 Genetic diversity and bottleneck effects in rescue versus extinct populations

As an additional quality-control analysis, we tested whether the stronger allele frequency divergence from unexposed populations in extinct than in rescue populations could be explained by reductions in genetic diversity associated with severe bottlenecks. This concern is especially relevant for extinct populations, which experienced pronounced demographic collapse leading to extinction and may have lost genetic diversity.

To evaluate this possibility, we estimated Watterson’s θ for all samples collected in August for each chromosome and genome wide (table S5). Diversity estimates were generated from HAF-pipe-imputed allele frequencies and analyzed using GLMs to test for differences among treatments. Under both effective-coverage assumptions (500× and 3,500×), genome-wide Watterson’s θ was lower in pre-adapted populations than in unexposed populations (*P* = 3.47 × 10⁻^6^ and *P* = 3.90 × 10⁻^7^, respectively). In contrast, genetic diversity did not differ detectably between extinct and rescue populations, either genome-wide (*P* = 0.95 and *P* = 0.93) or for any chromosome (all P ≥ 0.86), despite large differences in census size and demographic trajectories. Although statistical power to detect small differences is limited with five replicates per group, these results indicate that the pronounced allele frequency divergence observed in extinct populations is unlikely to be driven by generalized losses of genetic diversity or extreme bottleneck effects.

##### 2.2.2 Extinct populations shifted further toward the pre-adapted genomic state

By August, extinct populations had shifted in the direction of pre-adapted allelic change by a mean Δ*p* of 0.035 (range 0.015–0.060 across replicates) at SNPs differentiating pre-adapted and unexposed populations in November, compared with 0.005 (range −0.009 to 0.018) for rescue populations and 0.088 (range 0.076–0.106) for pre-adapted populations (Fig. 3J). Extinct and rescue populations differed significantly (exact permutation test, *P* = 0.016; Welch’s *t*-test, *t* = 3.23, *P* = 0.018; Wilcoxon rank-sum test, *P* = 0.016); only 1 of the 251 alternative assignments of populations to groups produced a larger difference, by exchanging the extinct population with the lowest value (Δ*p* = 0.015) and the rescue population with the highest value (Δ*p* = 0.018), the only populations whose values fell within the range of the other group (Fig. 3J).

#### 2.3 Pre-adaptation is costly but effective, and shares a genomic basis with de novo rescue

##### 2.3.1 Genomic targets of early adaptation in extinct populations

In August, 619 SNPs in extinct populations passed initial GLM-based filtering criteria, 499 of which were within 5 kb of annotated genomic features and 76 of those had FlyCADD scores > 0.6. After filtering for convergence between exposed and pre-adapted populations, 359 SNPs with intermediate allele frequencies (0.1–0.9) were significant in both ‘unexposed vs. extinct’ and ‘unexposed vs. pre-adapted’ contrasts, 53 of these with FlyCADD scores > 0.6 (table S10). All convergent outlier SNPs in August were confined to chromosome arms 3L and 3R. Gene Ontology enrichment analysis identified ‘aldehyde oxidase activity’ (table S8), driven by a single peak encompassing all four aldehyde oxidase genes (Aox1–4) on chromosome arm 3R. Because these duplicated genes are tightly linked, this enrichment reflects a single genomic region rather than independent signals at multiple loci. The top-ranked SNP in this region was a missense variant in aldehyde oxidase 2 (Aox2; FlyCADD = 0.95).

##### 2.3.2 Genomic targets of resistance in rescue populations

Early in the experiment in August, rescue populations showed no significant parallel divergence from unexposed populations after GLM result smoothing. We checked counts of outliers prior to rolling window smoothing and found 134 significant SNPs (FDR < 0.05), 127 of which were near to or within annotated features; 14 had FlyCADD scores > 0.6. Even with this relaxed detection criteria, we found only 4/134 significant SNPs were among the 32,253 significant SNPs in the August ‘unexposed vs. extinct’ contrast as well, no more than expected by chance (hypergeometric test, *P* = 0.56) (fig. S8).

At the final experimental time point in November, rescue populations exhibited a broader but largely distinct set of outlier loci associated with spinosad resistance. A total of 4,429 SNPs were significant in GLM contrasts between exposed and unexposed populations (FDR < 0.05), of which 3,844 were within 5 kb of annotated genomic features and 517 had FlyCADD scores > 0.6. Applying the same convergence filtering framework, 441 significant SNPs with intermediate allele frequencies additionally passed divergence criteria in the ‘unexposed vs. pre-adapted’ contrast, 81 of which had FlyCADD scores > 0.6 (table S7). As in August, convergent outlier SNPs were almost entirely confined to chromosome arms 3L and 3R (438 of 441). GO enrichment analysis (table S8) identified ‘AAA codon-amino acid adaptor activity’ and ‘L-galactose dehydrogenase activity’ as the most significantly enriched categories, driven respectively by a tandem cluster of four tRNA:Lys-TTT genes and a cluster of three tightly linked L-galactose 1-dehydrogenase genes, both on chromosome 3R. As with the aldehyde oxidase cluster in August, these enrichments reflect individual genomic regions rather than independent signals.

##### 2.3.3 Overlap and divergence of adaptive architectures across time points

Residual Manhattan plots (fig. S12) show, for each 21-SNP window, the −log_10_(*P*) of the contrast between exposed (or pre-adapted) and unexposed populations minus the −log_10_(*P*) of the contrast between pre-adapted and exposed populations, with the latter rescaled to the same maximum and negative values set to zero. High residuals therefore mark regions in which exposed populations diverged from unexposed populations while remaining similar to pre-adapted populations. Convergent candidate SNPs identified in August (extinct populations) and November (rescue populations) fell within the major residual peaks, where exposed and pre-adapted populations showed reduced differentiation despite broad genomic divergence from unexposed populations. This pattern is consistent with these variants being, or being closely linked to, targets of selection rather than being subject only to genetic draft.

The specific SNPs showing convergence with pre-adapted populations were largely non-overlapping between extinct and rescue populations (17 SNPs shared between the 359 August and 441 November convergent candidates). Outliers in extinct and rescue populations in August likewise overlapped little, both with and without 21-SNP window smoothing (Fig. 3D–F; fig. S8). To identify convergent outliers shared across extinct, rescue and pre-adapted populations, we applied a relaxed threshold (FDR < 0.1 in all four contrasts) and found 193 SNPs, 26 of which had FlyCADD scores > 0.6 (table S11). Both outlier gene sets were enriched for oxidoreductase activities: aldehyde oxidase activity in August and L-galactose dehydrogenase activity in November (table S8). In each case, however, the enrichment was driven by a single cluster of tightly linked genes, and this functional overlap accounts for only a small proportion of the outlier loci in each list.

#### 2.4 Parallel evolution in exposed populations

To assess parallelism in exposed populations, we compared focal outlier SNPs within clusters with matched background SNPs. Clustering of GLMs including all replicate populations identified 49 clusters for extinct populations in August, 3 for rescue populations in August, and 371 for rescue populations in November (fig. S9). Clustering of four-replicate GLMs yielded 45–65 clusters for extinct populations in August, 1–16 for rescue populations in August, and 195–231 for rescue populations in November, depending on the held-out replicate. Concordance *t*-tests were run for each held-out replicate within each cluster set, allowing us to assess parallel change across candidate regions for spinosad resistance (fig. S9; tables S3, S4, and S6; see *1.8.11 Leave-one-out tests of parallel evolution in exposed populations*).

##### 2.4.1 Parallelism among exposed populations in August

Leave-one-out tests confirmed parallel evolution among extinct populations in August. In clusters defined by the four remaining extinct populations, focal SNPs showed greater concordant allele frequency change in the held-out population than matched background SNPs in 100% of iterations for every held-out population (*n* = 100 iterations per population; median *t* = 7.16, median P = 8.05 × 10⁻^11^; table S3). In contrast, rescue populations showed little clustered divergence in August, and allele frequency change at these clusters was not detectably parallel: leave-one-out runs identified 1–16 clusters, and focal SNPs showed greater concordant change than matched background SNPs in 5.5% of iterations (median *t* = 0.91, median *P* = 0.20; table S4). One rescue population could not be tested, as the single cluster identified without it lacked a focal SNP with a sufficient number of matched background SNPs.

##### 2.4.2 Parallelism in rescue populations in November

By November, leave-one-out tests revealed strong parallel evolution among rescue populations. In clusters defined by the four remaining rescue populations, focal SNPs showed greater concordant allele frequency change in the held-out population than matched background SNPs in 98.4% of iterations, and in at least 96% for every held-out population (median t = 4.25, median *P* = 1.34 × 10⁻^5^; table S6). Together with the August results (see *2.4.1 Parallelism among exposed populations in August*), these tests show that allele frequency change at each group’s outlier loci relative to unexposed populations was parallel across replicate extinct populations by August and across replicate rescue populations by November, rather than driven by a few populations.

#### 2.5 Costs of resistance evolution

##### 2.5.1 Hypergeometric overlap tests

In nearly all temporal contrasts in both unexposed and pre-adapted populations, more SNPs with parallel allele frequency change moved in the direction of unexposed allelic change than in the direction of pre-adapted allelic change (all contrasts except August vs. September in unexposed populations; Tables S1 and S9). The same asymmetry holds for overlaps with ‘unexposed vs. pre-adapted’ August outlier loci. In unexposed populations, overlaps with these outliers were significantly larger than expected for the August–October, August–November and September–November intervals in the direction of unexposed allelic change, with no significant overlaps in the direction of pre-adapted allelic change. In pre-adapted populations, all temporal intervals except October–November showed significant overlap in the direction of unexposed allelic change, and none in the direction of pre-adapted allelic change.

These results are consistent with resistance carrying a cost: under field conditions, loci shifted in the direction of unexposed allelic change more often than expected by chance. This held even in unexposed populations, which showed few significant loci in temporal GLM contrasts and required modified filtering criteria to detect allele frequency change. Pre-adapted populations, moreover, shifted more strongly in the direction of unexposed allelic change than unexposed populations did, despite continued insecticide exposure throughout the experiment.

##### 2.5.2 Locus rank correlations

At the whole-genome level, temporal allele frequency changes were also consistent with a cost of insecticide resistance. Using sign-corrected −log_10_(*P*) values, we detected significant negative Spearman rank correlations between the August ‘unexposed vs. pre-adapted’ contrast and the majority of temporal contrasts in both unexposed and pre-adapted populations (pre-adapted: ρ = −0.13 to −0.31; unexposed: ρ = −0.04 to −0.20; table S12). These negative correlations indicate broad allele frequency change over time in the direction of unexposed allelic change rather than pre-adapted allelic change, consistent with selection against resistance alleles or variants linked to them. Although the directions of pre-adapted and unexposed allelic change were defined from the August contrast, temporal contrasts that exclude August show the same pattern. This signal is therefore not an artifact of the time point used to define the direction of change, and is consistent with a cost that persisted across the experiment in both continuously exposed and unexposed populations.

##### 2.5.3 Top outlier temporal trends

Two features distinguish this test from those above: shifts at outlier loci are compared with matched background loci, controlling for allele frequency change expected regardless of selection history, and it extends to loci distinguishing extinct and rescue populations from unexposed populations. The 10,000 loci most strongly differentiated (lowest GLM FDR) in the August ‘unexposed vs. pre-adapted’ contrast shifted, relative to 100 sets of matched background SNPs, in the direction of unexposed allelic change in five of six temporal contrasts in unexposed populations (all except August–September), and in four of six contrasts in pre-adapted populations (August–October, August–November, September–October, and September–November; table S13). Loci distinguishing extinct populations in August shifted in the direction of unexposed allelic change in all contrasts except August–September, in both unexposed and pre-adapted populations. Loci distinguishing rescue populations showed no such shifts in pre-adapted populations, in either August or November. In unexposed populations, they shifted in the direction of unexposed allelic change in three contrasts for the August set (September–October, September–November and October–November) and in all six contrasts for the November set. The direction of change at these loci in pre-adapted populations is shown in fig. S13.

Across these tests, loci differentiating pre-adapted from unexposed populations shifted in the direction of unexposed allelic change despite continued insecticide exposure in pre-adapted populations. This may reflect the genomic consequences of prior laboratory selection. Pre-adapted populations entered the field already partially resistant and carrying broad signatures of sweeps arising from selection and linkage, so the loci differentiating them from unexposed populations include many variants linked to, rather than directly involved in, resistance. Core resistance loci may have remained under positive selection in the field while selection reduced the frequency of some deleterious linked variants, producing net movement toward unexposed allele frequencies. Alternatively, alleles favored during laboratory selection may have been selected against under field conditions, independently of insecticide resistance. Distinguishing a cost of resistance from selection against linked variants, or against alleles favored under laboratory conditions, would require experiments that separate these sources of selection (e.g., pre-adapted populations maintained with and without insecticide in both laboratory and field environments), together with identification of the causal resistance loci.

---

**Fig. S1.**
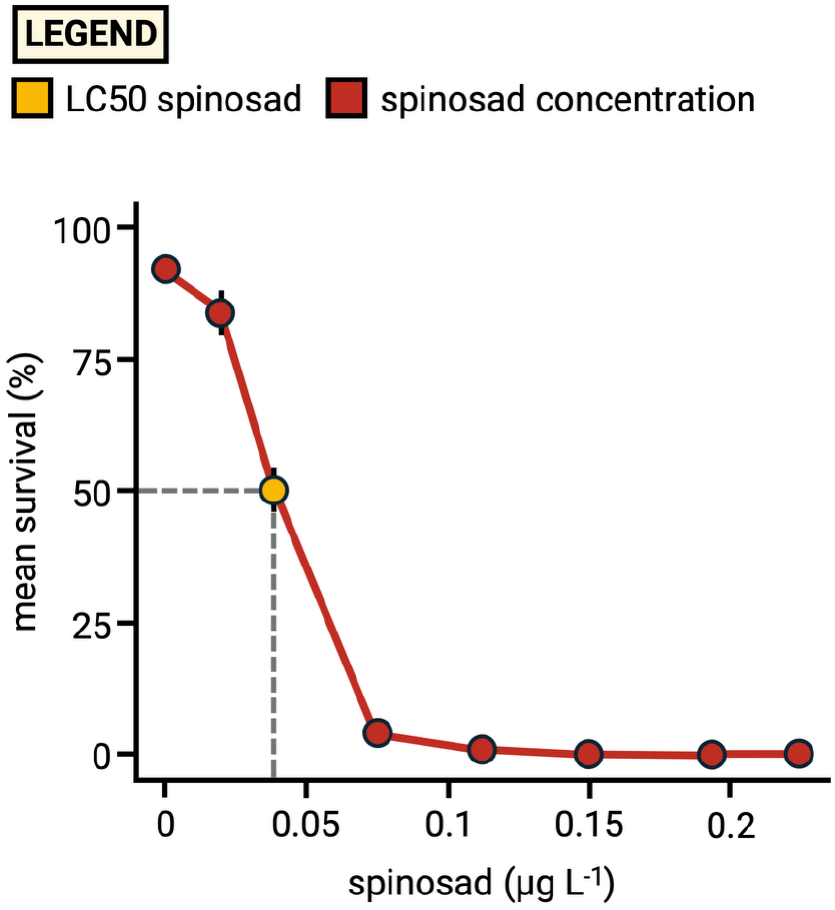
Concentration-response relationship for resistance measured as % survival from egg to adult in *Drosophila melanogaster* exposed to the insecticide spinosad. Mean survival (± 1 SE) across a range of spinosad concentrations (µg L⁻¹) are plotted. The lethal concentration at which 50% of *D. melanogaster* experienced mortality (i.e., LC50) was measured and used as the dose in all experimental treatments exposed to toxic media (see *Materials and Methods*).

**Fig. S2.**
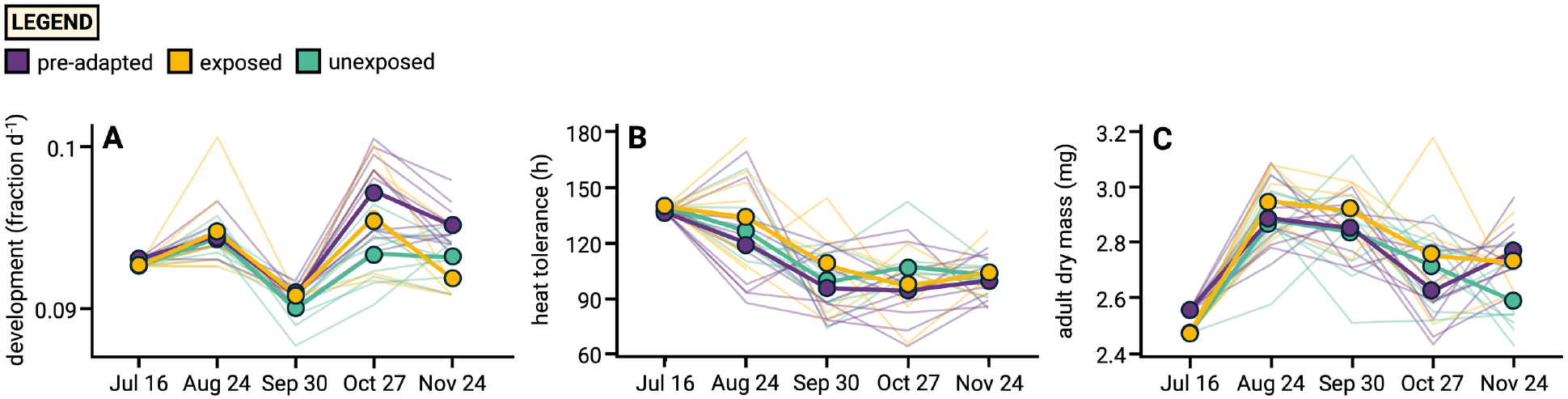
Phenotypic evolution of development rate, heat tolerance, and adult dry mass in unexposed, exposed, and pre-adapted populations. Phenotypic trajectories were measured at each time point following two generations of common garden rearing, with all assays conducted on flies reared on spinosad-amended media at the LC50 concentration. **(A)** Development rate expressed as the fraction of development completed per day [1/(total hours/24)]. **(B)** Heat tolerance, measured as the mean time to death among 15 females in a water bath at 53°C. **(C)** Adult dry mass, computed as the average weight among three pools of five females, dried at 55°C for 24 h. Thin colored lines represent the mean trajectory of each population, and thick colored lines show the treatment means (averaged across cages).

**Fig. S3.**
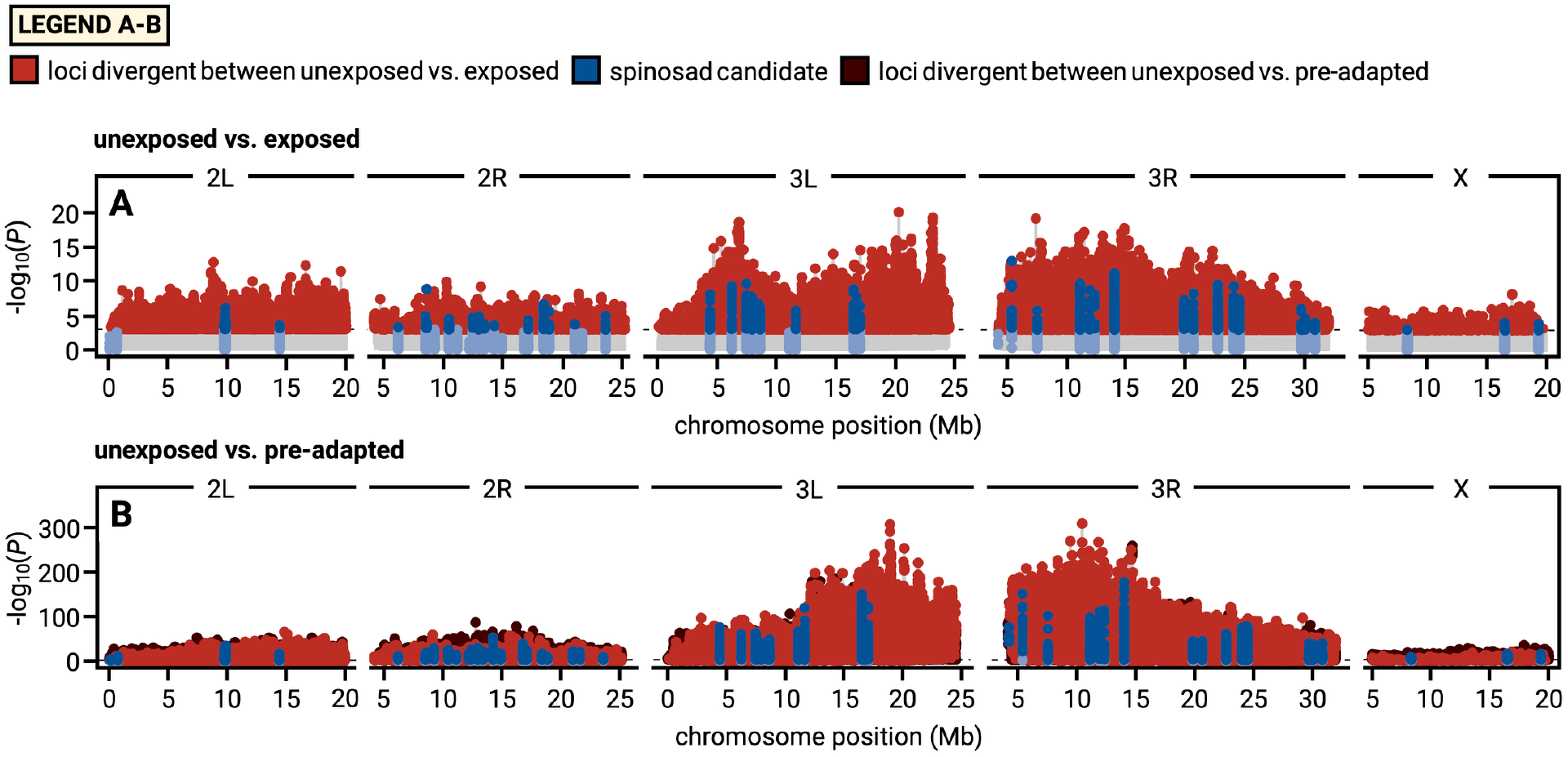
Genomic patterns of spinosad resistance across all time points, without sliding-window smoothing. Genome-wide signatures of allele frequency divergence between unexposed field populations and (**A**) exposed or (**B**) pre-adapted populations across all time points, shown as −log_10_(*P*) from generalized linear models applied to individual SNPs. These panels correspond to Fig. 2D-E, in which the same contrasts are summarized using 21-SNP rolling windows. In (**A**) and (**B**), red points denote loci divergent between unexposed and exposed populations (FDR < 0.01), dark blue points denote significant loci within previously characterized spinosad resistance candidate genes, and light blue points denote non-significant candidate loci. In (**B**), dark brown points denote loci divergent between unexposed and pre-adapted populations (FDR < 0.01).

**Fig. S4.**
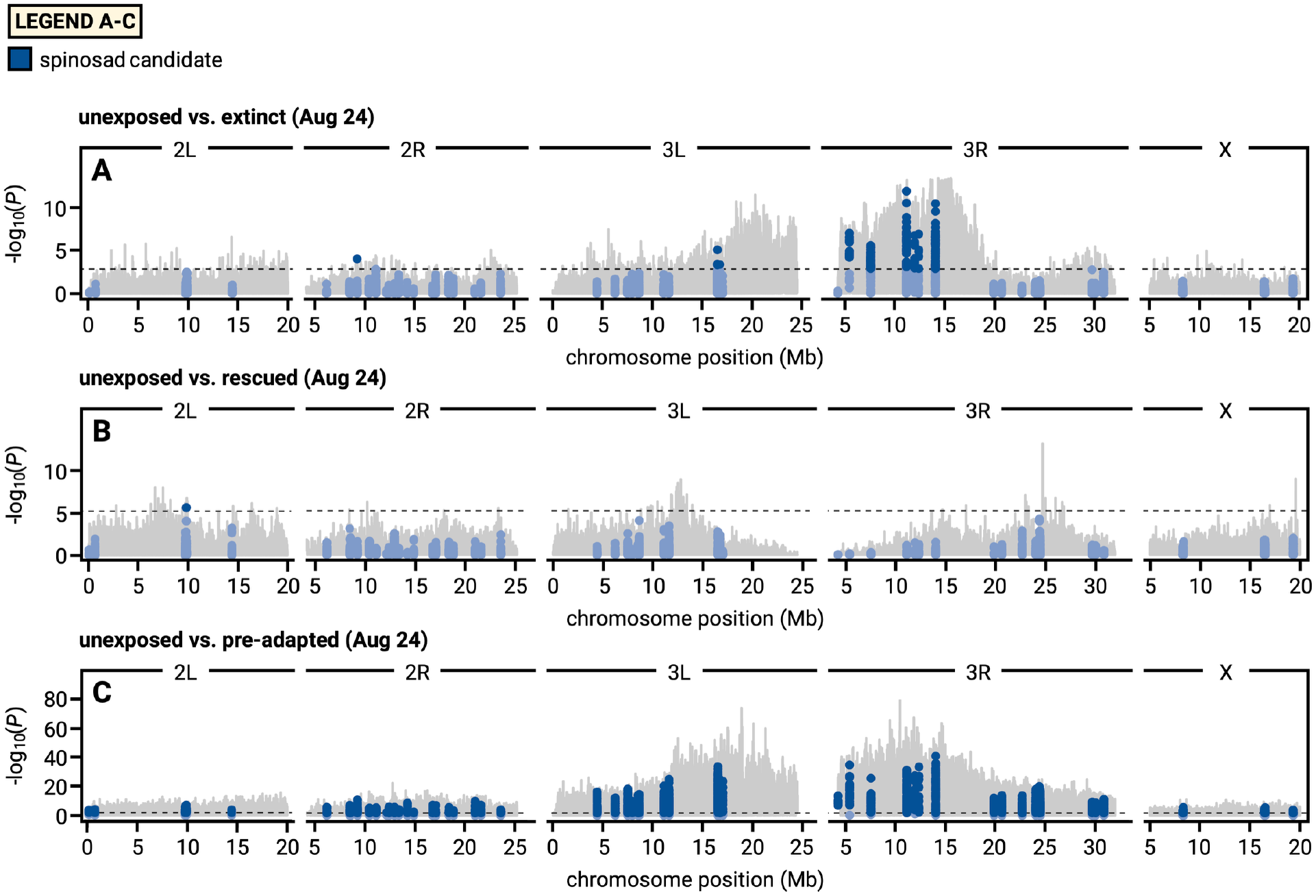
Spinosad resistance candidate genes among genome-wide signals of divergence in insecticide-treated populations in August. Genome-wide allele frequency divergence in August in (**A**) extinct, (**B**) rescue, and (**C**) pre-adapted populations relative to unexposed populations, shown as −log_10_(*P*) from generalized linear models applied to individual SNPs. Dark blue points denote significant loci within previously characterized spinosad resistance candidate genes (Table S2), and light blue points denote non-significant candidate loci. Candidate genes were not significantly enriched among outlier SNPs in any contrast after correction for multiple testing. In the ‘unexposed vs. pre-adapted’ contrast, most genes genome-wide contained significant SNPs, limiting the scope for enrichment (see *2.1.4 Enrichment tests for spinosad candidate genes*).

**Fig. S5.**
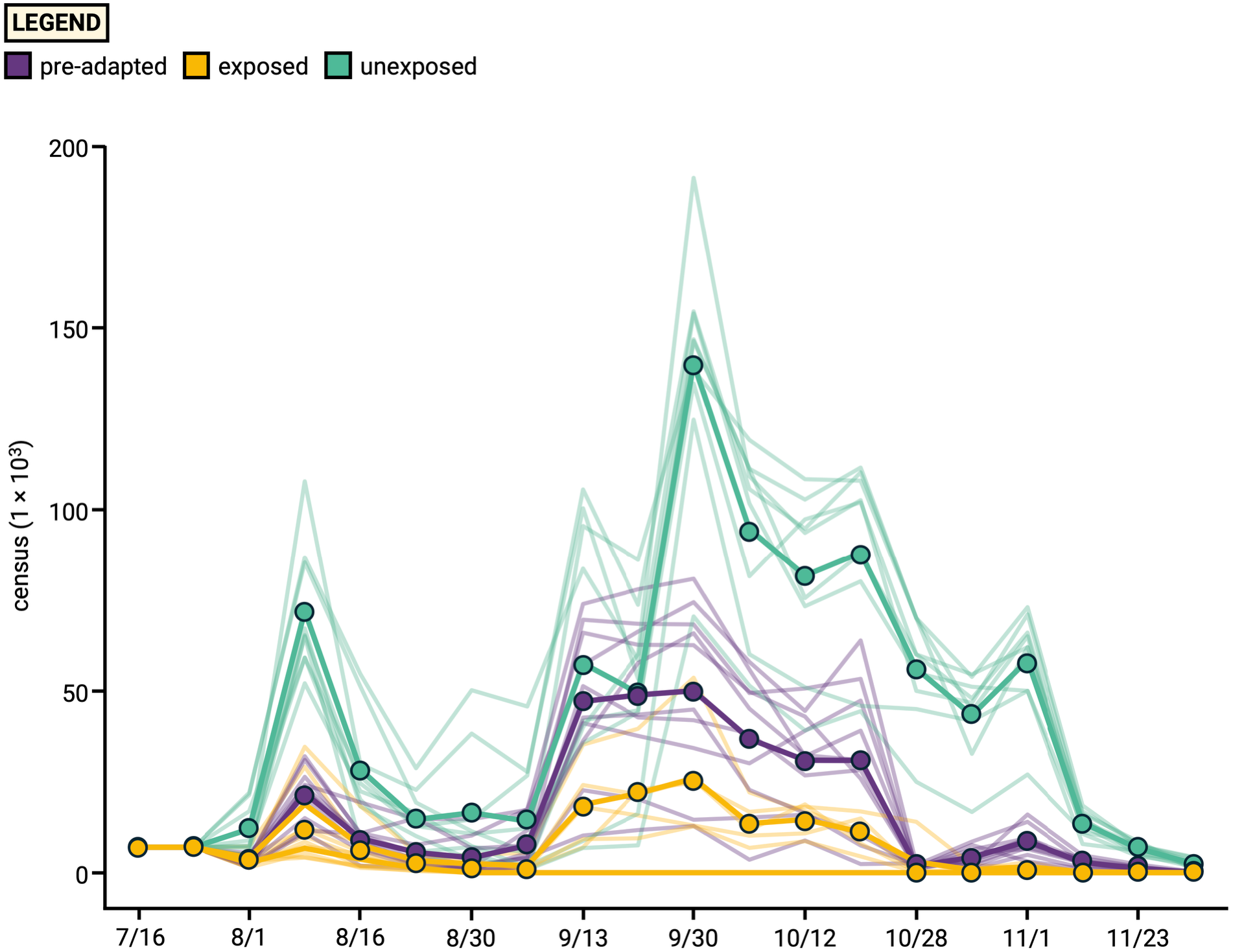
Adult census size for all populations over the full time series. Adult census size was estimated weekly for each replicate across five months of seasonal change (thick line, population mean; thin line, replicate).

**Fig. S6.**
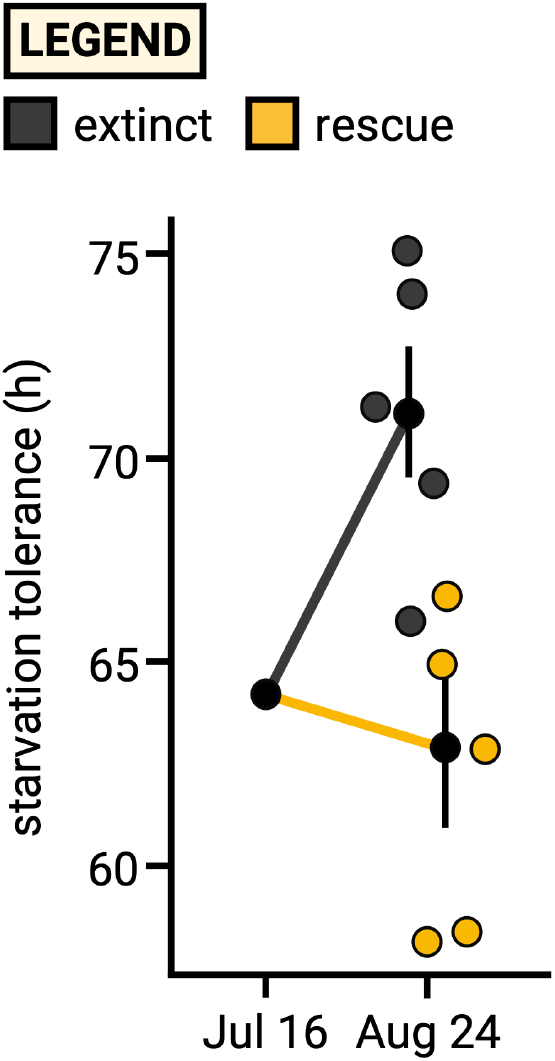
Starvation tolerance in August is associated with failed and successful evolutionary rescue. Points represent individual cages; and points with vertical symbols denote group means ± SE.

**Fig. S7.**
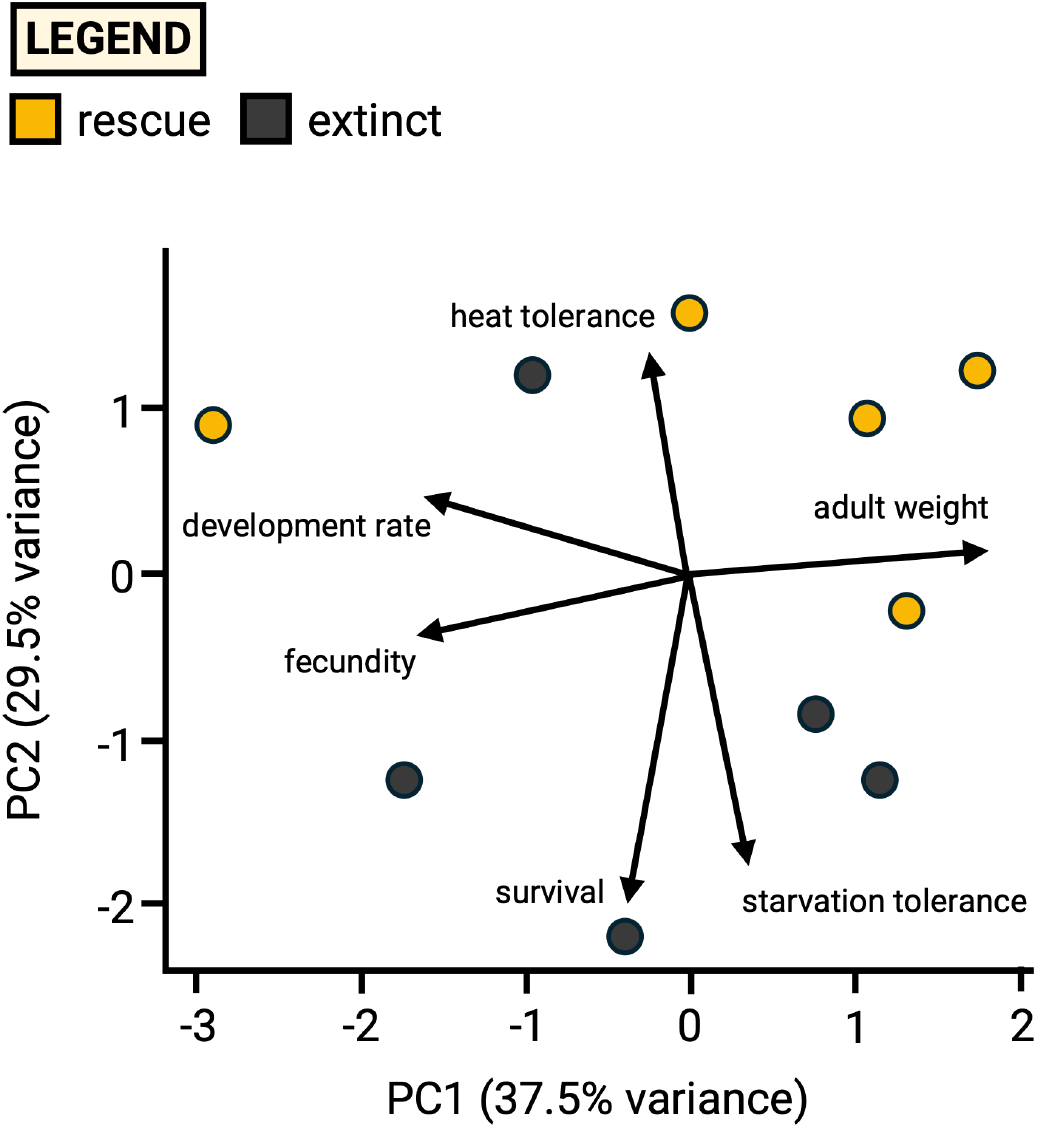
Principal component analysis of multivariate phenotypic divergence among exposed populations in August. Points represent mesocosm means for extinct and rescue populations. Arrows indicate trait loadings for resistance, development rate, fecundity, starvation tolerance, heat tolerance, and adult dry mass. PC1 (37.9% of variance) primarily reflects variation in adult dry mass, fecundity, and development rate, with dry mass loading opposite to the other two traits. PC2 (29.7% of variance) separates extinct from rescue populations: extinct populations scored lower on PC2 (β = −1.77, *P* = 0.024, R² = 0.49). Extinct populations therefore diverged early toward higher resistance and starvation tolerance, and toward lower heat tolerance.

**Fig. S8.**
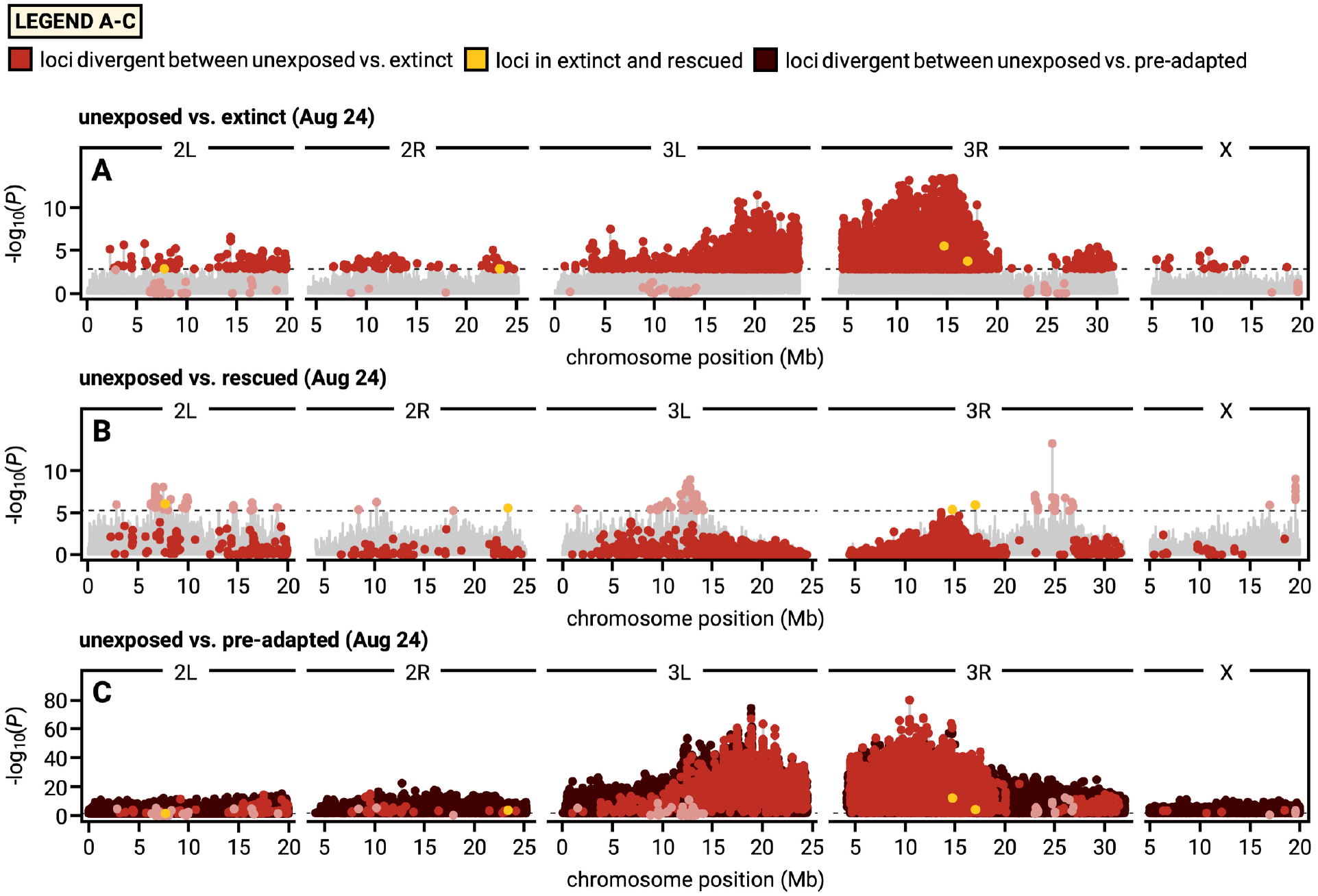
Genomic patterns of spinosad resistance in August, without sliding-window smoothing. Genome-wide allele frequency divergence in August in **(A)** extinct, **(B)** rescue, and (**C**) pre-adapted populations relative to unexposed populations, shown as −log_10_(*P*) from generalized linear models applied to individual SNPs. These panels correspond to Fig. 3D-F, in which the same contrasts are summarized using 21-SNP rolling windows. Red points denote loci divergent between unexposed and extinct populations (FDR < 0.05 in panel A) and are shown across all three panels to facilitate comparison of adaptive signals across treatments. Light red points denote loci divergent between unexposed and rescue populations (FDR < 0.05 in panel B) and are likewise shown in all panels. Yellow points denote SNPs significant in both extinct and rescue populations relative to unexposed populations. In panel C, brown points denote loci divergent between unexposed and pre-adapted populations (FDR < 0.05).

**Fig. S9.**
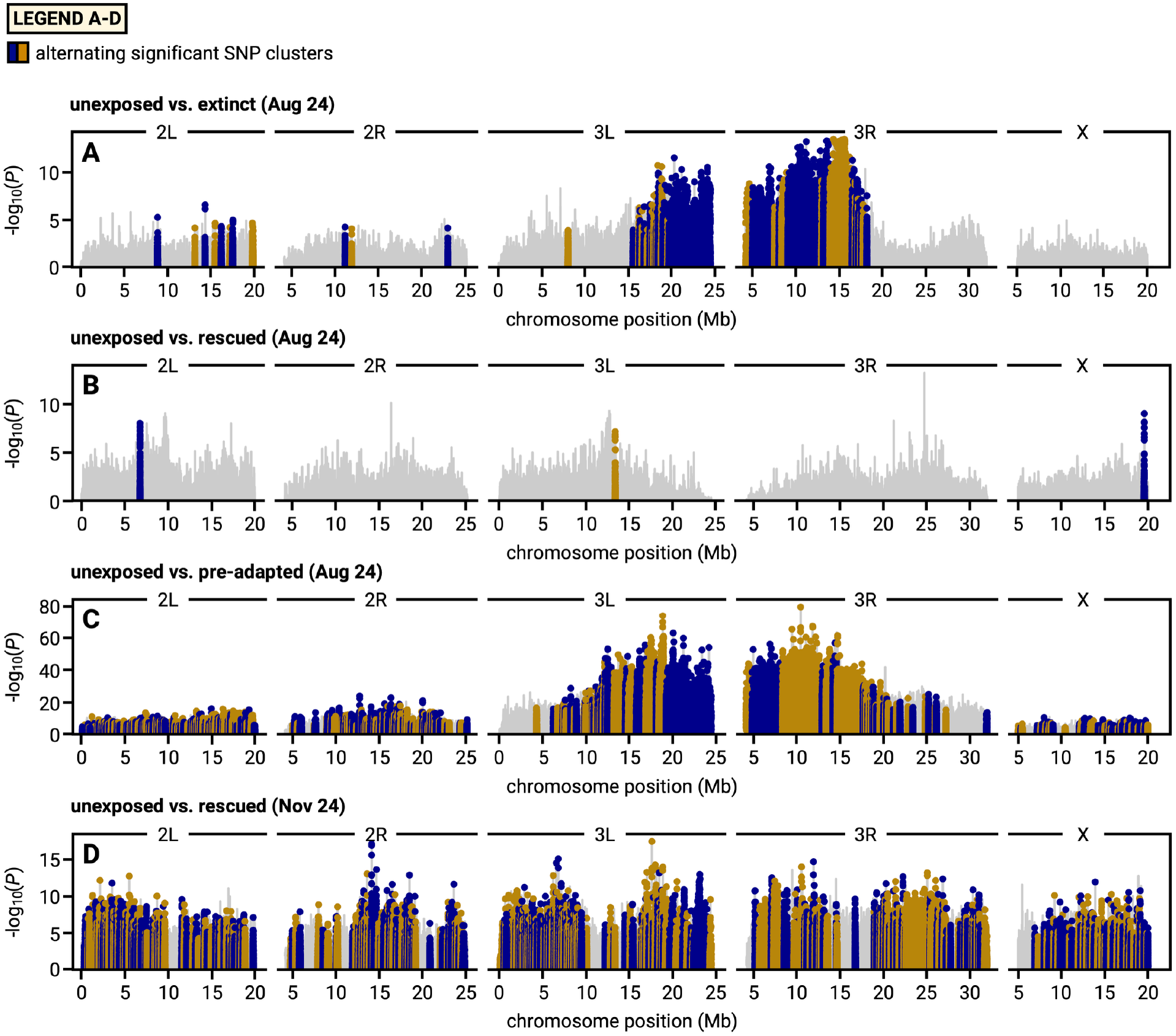
Genomic clusters of divergence from unexposed populations. Manhattan plots of −log_10_(*P*) from per-SNP GLMs contrasting unexposed populations with (**A**) extinct populations in August, (**B**) rescue populations in August, (**C**) pre-adapted populations in August, and (**D**) rescue populations in November. Colored points mark SNPs within significant clusters, shown in alternating colors to distinguish adjacent clusters (A, 49; B, 3; C, 334; D, 371), defined from all replicate populations of each group as described in *1.8.11 Leave-one-out tests of parallel evolution in exposed populations*. Gray lines show all other SNPs.

**Fig. S10.**
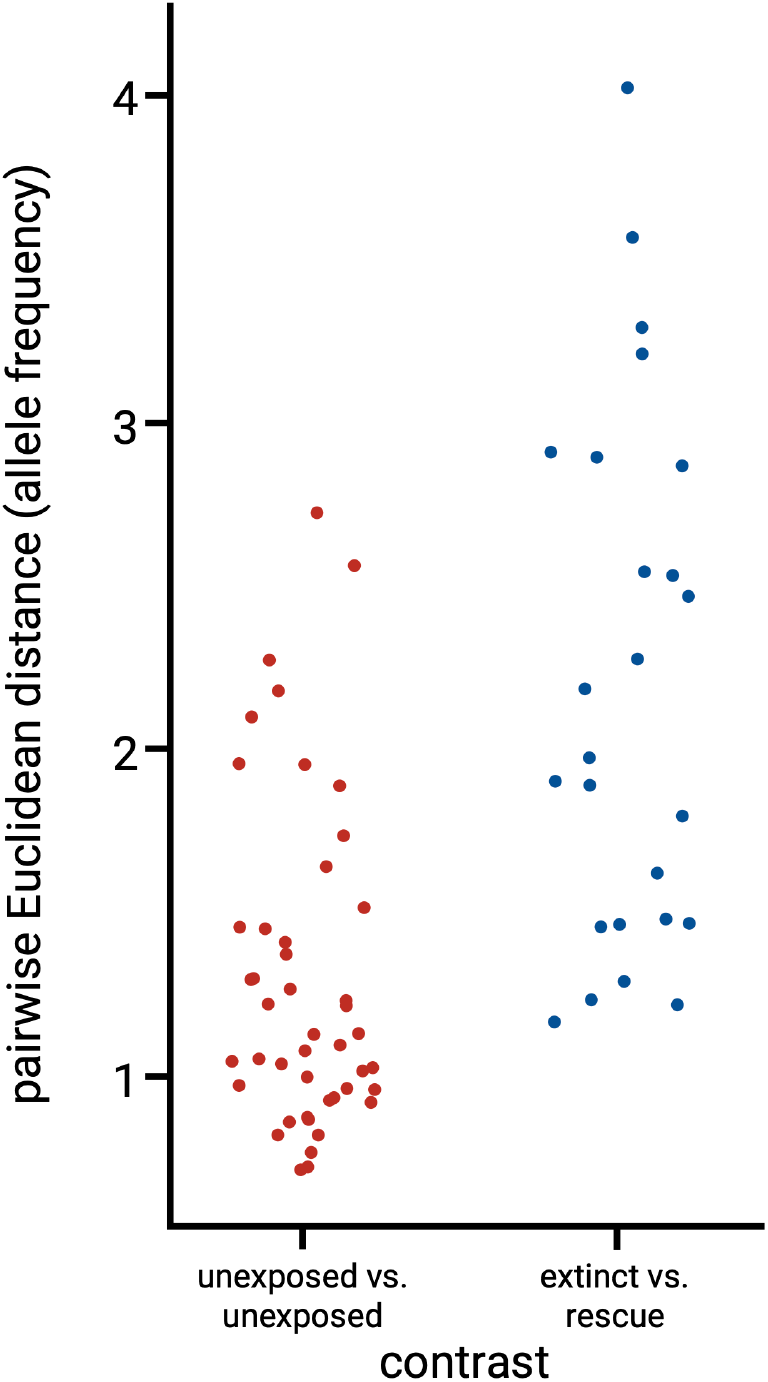
Pairwise Euclidean distances in allele frequencies at November convergent candidate loci, calculated among populations sampled in August. Each point represents the distance between a pair of populations at loci identified as convergent outliers between pre-adapted and rescue populations in November. Lower distance values indicate greater similarity in allele frequencies between population pairs. Contrast types along the x-axis reflect comparisons of populations either within unexposed or between extinct and rescue treatment groups.

**Fig. S11.**
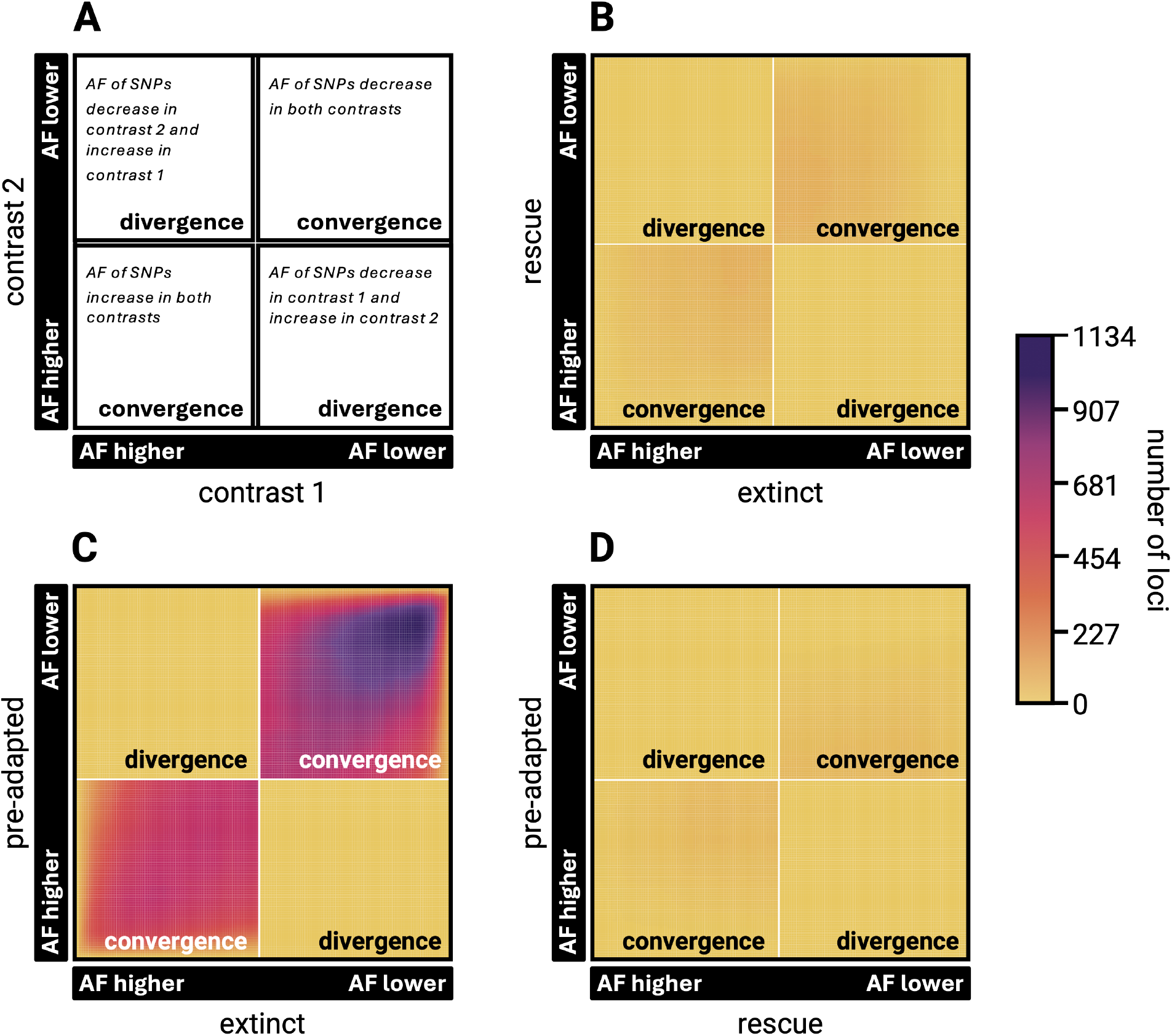
Rank-rank hypergeometric overlap (RRHO) analyses comparing genome-wide concordance in the direction and significance of allele frequency change across treatment contrasts. **(A)** Schematic illustrating RRHO quadrant interpretation: concordant signals appear in the top-right (SNPs decreasing in both contrasts) and bottom-left (SNPs increasing in both contrasts) quadrants, while discordant signals appear in the top-left and bottom-right quadrants. Color intensity reflects the number of loci overlapping across both contrasts, sorted by both Δ*p*-direction and −log₁₀(*P*) rank. **(B)** RRHO comparing significance of loci in extinct and rescue contrasts in August relative to unexposed populations, showing negligible concordance between extinct and rescue populations at this early time point. **(C)** RRHO comparing significance of loci in pre-adapted and extinct contrasts in August relative to unexposed populations, showing significant concordance between pre-adapted and extinct populations at this early time point. **(D)** RRHO comparing significance of loci in pre-adapted and rescue contrasts in August relative to unexposed populations, showing negligible concordance between pre-adapted and rescue populations at this early time point.

**Fig. S12.**
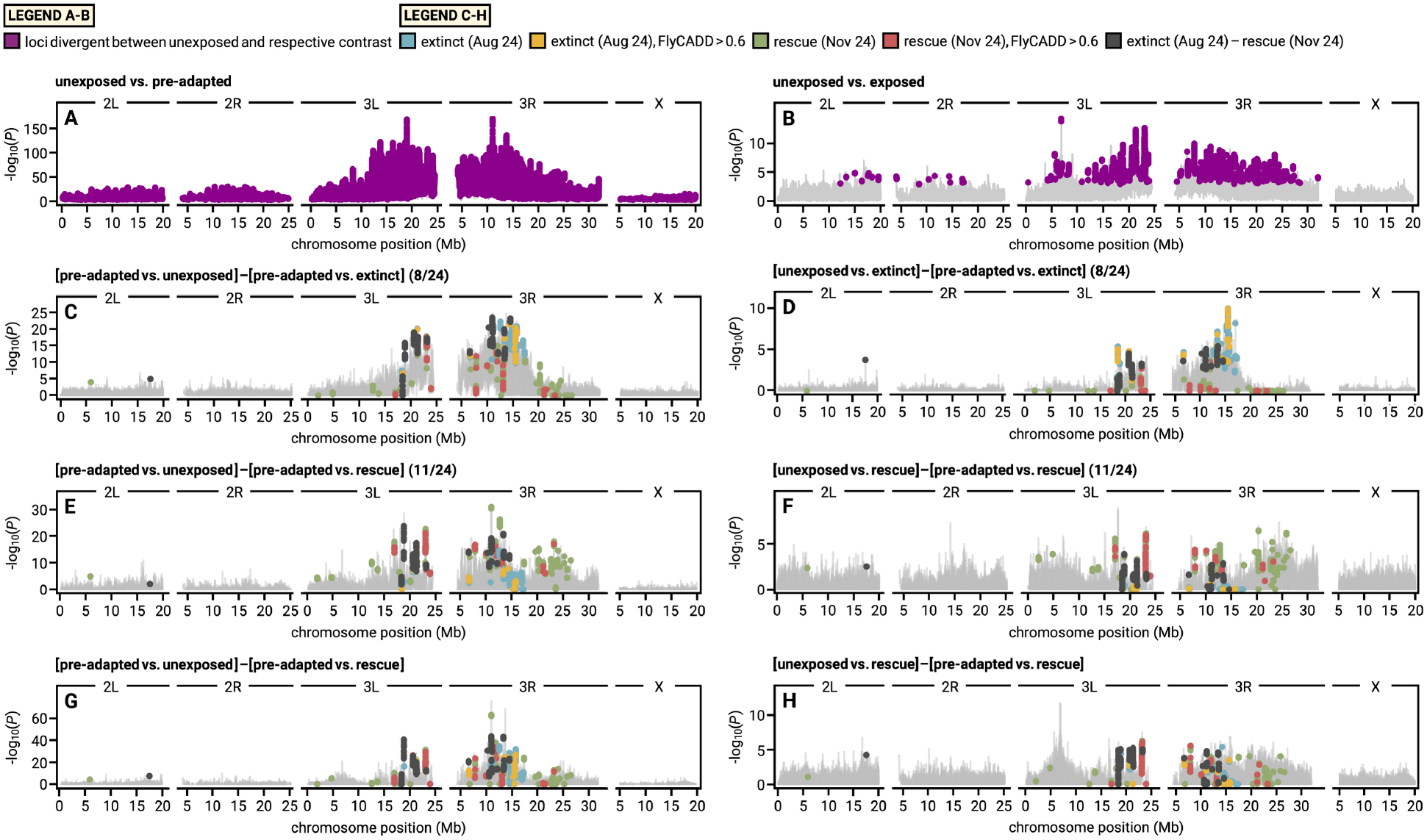
Identification of convergent candidate loci underlying *de novo* and pre-adapted evolutionary rescue via a residual subtraction framework. **(A and B)** Genome-wide allele frequency divergence from unexposed populations across the full time series in pre-adapted **(A)** and exposed **(B)** populations, shown as −log₁₀(*P*) from generalized linear models in 21-SNP sliding windows. Purple points denote loci divergent from unexposed populations in each contrast (FDR < 0.01). **(C to H)** Residual Manhattan plots isolating convergent candidate loci by subtracting treatment-specific background signals—including those driven by genomic drift—to highlight alleles exhibiting temporal convergent selection responses across spinosad-exposed treatments. In all panels C–H, colored points denote empirical outlier SNPs from extinct (August 24, cyan), extinct with FlyCADD > 0.6 (August 24, yellow), rescue (November 24, green), rescue with FlyCADD > 0.6 (November 24, red), and SNPs identified in both extinct (August 24) and rescue (November 24) analyses (dark). Panels C and D show residual contrasts in August: **(C)** isolates pre-adaptation-specific divergence after removing signals shared with extinct populations; **(D)** isolates divergence in extinct populations after removing the pre-adapted background signal. Panels E and F show equivalent contrasts in November: **(E)** isolates pre-adaptation-specific divergence after removing signals shared with rescue populations; **(F)** isolates rescue-specific divergence after removing the pre-adapted background signal. Panels G and H show the same contrasts as E and F respectively, integrated across the full experimental time series. Candidate outlier SNPs are overlaid on all residual panels to facilitate visual comparison of convergent loci across analytical frameworks and time points.

**Fig. S13.**
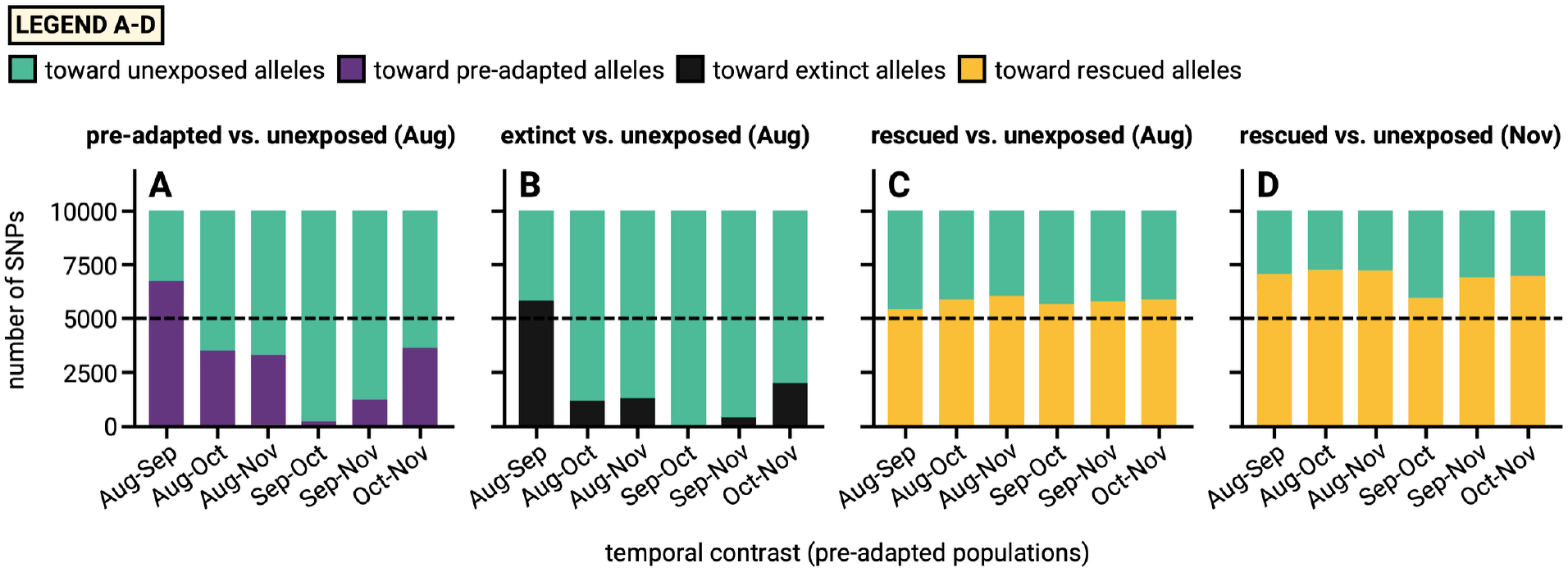
Direction of allele frequency change at top GLM outlier SNPs in pre-adapted populations. For the top 10,000 SNPs (lowest GLM FDR) in each focal contrast, bars show the number of SNPs whose allele frequency in pre-adapted populations shifted in the direction of allelic change of the exposed group in that contrast (pre-adapted, extinct or rescue) or in the direction of unexposed allelic change, during each temporal interval. Focal contrasts are (**A**) ‘unexposed vs. pre-adapted’, August; (**B**) ‘unexposed vs. extinct’, August; (**C**) ‘unexposed vs. rescue’, August; and (**D**) ‘unexposed vs. rescue’, November. SNPs were restricted to those with mean allele frequency between 0.15 and 0.85 (n = 9,977–10,000 per set). Dashed lines mark half of the SNPs in each set. Counts are not compared with background SNPs; see Table S13 for tests relative to matched background SNPs.

**Fig. S14.**
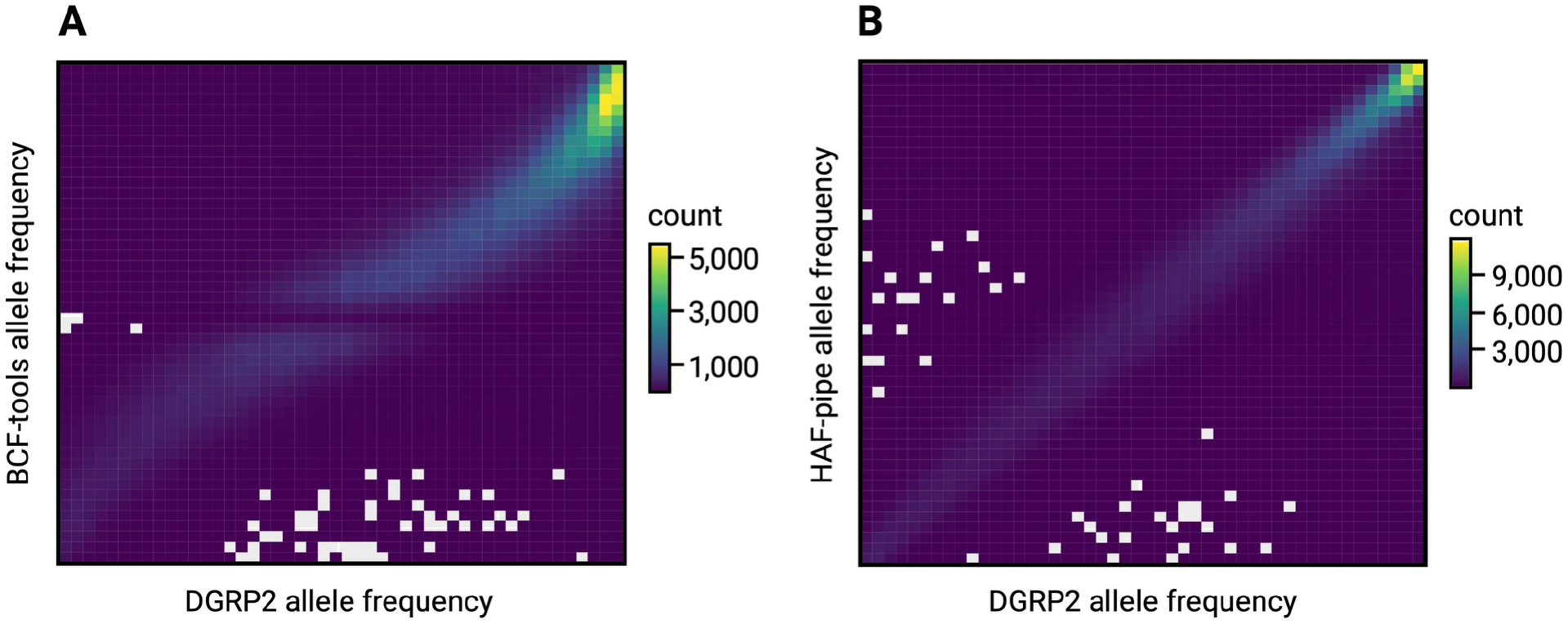
Comparison of raw and HAF-pipe-imputed allele frequency estimates. Mean reference-allele frequencies estimated from pooled sequencing data using (**A**) raw read counts (*BCFtools*) or (**B**) haplotype-informed inference (HAF-pipe) are plotted against mean reference-allele frequencies calculated directly from DGRP2 founder genotypes, for SNPs present in all three sources. Both approaches were strongly correlated with founder frequencies, but the correlation was higher for HAF-pipe-imputed frequencies (Pearson’s R = 0.90) than for *BCFtools*-derived frequencies (R = 0.86). Two-dimensional density plots show that HAF-pipe estimates more closely follow the 1:1 relationship with founder frequencies, whereas *BCFtools* estimates are more dispersed. HAF-pipe allele frequency estimates were therefore used for all downstream genomic analyses.

**Table S1.** Overlap between parallel temporal allele frequency change in unexposed populations and August pre-adapted outliers. At each SNP, the direction of pre-adapted allelic change was defined from the August ‘unexposed vs. pre-adapted’ contrast as the direction in which pre-adapted populations differed from unexposed populations, and the direction of unexposed allelic change as the opposite direction. For each temporal interval, allele frequency change in each unexposed population was assigned a sign so that positive values indicate change in the direction of pre-adapted allelic change. SNPs were classified as changing in parallel in the direction of pre-adapted allelic change (‘pre-adapted’) if all, or all but one, populations changed by more than 0.03 in that direction, and in the direction of unexposed allelic change (‘unexposed’) if all, or all but one, populations changed by more than 0.03 in the opposite direction. Hypergeometric tests assess whether these SNPs (‘temporal outliers’) overlap with SNPs differentiated between unexposed and pre-adapted populations in August (‘PAvE.T1 outliers’; GLM FDR < 0.05) more than expected by chance; ‘non-temporal outliers’ gives the number of remaining SNPs. Nine unexposed populations were included (eight for intervals involving September). Time points are T1 = August 24, T2 = September 30, T3 = October 27, and T4 = November 24, 2023.

| **contrast** | **direction** | **overlap** | **temporal outliers** | **non-temporal outliers** | **PAvE.T1 outliers** | **hypergeometric *P*-value** |
|---|---|---|---|---|---|---|
| T1 vs. T2 | pre-adapted | 240 | 656 | 1095183 | 455515 | 0.99591688 |
| T1 vs. T3 | pre-adapted | 64 | 188 | 1095651 | 455515 | 0.98564203 |
| T1 vs. T4 | pre-adapted | 253 | 716 | 1095123 | 455515 | 0.99972041 |
| T2 vs. T3 | pre-adapted | 91 | 623 | 1095216 | 455515 | 1 |
| T2 vs. T4 | pre-adapted | 564 | 2274 | 1093565 | 455515 | 1 |
| T3 vs. T4 | pre-adapted | 84 | 340 | 1095499 | 455515 | 1 |
| T1 vs. T2 | unexposed | 226 | 558 | 1095281 | 455515 | 0.7096114 |
| T1 vs. T3 | unexposed | 120 | 207 | 1095632 | 455515 | 1.47E-06 |
| T1 vs. T4 | unexposed | 817 | 1190 | 1094649 | 455515 | 2.16E-79 |
| T2 vs. T3 | unexposed | 547 | 1247 | 1094592 | 455515 | 0.05304272 |
| T2 vs. T4 | unexposed | 1343 | 2703 | 1093136 | 455515 | 9.84E-18 |
| T3 vs. T4 | unexposed | 169 | 386 | 1095453 | 455515 | 0.20263766 |

**Table S2.** Candidate loci for spinosad resistance identified through a literature search. Each row represents a gene or genomic feature previously implicated in spinosad resistance, with FlyBase gene identifier (ID), CG annotation symbol, chromosomal location (chromosome, start, and stop positions in bp), gene symbol, and the primary literature source (DOI) from which the candidate was identified. Candidates were drawn from knockout and RNAi screens, differential expression under insecticide exposure, heterologous expression assays, and known involvement in spinosad’s mode of action, and span target-site, detoxification, transport, and cuticular pathways. This candidate list was used to test whether previously characterized resistance loci showed elevated allele frequency divergence in spinosad-exposed populations relative to the genomic background (see *Materials and Methods*).

| **ID** | **CG SYMBOL** | **CHROM** | **START** | **STOP** | **SYMBOL** | **DOI** |
|---|---|---|---|---|---|---|
| FBal0208935 |  |  |  |  |  | 10.1111/1744-7917.12498 |
| FBgn0000036 | CG5610 | 3R | 24393896 | 24457157 | nAChRalpha1 | 10.1371/journal.pone.0103689 |
| FBgn0000038 | CG11348 | 3L | 4429349 | 4435993 | nAChRbeta1 | 10.1371/journal.pone.0103689 |
| FBgn0000326 | CG9858 | 2R | 21617061 | 21619079 | clt | 10.1038/s41598-024-70037-x |
| FBgn0001230 | CG5436 | 3R | 24055080 | 24057310 | Hsp68 | 10.1038/s41598-024-70037-x |
| FBgn0002626 | CG7939 | 3R | 30045229 | 30046161 | RpL32 | 10.1371/journal.pone.0103689 |
| FBgn0003462 | CG11793 | 3L | 11112281 | 11113740 | Sod1 | 10.7554/eLife.73812 |
| FBgn0003515 | CG4314 | 3L | 16497651 | 16500460 | st | 10.1098/rspb.2022.0625 |
| FBgn0004118 | CG6798 | 3R | 24509228 | 24515106 | nAChRbeta2 | 10.1371/journal.pone.0103689 |
| FBgn0004240 | CG12763 | 2R | 18865765 | 18866260 | DptA | 10.1016/j.pestbp.2021.104852 |
| FBgn0004512 | CG3879 | 2R | 12940323 | 12946347 | Mdr49 | 10.1098/rspb.2022.0625 |
| FBgn0004513 | CG10181 | 3L | 6240500 | 6245746 | Mdr65 | 10.1098/rspb.2022.0625 |
| FBgn0010019 | CG3972 | X | 467057 | 469335 | Cyp4g1 | 10.1371/journal.pone.0103689 |
| FBgn0010213 | CG8905 | 2R | 16772785 | 16773915 | Sod2 | 10.7554/eLife.73812 |
| FBgn0010226 | CG8938 | 2R | 17093246 | 17099501 | GstS1 | 10.1371/journal.pone.0103689 |
| FBgn0010241 | CG8523 | 2R | 14246966 | 14252597 | Mdr50 | 10.1098/rspb.2022.0625 |
| FBgn0011244 | CG2830 | 2L | 728362 | 730564 | Hsp60B | 10.1038/s41598-024-70037-x |
| FBgn0011293 | CG6642 | 3L | 17023123 | 17023741 | a10 | 10.1016/j.pestbp.2021.104852 |
| FBgn0013275 | CG31366 | 3R | 11954163 | 11956544 | Hsp70Aa | 10.1038/s41598-024-70037-x |
| FBgn0013772 | CG10248 | 2R | 14887171 | 14889010 | Cyp6a8 | 10.1038/s41598-024-70037-x |
| FBgn0015033 | CG4321 | 3L | 7493557 | 7496197 | Cyp4d8 | 10.1038/s41598-024-70037-x |
| FBgn0015035 | CG4105 | 2L | 9747839 | 9750071 | Cyp4e3 | 10.1111/1744-7917.12498 |
| FBgn0015519 | CG2302 | X | 8276200 | 8367169 | nAChRalpha3 | 10.1371/journal.pone.0103689 |
| FBgn0015568 | CG1031 | 3R | 7545733 | 7549124 | alpha-Est1 | 10.1371/journal.pone.0103689 |
| FBgn0023541 | CG3540 | X | 2173415 | 2175315 | Cyp4d14 | 10.1038/s41598-024-70037-x |
| FBgn0025454 | CG8453 | 2R | 12185667 | 12188431 | Cyp6g1 | 10.1371/journal.pone.0103689 |
| FBgn0028871 | CG3474 | 2L | 14424315 | 14425189 | Cpr35B | 10.1038/s41598-024-70037-x |
| FBgn0031109 | CG11748 | X | 20415744 | 20416521 | Obp19a | 10.1016/j.pestbp.2021.104852 |
| FBgn0031111 | CG15457 | X | 20417921 | 20418964 | Obp19c | 10.1016/j.pestbp.2021.104852 |
| FBgn0032151 | CG4128 | 2L | 9793317 | 9886250 | nAChRalpha6 | 10.7554/eLife.73812 |
| FBgn0033065 | CG8345 | 2R | 6173684 | 6176195 | Cyp6w1 | 10.1038/s41598-024-70037-x |
| FBgn0033294 | CG8693 | 2R | 8458815 | 8460884 | Mal-A4 | 10.1016/j.pestbp.2021.104852 |
| FBgn0033302 | CG8687 | 2R | 8564187 | 8567202 | Cyp6a14 | 10.1111/1744-7917.12498 |
| FBgn0033395 | CG1944 | 2R | 9237512 | 9239831 | Cyp4p2 | 10.1111/1744-7917.12498 |
| FBgn0033524 | CG18377 | 2R | 10449350 | 10458892 | Cyp49a1 | 10.1111/1744-7917.12498 |
| FBgn0033728 | CG8505 | 2R | 12402216 | 12405108 | Cpr49Ae | 10.1111/1744-7917.12498 |
| FBgn0035907 | CG6662 | 3L | 8523742 | 8524866 | GstO1 | 10.1038/s41598-024-70037-x |
| FBgn0037224 | CG14639 | 3R | 4247020 | 4248313 | TwdlF | 10.1038/s41598-024-70037-x |
| FBgn0037225 | CG14643 | 3R | 4248723 | 4251341 | TwdlG | 10.1038/s41598-024-70037-x |
| FBgn0037227 | CG14640 | 3R | 4254708 | 4255833 | TwdlV | 10.1038/s41598-024-70037-x |
| FBgn0037324 | CG10609 | 3R | 5407183 | 5412406 | Orco | 10.1016/j.pestbp.2021.104852 |
| FBgn0038194 | CG3050 | 3R | 14029228 | 14031483 | Cyp6d5 | 10.1111/1744-7917.12498 |
| FBgn0038226 |  |  |  |  |  | 10.1016/j.ibmb.2015.01.018 |
| FBgn0038740 | CG4562 | 3R | 19865417 | 19875678 | Mrp5 | 10.1098/rspb.2022.0625 |
| FBgn0038819 | CG5494 | 3R | 20669571 | 20672821 | Cpr92F | 10.1111/1744-7917.12498 |
| FBgn0039006 | CG12800 | 3R | 22698319 | 22700307 | Cyp6d4 | 10.1371/journal.pone.0103689 |
| FBgn0039685 | CG7592 | 3R | 29716254 | 29716896 | Obp99b | 10.1016/j.pestbp.2021.104852 |
| FBgn0039805 | CG12045 | 3R | 30867521 | 30869184 | Cpr100A | 10.1111/1744-7917.12498 |
| FBgn0041156 | CG8254 | 3L | 7948186 | 7951563 | exex | 10.1016/j.pestbp.2021.104852 |
| FBgn0041579 | CG4740 | 2R | 13393705 | 13394667 | AttC | 10.1016/j.pestbp.2021.104852 |
| FBgn0042206 | CG18548 | 3R | 12364776 | 12365735 | GstD10 | 10.1111/1744-7917.12498 |
| FBgn0043578 | CG9681 | 3L | 16727299 | 16727989 | PGRP-SB1 | 10.1016/j.pestbp.2021.104852 |
| FBgn0050296 | CG30296 | 2R | 21017784 | 21024918 | RIC-3 | 10.1016/j.ibmb.2009.11.004 |
| FBgn0050489 | CG30489 | 2R | 11117225 | 11121884 | Cyp12d1-p | 10.1111/1744-7917.12498 |
| FBgn0052029 | CG32029 | 3L | 8640439 | 8643584 | Cpr66D | 10.1038/s41598-024-70037-x |
| FBgn0052091 | CG32091 | 3L | 11631392 | 11642902 | CG32091 | 10.1098/rspb.2022.0625 |
| FBgn0063496 | CG17525 | 2R | 18403248 | 18404086 | GstE4 | 10.1016/j.pestbp.2021.104852 |
| FBgn0063497 | CG17524 | 2R | 18400713 | 18401467 | GstE3 | 10.1038/s41598-024-70037-x |
| FBgn0067903 | CG33706 | 2R | 23600914 | 23601773 | Pai | 10.1016/j.pestbp.2021.104852 |
| FBgn0086778 | CG32538 | X | 19327075 | 19348447 | nAChRalpha7 | 10.1038/s41598-024-70037-x |
| FBgn0261930 | CG6172 | X | 582121 | 598151 | vnd | 10.1016/j.pestbp.2021.104852 |
| FBgn0285944 | CG9907 | X | 16455027 | 16533096 | para | 10.1038/s41598-024-70037-x |
| FBgn0286832 | - | - | - | - | Actin | 10.1371/journal.pone.0103689 |
| FBgn0286902 | - | - | - | - | Mdoa\nAChRalpha6 | 10.1111/j.1365-2583.2007.00730.x |
| FBgn0288229 | CG3164 | 2L | 122624 | 130791 | CG3164 | 10.1098/rspb.2022.0625 |
| FBgn0289113 | - | - | - | - | Mdoa\CYP6G4 | 10.1371/journal.pone.0103689 |
| FBrf0249385 |  |  |  |  |  | 10.1002/ps.6290 |
| FBgn0010100 | CG9244 | 2L | 21168570 | 21173185 | mAcon1 | 10.7554/eLife.73812 |
| FBgn0037862 | CG4706 | 3R | 11137177 | 11139742 | mAcon2 | 10.7554/eLife.73812 |

**Table S3.**
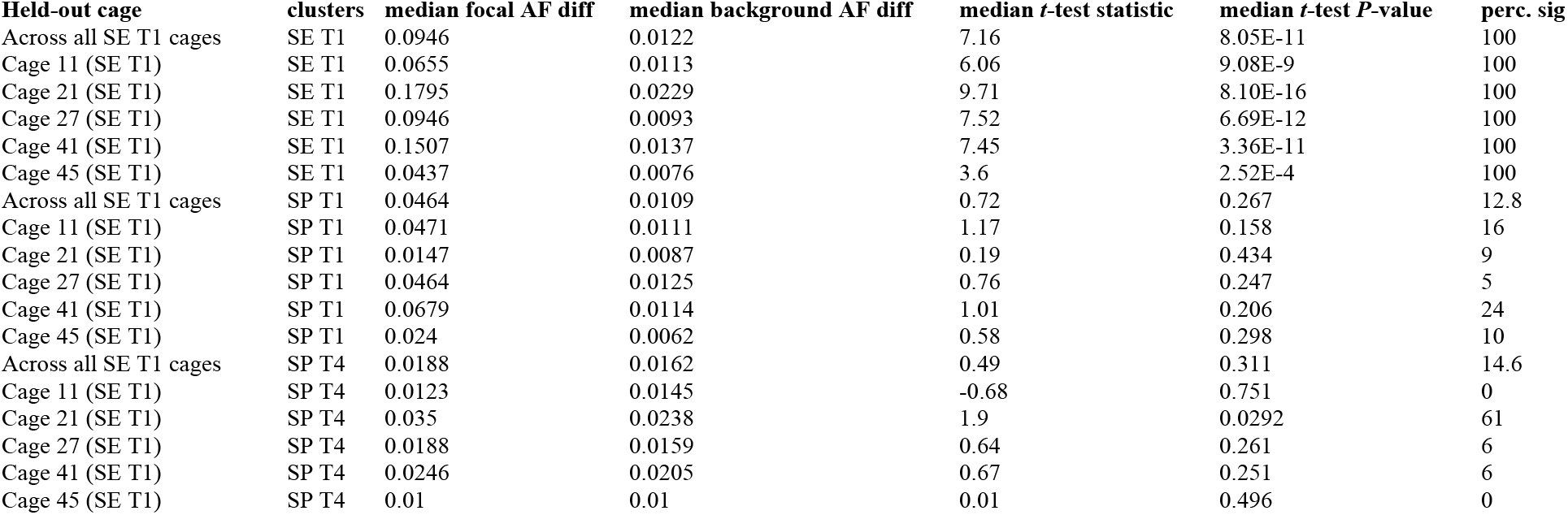
Leave-one-out parallelism tests for extinct populations in August. One-tailed Welch *t*-tests comparing allele frequency differences between unexposed populations and individual extinct (‘left out’) populations in August at top SNPs (one per cluster) against 100 sets of random matched non-outlier SNPs. For extinct-population August clusters, each extinct replicate was left out in turn, and SNPs and the direction of divergence were identified from GLMs of the four remaining replicates, so SNP discovery was independent of the replicate tested. For comparison across contrasts, the same extinct replicates were also tested at SNPs identified from all-sample GLMs of the ‘unexposed vs. rescue’ contrasts in August and November (rescue-population August and November clusters, respectively). Rows beginning ‘Across all’ report medians pooled across all 500 tests (100 iterations × 5 replicates). Mesocosm labels are given in parentheses. Abbreviations: SE = extinct, SP = rescue; T1 = August 24 and T4 = November 24, 2023. ‘perc. sig’ denotes the percentage of the 100 matched-set iterations yielding P < 0.05.

**Table S4.** Leave-one-out parallelism tests for rescue populations in August. One-tailed Welch *t*-tests comparing allele frequency differences between unexposed populations and individual rescue (‘left out’) populations in August at top SNPs (one per cluster) against 100 sets of random matched non-outlier SNPs. For rescue-population August clusters, each rescue replicate was left out in turn, and SNPs and the direction of divergence were identified from GLMs of the four remaining replicates in August, so SNP discovery was independent of the replicate tested. For comparison across contrasts, the same rescue replicates were also tested at SNPs identified from all-sample GLMs of the ‘unexposed vs. extinct’ contrast in August and the ‘unexposed vs. rescue’ contrast in November (extinct-population August and rescue-population November clusters, respectively). Rows beginning ‘Across all’ report medians pooled across all 500 tests (100 iterations × 5 replicates). Mesocosm labels are given in parentheses. Abbreviations: SE = extinct, SP = rescue; T1 = August 24 and T4 = November 24, 2023. ‘perc. sig’ denotes the percentage of the 100 matched-set iterations yielding P < 0.05.

| **Held-out cage** | **clusters** | **median focal AF diff** | **median background AF diff** | **median *t*-test statistic** | **median *t*-test *P*-value** | **perc. sig** |
|---|---|---|---|---|---|---|
| Across all SP T1 cages | SE T1 | 0.011 | 0.0091 | 0.34 | 0.368 | 12.4 |
| Cage 3 (SP T1) | SE T1 | 0.0141 | 0.0099 | 0.3 | 0.382 | 5 |
| Cage 7 (SP T1) | SE T1 | 0.011 | 0.0104 | 0.06 | 0.474 | 4 |
| Cage 15 (SP T1) | SE T1 | 0.0092 | 0.0055 | 0.29 | 0.385 | 4 |
| Cage 33 (SP T1) | SE T1 | 0.0252 | 0.0051 | 1.61 | 0.0556 | 48 |
| Cage 37 (SP T1) | SE T1 | 0.009 | 0.0159 | -0.65 | 0.742 | 1 |
| Across all SP T1 cages | SP T1 | 0.0381 | 0.0067 | 0.91 | 0.201 | 5.5 |
| Cage 3 (SP T1) | SP T1 | NA | NA | NA | NA | NA |
| Cage 7 (SP T1) | SP T1 | 0.0794 | 0.0112 | 1.13 | 0.171 | 0 |
| Cage 15 (SP T1) | SP T1 | 0.0438 | 0.0017 | 0.69 | 0.258 | 6 |
| Cage 33 (SP T1) | SP T1 | 0.0324 | 0.0049 | 1.01 | 0.165 | 14 |
| Cage 37 (SP T1) | SP T1 | 0.0199 | 0.0068 | 0.53 | 0.3 | 2 |
| Across all SP T1 cages | SP T4 | 0.0087 | 0.0076 | 0.47 | 0.319 | 23.6 |
| Cage 3 (SP T1) | SP T4 | 0.0205 | 0.0082 | 2.27 | 0.0117 | 82 |
| Cage 7 (SP T1) | SP T4 | 0.0064 | 0.01 | -0.93 | 0.823 | 0 |
| Cage 15 (SP T1) | SP T4 | 0.0116 | 0.0078 | 0.65 | 0.259 | 16 |
| Cage 33 (SP T1) | SP T4 | 0.0087 | 0.0049 | 0.87 | 0.192 | 19 |
| Cage 37 (SP T1) | SP T4 | 0.0069 | 0.007 | -0.04 | 0.515 | 1 |

**Table S5.**
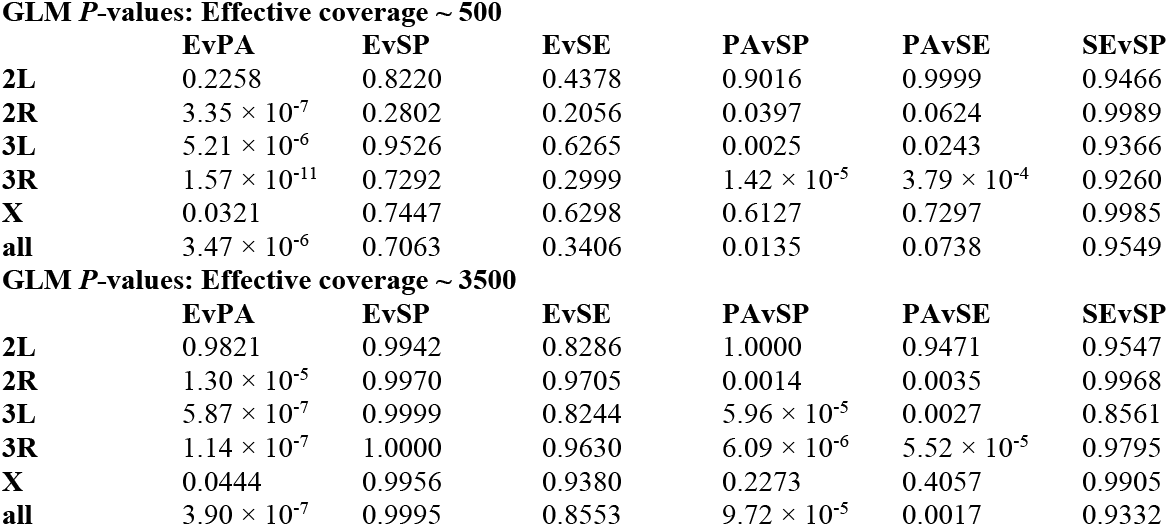
Comparisons of genetic diversity among treatments using Watterson’s θ in August. Generalized linear model *P*-values testing for differences in Watterson’s θ among experimental treatments at the first post-exposure time point (August 24, 2023). Estimates were calculated using HAF-pipe-imputed allele frequencies and evaluated at both chromosome-specific (2L, 2R, 3L, 3R, X) and genome-wide (‘all’) levels. To assess sensitivity to assumptions about sequencing depth, θ was estimated under two effective coverage values (approximately 500× and 3,500×), and GLMs were run separately for each assumption. Abbreviations: E = unexposed, PA = pre-adapted, SE = extinct, SP = rescue.

**Table S6.** Leave-one-out parallelism tests for rescue populations in November. One-tailed Welch *t*-tests comparing allele frequency differences between unexposed populations and individual rescue (‘left out’) populations in November at top SNPs (one per cluster) against 100 sets of random matched non-outlier SNPs. For rescue-population November clusters, each rescue replicate was left out in turn, and SNPs and the direction of divergence were identified from GLMs of the four remaining replicates in November, so SNP discovery was independent of the replicate tested. For comparison across contrasts, the same rescue replicates were also tested at SNPs identified from all-sample GLMs of the ‘unexposed vs. extinct’ and ‘unexposed vs. rescue’ contrasts in August (extinct-population and rescue-population August clusters, respectively). Rows beginning ‘Across all’ report medians pooled across all 500 tests (100 iterations × 5 replicates). Mesocosm labels are given in parentheses. Abbreviations: SE = extinct, SP = rescue; T1 = August 24 and T4 = November 24, 2023. ‘perc. sig’ denotes the percentage of the 100 matched-set iterations yielding P < 0.05.

| **Held-out cage** | **clusters** | **median focal AF diff** | **median background AF diff** | **median *t*-test statistic** | **median *t*-test *P*-value** | **perc. sig** |
|---|---|---|---|---|---|---|
| Across all SP T4 cages | SE T1 | 0.0287 | 0.0155 | 0.46 | 0.322 | 5.6 |
| Cage 3 (SP T4) | SE T1 | 0.0307 | 0.0057 | 1.16 | 0.125 | 16 |
| Cage 7 (SP T4) | SE T1 | 0.0334 | 0.0182 | 0.88 | 0.191 | 6 |
| Cage 15 (SP T4) | SE T1 | -0.0112 | 0.0082 | -0.96 | 0.83 | 0 |
| Cage 33 (SP T4) | SE T1 | 0.0281 | 0.0156 | 0.69 | 0.245 | 5 |
| Cage 37 (SP T4) | SE T1 | 0.0287 | 0.0286 | 0 | 0.499 | 1 |
| Across all SP T4 cages | SP T1 | 0.0031 | 0.0077 | -0.45 | 0.656 | 2.4 |
| Cage 3 (SP T4) | SP T1 | 0.0472 | 0.0118 | 0.36 | 0.37 | 0 |
| Cage 7 (SP T4) | SP T1 | 0.0309 | 0.0006 | 0.55 | 0.314 | 8 |
| Cage 15 (SP T4) | SP T1 | -0.1664 | 0.0072 | -3.14 | 0.972 | 0 |
| Cage 33 (SP T4) | SP T1 | 0.0031 | 0.0053 | -0.05 | 0.519 | 4 |
| Cage 37 (SP T4) | SP T1 | -0.1428 | 0.0214 | -1.03 | 0.808 | 0 |
| Across all SP T4 cages | SP T4 | 0.0587 | 0.0145 | 4.25 | 1.34E-5 | 98.4 |
| Cage 3 (SP T4) | SP T4 | 0.0364 | 0.0085 | 3.13 | 9.39E-4 | 96 |
| Cage 7 (SP T4) | SP T4 | 0.0547 | 0.0148 | 4.7 | 1.76E-6 | 100 |
| Cage 15 (SP T4) | SP T4 | 0.0608 | 0.0117 | 4.54 | 3.70E-6 | 100 |
| Cage 33 (SP T4) | SP T4 | 0.0713 | 0.0165 | 8.02 | 4.98E-15 | 100 |
| Cage 37 (SP T4) | SP T4 | 0.0587 | 0.0186 | 2.77 | 0.00295 | 96 |

**Table S7.** Genomic features associated with convergent candidate loci in rescue populations in November. Summary of features within 5 kb of convergent candidate SNPs identified in exposed populations that persisted, at the final experimental time point (November 24). Outlier loci were defined as SNPs significant in the ‘unexposed vs. rescue’ contrast in November (FDR < 0.05) and in the ‘unexposed vs. pre-adapted’ contrasts in both August and November (FDR < 0.01), with unexposed allele frequencies of 0.1–0.9 and an allele frequency difference > 0.1 between unexposed and pre-adapted populations in August. FlyCADD scores were used to prioritize variants with predicted functional effects; ‘YES’ indicates features containing or flanking at least one candidate SNP with a FlyCADD score > 0.60. FlyBase IDs marked with an asterisk contain candidate SNPs directly, whereas unmarked entries lie within 5 kb upstream or downstream flanking regions.

| **ID** | **SYMBOL** | **TYPE** | **CHROM** | **START** | **STOP** | **NAME** | **GENE SYMBOL** | **FLYCADD > 0.60** |
|---|---|---|---|---|---|---|---|---|
| FBgn0000003 | CR32864 | SRP_RNA_gene | 3R | 6822500 | 6822798 | Signal recognition particle 7SL RNA CR32864 | 7SLRNA:CR32864 | NO |
| FBgn0000551 | CG7673 | protein_coding_gene | 3L | 21296763 | 21297803 | Ecdysone-dependent gene 78E | Edg78E | NO |
| FBgn0000568* | CG8127 | protein_coding_gene | 3L | 17950953 | 18064696 | Ecdysone-induced protein 75B | Eip75B | NO |
| FBgn0001128 | CG9042 | protein_coding_gene | 2L | 5943643 | 5948968 | Glycerol-3-phosphate dehydrogenase 1 | Gpdh1 | NO |
| FBgn0001230* | CG5436 | protein_coding_gene | 3R | 24055080 | 24057310 | Heat shock protein 68 | Hsp68 | NO |
| FBgn0001323* | CG4761 | protein_coding_gene | 3L | 20597141 | 20620659 | knirps-like | knrl | YES |
| FBgn0002941 | CG6534 | protein_coding_gene | 3R | 21548661 | 21557182 | slouch | slou | YES |
| FBgn0003042* | CG32443 | protein_coding_gene | 3L | 21306138 | 21318020 | Polycomb | Pc | NO |
| FBgn0003339 | CG1030 | protein_coding_gene | 3R | 6823120 | 6849981 | Sex combs reduced | Scr | NO |
| FBgn0004841* | CG6515 | protein_coding_gene | 3R | 10737592 | 10751228 | Tachykinin-like receptor at 86C | TkR86C | NO |
| FBgn0004882 | CG10868 | protein_coding_gene | 3R | 23260610 | 23280856 | oo18 RNA-binding protein | orb | NO |
| FBgn0004901* | CG2867 | protein_coding_gene | 3R | 7909598 | 7911779 | Phosphoribosylamidotransferase | Prat | YES |
| FBgn0005674* | CG5394 | protein_coding_gene | 3R | 23952136 | 23958391 | Glutamyl-prolyl-tRNA synthetase | GluProRS | NO |
| FBgn0010424 | CG7930 | protein_coding_gene | 3L | 17046063 | 17048390 | Troponin C at 73F | TpnC73F | YES |
| FBgn0011904 | CR31486 | tRNA_gene | 3R | 6822963 | 6823035 | transfer RNA:Lysine-TTT 2-4 | tRNA:Lys-TTT-2-4 | NO |
| FBgn0014930 | CG2846 | protein_coding_gene | 3R | 7912017 | 7913290 | Riboflavin kinase | Rfk | YES |
| FBgn0014931 | CG2678 | protein_coding_gene | 3R | 7913851 | 7916550 | kipferl | kipf | YES |
| FBgn0015381 | CG9019 | protein_coding_gene | 2L | 5925937 | 5939261 | dissatisfaction | dsf | NO |
| FBgn0015919 | CG10605 | protein_coding_gene | 3L | 12609631 | 12621798 | caupolican | caup | NO |
| FBgn0024733* | CG17521 | protein_coding_gene | 3L | 23093190 | 23095471 | Ribosomal protein L10 | RpL10 | NO |
| FBgn0024909* | CG2670 | protein_coding_gene | 3R | 7907333 | 7909721 | TBP-associated factor 7 | Taf7 | YES |
| FBgn0025781* | CG6759 | protein_coding_gene | 3R | 23281201 | 23284123 | Cell division cycle 16 | Cdc16 | NO |
| FBgn0025807 | CG3945 | protein_coding_gene | 3L | 18870045 | 18872553 | Rad9 | Rad9 | YES |
| FBgn0026162 | CR31490 | tRNA_gene | 3R | 6819894 | 6819966 | transfer RNA:Lysine-TTT 2-1 | tRNA:Lys-TTT-2-1 | NO |
| FBgn0026163 | CR31487 | tRNA_gene | 3R | 6821068 | 6821140 | transfer RNA:Lysine-TTT 2-2 | tRNA:Lys-TTT-2-2 | NO |
| FBgn0026164 | CR31489 | tRNA_gene | 3R | 6821789 | 6821861 | transfer RNA:Lysine-TTT 2-3 | tRNA:Lys-TTT-2-3 | NO |
| FBgn0026576 | CG5991 | protein_coding_gene | 3R | 24050783 | 24053592 | Phosphatidylserine decarboxylase | Pisd | NO |
| FBgn0027085 | CG7479 | protein_coding_gene | 3L | 4825853 | 4829038 | Leucyl-tRNA synthetase, mitochondrial | LeuRS-m | NO |
| FBgn0027090 | CG10506 | protein_coding_gene | 3R | 25263267 | 25265721 | Glutaminyl-tRNA synthetase | GlnRS | NO |
| FBgn0035577 | CG13708 | protein_coding_gene | 3L | 4829092 | 4833591 | - | CG13708 | NO |
| FBgn0036695 | CG7853 | protein_coding_gene | 3L | 17042797 | 17044465 | PAPS transporter 2 | Papst2 | NO |
| FBgn0036696 | CG14057 | protein_coding_gene | 3L | 17044401 | 17045038 | POP5 ribonuclease P/MRP subunit | Pop5 | YES |
| FBgn0036697* | CG7725 | protein_coding_gene | 3L | 17049512 | 17055454 | rogdi | rogdi | YES |
| FBgn0036698 | CG7724 | protein_coding_gene | 3L | 17055613 | 17058533 | - | CG7724 | NO |
| FBgn0036817 | CG6865 | protein_coding_gene | 3L | 18863647 | 18865514 | - | CG6865 | YES |
| FBgn0036818* | CG14074 | protein_coding_gene | 3L | 18866225 | 18867431 | - | CG14074 | YES |
| FBgn0036819 | CG6856 | protein_coding_gene | 3L | 18867771 | 18869219 | Dysbindin | Dysb | YES |
| FBgn0036820 | CG6852 | protein_coding_gene | 3L | 18869230 | 18870057 | Glutaredoxin 1 | Grx1 | YES |
| FBgn0037014 | CG13251 | protein_coding_gene | 3L | 20637199 | 20640910 | - | CG13251 | YES |
| FBgn0037068 | CG7663 | protein_coding_gene | 3L | 21294492 | 21296698 | Cuticular protein 78Cb | Cpr78Cb | NO |
| FBgn0037069* | CG7658 | protein_coding_gene | 3L | 21300580 | 21301171 | Cuticular protein 78Cc | Cpr78Cc | NO |
| FBgn0037070* | CG11309 | protein_coding_gene | 3L | 21301785 | 21304658 | - | CG11309 | NO |
| FBgn0037071* | CG7632 | protein_coding_gene | 3L | 21304697 | 21306197 | - | CG7632 | NO |
| FBgn0037073* | CG7338 | protein_coding_gene | 3L | 21335215 | 21338232 | Tsr1 ribosome assembly factor | Tsr1 | YES |
| FBgn0037074* | CG7324 | protein_coding_gene | 3L | 21337681 | 21343716 | TBC1 domain family member 8/9 | Tbc1d8-9 | YES |
| FBgn0037076* | CG12972 | protein_coding_gene | 3L | 21344768 | 21349288 | earthbound 2 | ebd2 | YES |
| FBgn0037078 | CG12971 | protein_coding_gene | 3L | 21355100 | 21356495 | - | CG12971 | NO |
| FBgn0037525* | CG17816 | protein_coding_gene | 3R | 7855582 | 7886733 | - | CG17816 | YES |
| FBgn0037529 | CG2957 | protein_coding_gene | 3R | 7889277 | 7902667 | mitochondrial ribosomal protein S9 | mRpS9 | YES |
| FBgn0037530* | CG2943 | protein_coding_gene | 3R | 7903030 | 7907363 | ER membrane protein complex subunit 1 | EMC1 | YES |
| FBgn0037829 | CG14691 | protein_coding_gene | 3R | 10753331 | 10755634 | - | CG14691 | NO |
| FBgn0037956* | CG6959 | protein_coding_gene | 3R | 11822702 | 11869460 | - | CG6959 | YES |
| FBgn0037972 | CG10005 | protein_coding_gene | 3R | 11975156 | 11977197 | - | CG10005 | YES |
| FBgn0037973 | CG18547 | protein_coding_gene | 3R | 11978574 | 11980390 | - | CG18547 | YES |
| FBgn0037974* | CG12224 | protein_coding_gene | 3R | 11980704 | 11982622 | - | CG12224 | YES |
| FBgn0037975 | CG3397 | protein_coding_gene | 3R | 11983098 | 11984541 | - | CG3397 | YES |
| FBgn0037979 | CG3532 | protein_coding_gene | 3R | 12010772 | 12015059 | GRIP and coiled-coil domain containing 185 kDa | GCC185 | NO |
| FBgn0037980* | CG3313 | protein_coding_gene | 3R | 12014943 | 12017031 | DDB1 and CUL4 associated factor 12 | DCAF12 | NO |
| FBgn0037981 | CG3169 | protein_coding_gene | 3R | 12017156 | 12018550 | Spt3 | Spt3 | NO |
| FBgn0037986 | CG14736 | protein_coding_gene | 3R | 12040528 | 12043453 | - | CG14736 | NO |
| FBgn0037987 | CG14739 | protein_coding_gene | 3R | 12048253 | 12049213 | - | CG14739 | NO |
| FBgn0038082 | CG5724 | protein_coding_gene | 3R | 12739642 | 12741417 | UDP-glycosyltransferase family 37 member A2 | Ugt37A2 | NO |
| FBgn0038083 | CG5999 | protein_coding_gene | 3R | 12741958 | 12743680 | UDP-glycosyltransferase family 37 member A3 | Ugt37A3 | NO |
| FBgn0038118* | CG7855 | protein_coding_gene | 3R | 13088651 | 13163876 | timeout | timeout | YES |
| FBgn0038510* | CG14331 | protein_coding_gene | 3R | 17438968 | 17440728 | - | CG14331 | NO |
| FBgn0038511* | CG5873 | protein_coding_gene | 3R | 17425768 | 17446499 | Curly Su | cysu | NO |
| FBgn0038721* | CG16718 | protein_coding_gene | 3R | 19633418 | 19642722 | subdued | subdued | NO |
| FBgn0038722 | CG7360 | protein_coding_gene | 3R | 19645026 | 19646906 | Nucleoporin 58kD | Nup58 | NO |
| FBgn0038783 | CG4367 | protein_coding_gene | 3R | 20303955 | 20304977 | - | CG4367 | NO |
| FBgn0038784* | CG4362 | protein_coding_gene | 3R | 20306216 | 20307537 | - | CG4362 | NO |
| FBgn0038865 | CG10824 | protein_coding_gene | 3R | 21159238 | 21161092 | Common Dpr-interacting protein | cDIP | YES |
| FBgn0039002 | CG17625 | protein_coding_gene | 3R | 22686849 | 22687407 | - | CG17625 | NO |
| FBgn0039003 | CG4917 | protein_coding_gene | 3R | 22687542 | 22690380 | wolfram syndrome 1 | wfs1 | NO |
| FBgn0039004* | CG6958 | protein_coding_gene | 3R | 22690607 | 22694772 | Nucleoporin 133kD | Nup133 | NO |
| FBgn0039033 | CG17241 | protein_coding_gene | 3R | 22987281 | 22988503 | Odorant receptor 94a | Or94a | NO |
| FBgn0039041 | CG13838 | protein_coding_gene | 3R | 23042241 | 23042669 | - | CG13838 | NO |
| FBgn0039042 | CG13837 | protein_coding_gene | 3R | 23043990 | 23044661 | - | CG13837 | NO |
| FBgn0039043 | CG17121 | protein_coding_gene | 3R | 23044664 | 23048214 | - | CG17121 | NO |
| FBgn0039064* | CG4467 | protein_coding_gene | 3R | 23227727 | 23248903 | - | CG4467 | YES |
| FBgn0039069 | CG6763 | protein_coding_gene | 3R | 23284002 | 23286041 | - | CG6763 | NO |
| FBgn0039070 | CG17083 | protein_coding_gene | 3R | 23286093 | 23288051 | wampa | wam | NO |
| FBgn0039078* | CG4374 | protein_coding_gene | 3R | 23408684 | 23472278 | - | CG4374 | NO |
| FBgn0039079 | CG17381 | protein_coding_gene | 3R | 23483863 | 23485500 | Ionotropic receptor 94g | Ir94g | NO |
| FBgn0039091 | CG10182 | protein_coding_gene | 3R | 23587120 | 23589819 | - | CG10182 | NO |
| FBgn0039092 | CG16723 | protein_coding_gene | 3R | 23593625 | 23596273 | - | CG16723 | NO |
| FBgn0039093 | CG10183 | protein_coding_gene | 3R | 23596561 | 23599034 | - | CG10183 | NO |
| FBgn0039132 | CG5864 | protein_coding_gene | 3R | 23958651 | 23962555 | Adaptor Protein complex 1, sigma subunit | AP-1sigma | NO |
| FBgn0039145 | CG6000 | protein_coding_gene | 3R | 24054391 | 24055092 | - | CG6000 | NO |
| FBgn0039188 | CG17785 | protein_coding_gene | 3R | 24356070 | 24357938 | Golgin-84 | Golgin84 | NO |
| FBgn0039189 | CG18528 | protein_coding_gene | 3R | 24358095 | 24360144 | - | CG18528 | NO |
| FBgn0039190 | CG5762 | protein_coding_gene | 3R | 24360705 | 24361683 | - | CG5762 | NO |
| FBgn0039192 | CG17784 | protein_coding_gene | 3R | 24364081 | 24364980 | - | CG17784 | NO |
| FBgn0039193 | CG13613 | protein_coding_gene | 3R | 24365554 | 24366349 | - | CG13613 | NO |
| FBgn0039305 | CG11858 | protein_coding_gene | 3R | 25262741 | 25263292 | - | CG11858 | NO |
| FBgn0039306 | CG11859 | protein_coding_gene | 3R | 25265844 | 25267757 | RIO kinase 2 | RIOK2 | NO |
| FBgn0039307 | CR13656 | pseudogene | 3R | 25268661 | 25269965 | - | CR13656 | NO |
| FBgn0039308 | CG11889 | protein_coding_gene | 3R | 25270209 | 25273267 | - | CG11889 | NO |
| FBgn0039309 | CG11891 | protein_coding_gene | 3R | 25270209 | 25273267 | - | CG11891 | NO |
| FBgn0039431 | CG6490 | protein_coding_gene | 3R | 26470078 | 26535217 | plum | plum | NO |
| FBgn0040283 | CG6057 | protein_coding_gene | 3R | 24058287 | 24062815 | Structural maintenance of chromosomes 1 | SMC1 | NO |
| FBgn0041105 | CG12809 | protein_coding_gene | 3R | 10071117 | 10073761 | nervous fingers 2 | nerfin-2 | YES |
| FBgn0051093 | CG31093 | protein_coding_gene | 3R | 25789819 | 25790471 | - | CG31093 | NO |
| FBgn0051151* | CG31151 | protein_coding_gene | 3R | 22710883 | 22721150 | winged eye | wge | NO |
| FBgn0051225 | CG31225 | protein_coding_gene | 3R | 23481865 | 23483541 | Ionotropic receptor 94f | Ir94f | NO |
| FBgn0051380 | CG31380 | protein_coding_gene | 3R | 25787997 | 25789455 | - | CG31380 | NO |
| FBgn0052350* | CG32350 | protein_coding_gene | 3L | 23147656 | 23150499 | Vacuolar protein sorting 11 | Vps11 | YES |
| FBgn0052436* | CG32436 | protein_coding_gene | 3L | 21344033 | 21355049 | - | CG32436 | YES |
| FBgn0053337* | CG33337 | protein_coding_gene | 3R | 23590141 | 23592769 | - | CG33337 | NO |
| FBgn0053339* | CG33339 | protein_coding_gene | 3R | 24361847 | 24363142 | - | CG33339 | NO |
| FBgn0053340 | CG33340 | protein_coding_gene | 3R | 24363138 | 24364289 | - | CG33340 | NO |
| FBgn0053630 | CG33630 | protein_coding_gene | 3R | 10062624 | 10063327 | - | CG33630 | YES |
| FBgn0053631 | CG33631 | protein_coding_gene | 3R | 10064007 | 10064645 | - | CG33631 | YES |
| FBgn0053783 | CG33783 | protein_coding_gene | 3R | 10066770 | 10067567 | - | CG33783 | YES |
| FBgn0053784* | CG33784 | protein_coding_gene | 3R | 10067905 | 10068572 | - | CG33784 | YES |
| FBgn0058470* | CG40470 | protein_coding_gene | 3L | 23839860 | 23986410 | - | CG40470 | YES |
| FBgn0083974 | CG34138 | protein_coding_gene | 3R | 19634949 | 19635879 | - | CG34138 | NO |
| FBgn0085284 | CG34255 | protein_coding_gene | 3L | 18870224 | 18870897 | Biogenesis of lysosome-related organelles complex 1, subunit 3 | Blos3 | YES |
| FBgn0085288* | CG34259 | protein_coding_gene | 3L | 2026991 | 2028326 | Gustatory receptor-like 62c | Grl62c | NO |
| FBgn0085413* | CG34384 | protein_coding_gene | 3R | 7814361 | 7855392 | - | CG34384 | YES |
| FBgn0086913* | CG34410 | protein_coding_gene | 3L | 21318774 | 21335027 | Rab26 | Rab26 | YES |
| FBgn0259702 | CG42356 | protein_coding_gene | 3L | 2022070 | 2023391 | Gustatory receptor-like 62b | Grl62b | NO |
| FBgn0260004 | CG7000 | protein_coding_gene | 3R | 21165167 | 21168205 | Sensory neuron membrane protein 1 | Snmp1 | NO |
| FBgn0260005 | CG31284 | protein_coding_gene | 3R | 7891331 | 7901147 | water witch | wtrw | YES |
| FBgn0260745 | CG3359 | protein_coding_gene | 3R | 11984862 | 11993315 | midline fasciclin | mfas | YES |
| FBgn0260793* | CG42573 | protein_coding_gene | 3R | 13121951 | 13155687 | 2mit | 2mit | YES |
| FBgn0261055* | CG42603 | protein_coding_gene | 2L | 5942623 | 5943304 | Seminal fluid protein 26Ad | Sfp26Ad | NO |
| FBgn0261503 | CR42651 | lncRNA_gene | 3R | 6808796 | 6816379 | long non-coding RNA:CR42651 | lncRNA:CR42651 | NO |
| FBgn0261504 | CR42652 | SRP_RNA_gene | 3R | 6820131 | 6820429 | Signal recognition particle 7SL RNA CR42652 | 7SLRNA:CR42652 | NO |
| FBgn0261550 | CG42668 | protein_coding_gene | 3R | 20309344 | 20324196 | Oxysterol-binding protein-related protein 8 | Orp8 | NO |
| FBgn0261556* | CG42674 | protein_coding_gene | 3L | 20149614 | 20198317 | - | CG42674 | NO |
| FBgn0261797* | CG7507 | protein_coding_gene | 3L | 4807368 | 4825687 | Dynein heavy chain 64C | Dhc64C | NO |
| FBgn0261993 | CG42811 | protein_coding_gene | 3R | 24366527 | 24367220 | - | CG42811 | NO |
| FBgn0261994 | CG42812 | protein_coding_gene | 3R | 24367360 | 24368164 | - | CG42812 | NO |
| FBgn0262461 | CR43036 | miRNA_gene | 3L | 18860971 | 18861054 | mir-315 stem loop | mir-315 | NO |
| FBgn0262509* | CG43079 | protein_coding_gene | 3L | 22998372 | 23040300 | neuromusculin | nrm | NO |
| FBgn0262582 | CG43122 | protein_coding_gene | 3R | 20252770 | 20303942 | capicua | cic | NO |
| FBgn0263002* | CR43310 | pseudogene | 3R | 25268009 | 25268464 | - | CR43310 | NO |
| FBgn0263289 | CG43398 | protein_coding_gene | 3R | 26536350 | 26604051 | scribble | scrib | NO |
| FBgn0263390* | CG43442 | protein_coding_gene | 3R | 22981459 | 22998528 | Na[+]/H[+] hydrogen antiporter 2 | Nha2 | NO |
| FBgn0263622 | CR43631 | lncRNA_gene | 3R | 25791642 | 25792384 | long non-coding RNA:CR43631 | lncRNA:CR43631 | NO |
| FBgn0263623* | CR43632 | lncRNA_gene | 3R | 25792600 | 25793303 | long non-coding RNA:CR43632 | lncRNA:CR43632 | NO |
| FBgn0263624 | CR43633 | lncRNA_gene | 3R | 25794066 | 25794421 | long non-coding RNA:CR43633 | lncRNA:CR43633 | NO |
| FBgn0263625 | CR43634 | lncRNA_gene | 3R | 25794848 | 25795186 | long non-coding RNA:CR43634 | lncRNA:CR43634 | NO |
| FBgn0263841 | CG43702 | protein_coding_gene | 3L | 21298675 | 21299063 | - | CG43702 | NO |
| FBgn0263986 | CG6969 | protein_coding_gene | 3R | 22694959 | 22697990 | cardinal | cd | NO |
| FBgn0264001* | CG43744 | protein_coding_gene | 3L | 13521907 | 13803400 | bruno 3 | bru3 | NO |
| FBgn0264357* | CG17299 | protein_coding_gene | 3R | 21140739 | 21214269 | SNF4/AMP-activated protein kinase gamma subunit | SNF4Agamma | YES |
| FBgn0264428 | CR43846 | lncRNA_gene | 3R | 22851088 | 22851804 | long non-coding RNA:CR43846 | lncRNA:CR43846 | NO |
| FBgn0264429 | CR43847 | lncRNA_gene | 3R | 25796591 | 25797778 | long non-coding RNA:CR43847 | lncRNA:CR43847 | NO |
| FBgn0264492 | CG17520 | protein_coding_gene | 3L | 23097072 | 23103460 | casein kinase IIalpha | CkIIalpha | NO |
| FBgn0264906 | CR44097 | lncRNA_gene | 3R | 24054391 | 24055092 | long non-coding RNA:CR44097 | lncRNA:CR44097 | NO |
| FBgn0264984 | CR44135 | lncRNA_gene | 3R | 11977640 | 11978178 | long non-coding RNA:CR44135 | lncRNA:CR44135 | YES |
| FBgn0265157 | CR44226 | lncRNA_gene | 3R | 23040776 | 23041340 | long non-coding RNA:CR44226 | lncRNA:CR44226 | NO |
| FBgn0265383 | CR44324 | lncRNA_gene | 3R | 23485481 | 23486085 | long non-coding RNA:CR44324 | lncRNA:CR44324 | NO |
| FBgn0265896 | CR44685 | lncRNA_gene | 3L | 20652906 | 20654126 | long non-coding RNA:CR44685 | lncRNA:CR44685 | YES |
| FBgn0266321 | CR44986 | lncRNA_gene | 2L | 5947647 | 5948397 | long non-coding RNA:CR44986 | lncRNA:CR44986 | NO |
| FBgn0266347* | CG12414 | protein_coding_gene | 3L | 23206669 | 23287448 | nicotinic Acetylcholine Receptor alpha4 | nAChRalpha4 | YES |
| FBgn0266578 | CR45109 | lncRNA_gene | 3R | 13132429 | 13155687 | long non-coding RNA:CR45109 | lncRNA:CR45109 | YES |
| FBgn0266657* | CR45164 | antisense_lncRNA_gene | 3L | 23149530 | 23150668 | antisense RNA:CR45164 | asRNA:CR45164 | YES |
| FBgn0266741* | CR45214 | antisense_lncRNA_gene | 3R | 25266813 | 25270007 | antisense RNA:CR45214 | asRNA:CR45214 | NO |
| FBgn0266826 | CR45288 | lncRNA_gene | 2L | 5940033 | 5940250 | long non-coding RNA:CR45288 | lncRNA:CR45288 | NO |
| FBgn0266827 | CR45289 | lncRNA_gene | 2L | 5941995 | 5942584 | long non-coding RNA:CR45289 | lncRNA:CR45289 | NO |
| FBgn0266937* | CR45388 | lncRNA_gene | 3L | 18852573 | 18861363 | long non-coding RNA:CR45388 | lncRNA:CR45388 | NO |
| FBgn0266981* | CR45432 | lncRNA_gene | 3L | 21301477 | 21302159 | long non-coding RNA:CR45432 | lncRNA:CR45432 | NO |
| FBgn0267234 | CR45674 | lncRNA_gene | 3L | 21318223 | 21318650 | long non-coding RNA:CR45674 | lncRNA:CR45674 | NO |
| FBgn0267572* | CR45912 | antisense_lncRNA_gene | 3L | 18858690 | 18866107 | antisense RNA:CR45912 | asRNA:CR45912 | YES |
| FBgn0267604 | CR45942 | lncRNA_gene | 3L | 20159990 | 20160486 | long non-coding RNA:CR45942 | lncRNA:CR45942 | NO |
| FBgn0267614* | CR45952 | antisense_lncRNA_gene | 3L | 21302421 | 21304135 | antisense RNA:CR45952 | asRNA:CR45952 | NO |
| FBgn0267683 | CR46019 | lncRNA_gene | 3R | 13105070 | 13105491 | long non-coding RNA:CR46019 | lncRNA:CR46019 | NO |
| FBgn0267737 | CR46068 | antisense_lncRNA_gene | 3L | 21333010 | 21333511 | antisense RNA:CR46068 | asRNA:CR46068 | YES |
| FBgn0267761 | CR46092 | antisense_lncRNA_gene | 3R | 23957305 | 23958388 | antisense RNA:CR46092 | asRNA:CR46092 | NO |
| FBgn0283536 | CG6213 | protein_coding_gene | 3R | 19643018 | 19644386 | Vacuolar H[+] ATPase 13kD subunit | Vha13 | NO |
| FBgn0287600* | CR46469 | antisense_lncRNA_gene | 3R | 23588885 | 23593151 | antisense RNA:CR46469 | asRNA:CR46469 | NO |
| FBti0019356 | - | transposable_element | 3R | 11065640 | 11071240 | - | 297{}1289 | NO |
| FBti0019365 | - | transposable_element | 3R | 13135881 | 13138000 | - | pogo{}1350 | YES |
| FBti0019413 | - | transposable_element | 3R | 21141729 | 21150154 | - | Tirant{}1399 | YES |
| FBti0020187 | - | transposable_element | 3L | 21306203 | 21315303 | - | roo{}1076 | NO |
| FBti0020221 | - | transposable_element | 3L | 23049900 | 23053301 | - | Cr1a{}1110 | YES |
| FBti0020222 | - | transposable_element | 3L | 23053311 | 23054210 | - | Cr1a{}1111 | YES |
| FBti0020223* | - | transposable_element | 3L | 23055986 | 23057424 | - | transib3{}1112 | YES |
| FBti0020224* | - | transposable_element | 3L | 23061575 | 23062678 | - | 1360{}1113 | YES |
| FBti0020225 | - | transposable_element | 3L | 23065350 | 23066420 | - | 1360{}1114 | YES |
| FBti0020226 | - | transposable_element | 3L | 23070953 | 23071830 | - | mariner2{}1115 | YES |
| FBti0020227* | - | transposable_element | 3L | 23072281 | 23079602 | - | 17.6{}1116 | YES |
| FBti0020229* | - | transposable_element | 3L | 23083668 | 23084655 | - | 1360{}1118 | YES |
| FBti0020230 | - | transposable_element | 3L | 23089320 | 23090522 | - | 1360{}1119 | NO |
| FBti0020231 | - | transposable_element | 3L | 23096281 | 23097342 | - | S2{}1120 | NO |
| FBti0020232 | - | transposable_element | 3L | 23097850 | 23099411 | - | S{}1121 | NO |
| FBti0020244 | - | transposable_element | 3L | 23146549 | 23147584 | - | 1360{}1133 | YES |
| FBti0020284* | - | transposable_element | 3L | 23221385 | 23224264 | - | X{}1173 | YES |
| FBti0020322 | - | transposable_element | 3R | 7849700 | 7850806 | - | 1360{}1257 | YES |
| FBti0059657* | - | transposable_element | 3L | 23027806 | 23029190 | - | Cr1a{}1596 | NO |
| FBti0059658* | - | transposable_element | 3L | 23219958 | 23220245 | - | X{}1597 | YES |
| FBti0059664* | - | transposable_element | 3L | 23080432 | 23081420 | - | S2{}1603 | YES |
| FBti0059667* | - | transposable_element | 3L | 23219852 | 23224264 | - | X{}1606 | YES |
| FBti0059783 | - | transposable_element | 3R | 21560371 | 21560421 | - | INE-1{}1722 | YES |
| FBti0060305 | - | transposable_element | 3R | 23593292 | 23593418 | - | INE-1{}2244 | NO |
| FBti0061276 | - | transposable_element | 3L | 23022458 | 23023339 | - | INE-1{}3215 | NO |
| FBti0061278 | - | transposable_element | 3L | 23022550 | 23023214 | - | INE-1{}3217 | NO |
| FBti0061281 | - | transposable_element | 3L | 23023576 | 23023803 | - | INE-1{}3220 | NO |
| FBti0061306 | - | transposable_element | 3L | 23054956 | 23055001 | - | INE-1{}3245 | YES |
| FBti0061309* | - | transposable_element | 3L | 23055008 | 23055408 | - | INE-1{}3248 | YES |
| FBti0061358 | - | transposable_element | 3L | 23026952 | 23027092 | - | flea{}3297 | NO |
| FBti0061372 | - | transposable_element | 3L | 23054474 | 23054955 | - | R1A1{}3311 | YES |
| FBti0061412* | - | transposable_element | 3L | 23081775 | 23081951 | - | INE-1{}3351 | YES |
| FBti0061414* | - | transposable_element | 3L | 23083575 | 23084756 | - | INE-1{}3353 | YES |
| FBti0061416 | - | transposable_element | 3L | 23085084 | 23085209 | - | INE-1{}3355 | YES |
| FBti0061419 | - | transposable_element | 3L | 23088679 | 23088868 | - | INE-1{}3358 | NO |
| FBti0061545 | - | transposable_element | 3R | 13105556 | 13105592 | - | INE-1{}3484 | NO |
| FBti0061816 | - | transposable_element | 3L | 23216353 | 23216596 | - | INE-1{}3755 | YES |
| FBti0061817 | - | transposable_element | 3L | 23216718 | 23216758 | - | INE-1{}3756 | YES |
| FBti0061819 | - | transposable_element | 3L | 23218788 | 23219072 | - | INE-1{}3758 | YES |
| FBti0061822 | - | transposable_element | 3L | 23225809 | 23226574 | - | INE-1{}3761 | NO |
| FBti0061859 | - | transposable_element | 3L | 23215676 | 23215900 | - | gypsy{}3798 | YES |
| FBti0061861 | - | transposable_element | 3L | 23086918 | 23087165 | - | F{}3800 | YES |
| FBti0061868 | - | transposable_element | 3L | 23219681 | 23219797 | - | 297{}3807 | YES |
| FBti0061871 | - | transposable_element | 3L | 23214167 | 23214297 | - | mdg1{}3810 | YES |
| FBti0061888 | - | transposable_element | 3L | 23214600 | 23214924 | - | mdg1{}3827 | YES |
| FBti0061891 | - | transposable_element | 3L | 23152717 | 23153182 | - | springer{}3830 | YES |
| FBti0061930 | - | transposable_element | 3L | 23157197 | 23164583 | - | antonia{}3869 | NO |
| FBti0061976 | - | transposable_element | 3L | 23087166 | 23087579 | - | Doc4{}3915 | YES |
| FBti0061988 | - | transposable_element | 3L | 23145011 | 23145542 | - | Cr1a{}3927 | YES |
| FBti0061993* | - | transposable_element | 3L | 23069429 | 23069712 | - | hopper2{}3932 | YES |
| FBti0062002* | - | transposable_element | 3L | 23069746 | 23070677 | - | HB{}3941 | YES |
| FBti0062010 | - | transposable_element | 3L | 23142884 | 23143311 | - | invader1{}3949 | YES |
| FBti0062011 | - | transposable_element | 3L | 23144415 | 23144487 | - | Max{}3950 | YES |
| FBti0062063 | - | transposable_element | 3L | 23032403 | 23032718 | - | G{}4002 | NO |
| FBti0062445 | - | transposable_element | 3L | 23092505 | 23092993 | - | diver2{}4384 | NO |
| FBti0062483* | - | transposable_element | 3L | 23065031 | 23065345 | - | gypsy4{}4422 | YES |
| FBti0062484* | - | transposable_element | 3L | 23066618 | 23068219 | - | gypsy4{}4423 | YES |
| FBti0062488* | - | transposable_element | 3L | 23062982 | 23063035 | - | transib3{}4427 | YES |
| FBti0062491* | - | transposable_element | 3L | 23151110 | 23155317 | - | gypsy6{}4430 | YES |
| FBti0062516 | - | transposable_element | 3L | 23155855 | 23156631 | - | gypsy8{}4455 | YES |
| FBti0062534* | - | transposable_element | 3L | 23057480 | 23060851 | - | ninja-Dsim-like{}4473 | YES |
| FBti0062542* | - | transposable_element | 3L | 23064787 | 23064970 | - | gypsy9{}4481 | YES |
| FBti0062543 | - | transposable_element | 3L | 23066421 | 23066617 | - | gypsy9{}4482 | YES |
| FBti0062545 | - | transposable_element | 3L | 23152512 | 23152716 | - | gypsy3{}4484 | YES |
| FBti0062550 | - | transposable_element | 3L | 23085382 | 23086899 | - | Fw3{}4489 | YES |
| FBti0063829 | - | transposable_element | 3R | 20305992 | 20306025 | - | jockey{}5768 | NO |
| FBti0063902 | - | transposable_element | 3L | 20641114 | 20641206 | - | INE-1{}5841 | YES |
| FBti0063917 | - | transposable_element | 3R | 6820676 | 6820837 | - | X{}5856 | NO |
| FBti0063919 | - | transposable_element | 3R | 6822092 | 6822253 | - | X{}5858 | NO |
| FBtr0304507 | - | - | - | - | - | - | - | NO |
| FBtr0472757 | - | - | - | - | - | - | - | NO |

**Table S8.**
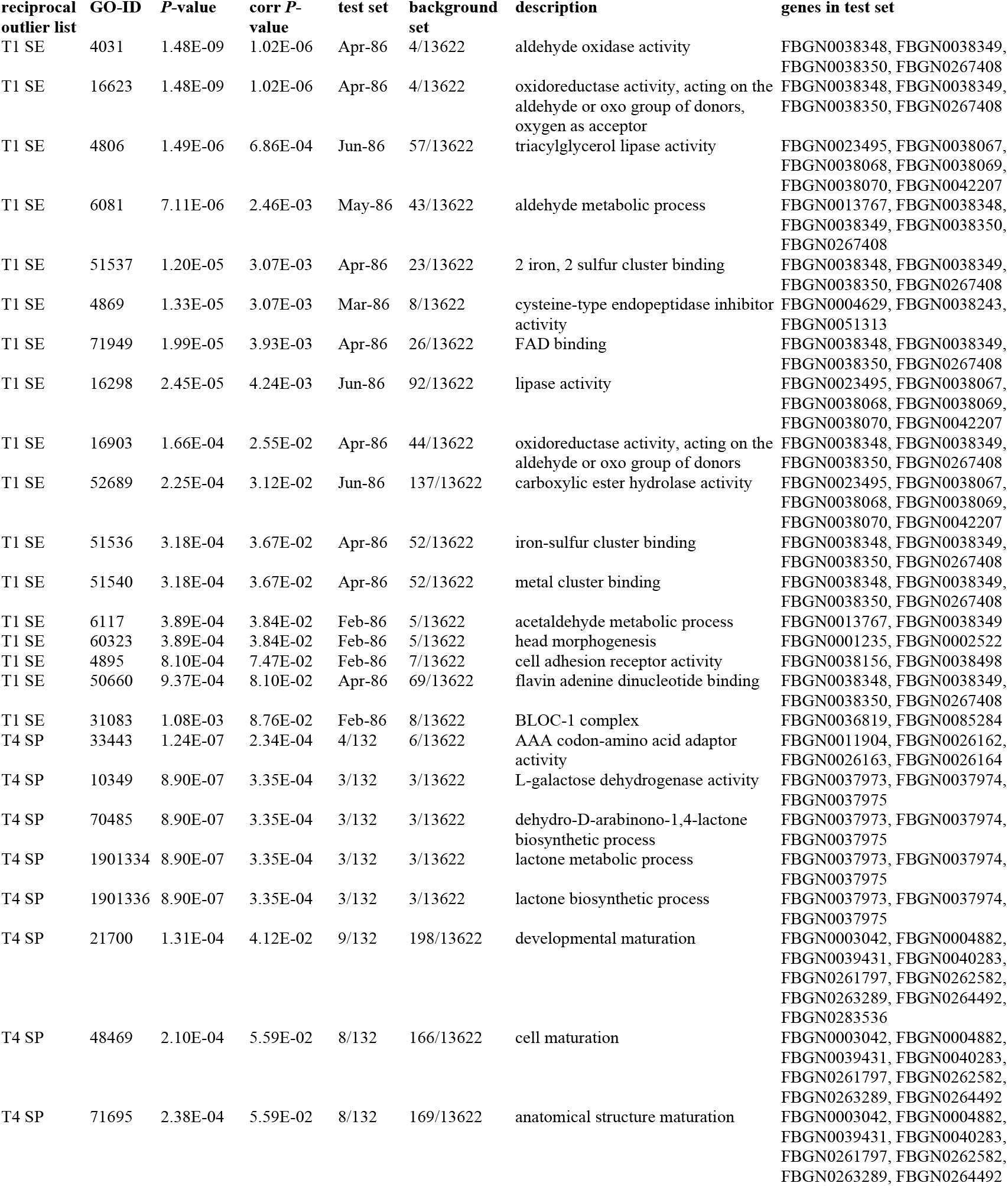
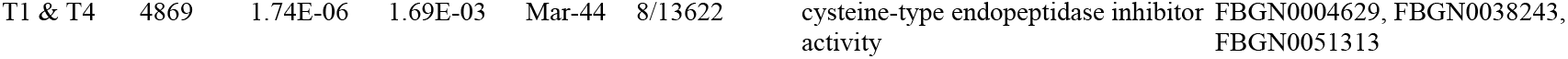
Enriched Gene Ontology categories among convergent candidate gene sets. Gene Ontology (GO) enrichment results for gene sets overlapping, or immediately adjacent to, convergent outlier SNPs identified in extinct populations in August, rescue populations in November, and an intersectional list of the two contrasts using relaxed GLM FDR thresholds. Enrichment was assessed using empirical outlier gene lists derived from filtered GLM contrasts, including genes harboring candidate SNPs and genes located within regions flanking adaptive peaks. Reported GO terms span molecular function, biological process, and cellular component categories that were significantly enriched relative to the *Drosophila melanogaster* genomic background (*n* = 13,622 genes). Enrichment was assessed using BiNGO in Cytoscape, with categories considered significant at corrected P < 0.1. Test set and background set columns give the number of genes in the category relative to the total gene list size. Abbreviations in the first column: ‘T1 SE’ = extinct populations in August, ‘T4 SP’ = rescue populations in November, ‘T1 & T4’ = features shared between both sets using relaxed filtering criteria.

| **reciprocal outlier list** | **GO-ID** | ***P*-value** | **corr *P*-value** | **test set** | **background set** | **description** | **genes in test set** |
|---|---|---|---|---|---|---|---|
| T1 SE | 4031 | 1.48E-09 | 1.02E-06 | Apr-86 | 4/13622 | aldehyde oxidase activity | FBGN0038348, FBGN0038349, FBGN0038350, FBGN0267408 |
| T1 SE | 16623 | 1.48E-09 | 1.02E-06 | Apr-86 | 4/13622 | oxidoreductase activity, acting on the aldehyde or oxo group of donors, oxygen as acceptor | FBGN0038348, FBGN0038349, FBGN0038350, FBGN0267408 |
| T1 SE | 4806 | 1.49E-06 | 6.86E-04 | Jun-86 | 57/13622 | triacylglycerol lipase activity | FBGN0023495, FBGN0038067, FBGN0038068, FBGN0038069, FBGN0038070, FBGN0042207 |
| T1 SE | 6081 | 7.11E-06 | 2.46E-03 | May-86 | 43/13622 | aldehyde metabolic process | FBGN0013767, FBGN0038348, FBGN0038349, FBGN0038350, FBGN0267408 |
| T1 SE | 51537 | 1.20E-05 | 3.07E-03 | Apr-86 | 23/13622 | 2 iron, 2 sulfur cluster binding | FBGN0038348, FBGN0038349, FBGN0038350, FBGN0267408 |
| T1 SE | 4869 | 1.33E-05 | 3.07E-03 | Mar-86 | 8/13622 | cysteine-type endopeptidase inhibitor activity | FBGN0004629, FBGN0038243, FBGN0051313 |
| T1 SE | 71949 | 1.99E-05 | 3.93E-03 | Apr-86 | 26/13622 | FAD binding | FBGN0038348, FBGN0038349, FBGN0038350, FBGN0267408 |
| T1 SE | 16298 | 2.45E-05 | 4.24E-03 | Jun-86 | 92/13622 | lipase activity | FBGN0023495, FBGN0038067, FBGN0038068, FBGN0038069, FBGN0038070, FBGN0042207 |
| T1 SE | 16903 | 1.66E-04 | 2.55E-02 | Apr-86 | 44/13622 | oxidoreductase activity, acting on the aldehyde or oxo group of donors | FBGN0038348, FBGN0038349, FBGN0038350, FBGN0267408 |
| T1 SE | 52689 | 2.25E-04 | 3.12E-02 | Jun-86 | 137/13622 | carboxylic ester hydrolase activity | FBGN0023495, FBGN0038067, FBGN0038068, FBGN0038069, FBGN0038070, FBGN0042207 |
| T1 SE | 51536 | 3.18E-04 | 3.67E-02 | Apr-86 | 52/13622 | iron-sulfur cluster binding | FBGN0038348, FBGN0038349, FBGN0038350, FBGN0267408 |
| T1 SE | 51540 | 3.18E-04 | 3.67E-02 | Apr-86 | 52/13622 | metal cluster binding | FBGN0038348, FBGN0038349, FBGN0038350, FBGN0267408 |
| T1 SE | 6117 | 3.89E-04 | 3.84E-02 | Feb-86 | 5/13622 | acetaldehyde metabolic process | FBGN0013767, FBGN0038349 |
| T1 SE | 60323 | 3.89E-04 | 3.84E-02 | Feb-86 | 5/13622 | head morphogenesis | FBGN0001235, FBGN0002522 |
| T1 SE | 4895 | 8.10E-04 | 7.47E-02 | Feb-86 | 7/13622 | cell adhesion receptor activity | FBGN0038156, FBGN0038498 |
| T1 SE | 50660 | 9.37E-04 | 8.10E-02 | Apr-86 | 69/13622 | flavin adenine dinucleotide binding | FBGN0038348, FBGN0038349, FBGN0038350, FBGN0267408 |
| T1 SE | 31083 | 1.08E-03 | 8.76E-02 | Feb-86 | 8/13622 | BLOC-1 complex | FBGN0036819, FBGN0085284 |
| T4 SP | 33443 | 1.24E-07 | 2.34E-04 | 4/132 | 6/13622 | AAA codon-amino acid adaptor activity | FBGN0011904, FBGN0026162, FBGN0026163, FBGN0026164 |
| T4 SP | 10349 | 8.90E-07 | 3.35E-04 | 3/132 | 3/13622 | L-galactose dehydrogenase activity | FBGN0037973, FBGN0037974, FBGN0037975 |
| T4 SP | 70485 | 8.90E-07 | 3.35E-04 | 3/132 | 3/13622 | dehydro-D-arabinono-1,4-lactone biosynthetic process | FBGN0037973, FBGN0037974, FBGN0037975 |
| T4 SP | 1901334 | 8.90E-07 | 3.35E-04 | 3/132 | 3/13622 | lactone metabolic process | FBGN0037973, FBGN0037974, FBGN0037975 |
| T4 SP | 1901336 | 8.90E-07 | 3.35E-04 | 3/132 | 3/13622 | lactone biosynthetic process | FBGN0037973, FBGN0037974, FBGN0037975 |
| T4 SP | 21700 | 1.31E-04 | 4.12E-02 | 9/132 | 198/13622 | developmental maturation | FBGN0003042, FBGN0004882, FBGN0039431, FBGN0040283, FBGN0261797, FBGN0262582, FBGN0263289, FBGN0264492, FBGN0283536 |
| T4 SP | 48469 | 2.10E-04 | 5.59E-02 | 8/132 | 166/13622 | cell maturation | FBGN0003042, FBGN0004882, FBGN0039431, FBGN0040283, FBGN0261797, FBGN0262582, FBGN0263289, FBGN0264492 |
| T4 SP | 71695 | 2.38E-04 | 5.59E-02 | 8/132 | 169/13622 | anatomical structure maturation | FBGN0003042, FBGN0004882, FBGN0039431, FBGN0040283, FBGN0261797, FBGN0262582, FBGN0263289, FBGN0264492 |
| T1 & T4 | 4869 | 1.74E-06 | 1.69E-03 | Mar-44 | 8/13622 | cysteine-type endopeptidase inhibitor activity | FBGN0004629, FBGN0038243, FBGN0051313 |

**Table S9.** Overlap between parallel temporal allele frequency change in pre-adapted populations and August pre-adapted outliers. At each SNP, the direction of pre-adapted allelic change was defined from the August ‘unexposed vs. pre-adapted’ contrast as the direction in which pre-adapted populations differed from unexposed populations, and the direction of unexposed allelic change as the opposite direction. For each temporal interval, allele frequency change in each pre-adapted population was assigned a sign so that positive values indicate change in the direction of pre-adapted allelic change. SNPs were classified as changing in parallel in the direction of pre-adapted allelic change (‘pre-adapted’) if all, or all but one, populations changed by more than 0.03 in that direction, and in the direction of unexposed allelic change (‘unexposed’) if all, or all but one, populations changed by more than 0.03 in the opposite direction. Hypergeometric tests assess whether these SNPs (‘temporal outliers’) overlap with SNPs differentiated between unexposed and pre-adapted populations in August (‘PAvE.T1 outliers’; GLM FDR < 0.05) more than expected by chance; ‘non-temporal outliers’ gives the number of remaining SNPs. All ten pre-adapted populations were included. Time points are T1 = August 24, T2 = September 30, T3 = October 27, and T4 = November 24, 2023.

| **contrast** | **direction** | **overlap** | **temporal outliers** | **non-temporal outliers** | **PAvE.T1 outliers** | **hypergeometric *P*-value** |
|---|---|---|---|---|---|---|
| T1 vs. T2 | pre-adapted | 1141 | 3699 | 1092140 | 455515 | 1 |
| T1 vs. T3 | pre-adapted | 1183 | 7094 | 1088745 | 455515 | 1 |
| T1 vs. T4 | pre-adapted | 4372 | 19341 | 1076498 | 455515 | 1 |
| T2 vs. T3 | pre-adapted | 85 | 222 | 1095617 | 455515 | 0.85546927 |
| T2 vs. T4 | pre-adapted | 909 | 3586 | 1092253 | 455515 | 1 |
| T3 vs. T4 | pre-adapted | 531 | 1852 | 1093987 | 455515 | 1 |
| T1 vs. T2 | unexposed | 2659 | 6134 | 1089705 | 455515 | 0.00239912 |
| T1 vs. T3 | unexposed | 9514 | 19497 | 1076342 | 455515 | 5.91E-94 |
| T1 vs. T4 | unexposed | 23217 | 43604 | 1052235 | 455515 | 0 |
| T2 vs. T3 | unexposed | 125 | 262 | 1095577 | 455515 | 0.02577876 |
| T2 vs. T4 | unexposed | 5184 | 8761 | 1087078 | 455515 | 1.49E-242 |
| T3 vs. T4 | unexposed | 677 | 1838 | 1094001 | 455515 | 0.99998479 |

**Table S10.** Genomic features associated with convergent candidate loci in extinct populations in August. Summary of features within 5 kb of convergent candidate SNPs identified in exposed populations that subsequently went extinct, at the first post-treatment time point (August 24). Outlier loci were defined as SNPs significant in the ‘unexposed vs. extinct’ contrast in August (FDR < 0.05) and in the ‘unexposed vs. pre-adapted’ contrasts in both August and November (FDR < 0.01), with unexposed allele frequencies of 0.1–0.9 and an allele frequency difference > 0.1 between unexposed and pre-adapted populations in August. FlyCADD scores were used to prioritize variants with predicted functional effects; ‘YES’ indicates features containing or flanking at least one candidate SNP with a FlyCADD score > 0.60. FlyBase IDs marked with an asterisk contain candidate SNPs directly, whereas unmarked entries lie within 5 kb upstream or downstream flanking regions.

| **ID** | **SYMBOL** | **TYPE** | **CHROM** | **START** | **STOP** | **NAME** | **GENE SYMBOL** | **FLYCADD > 0.60** |
|---|---|---|---|---|---|---|---|---|
| FBgn0000246 | CG17604 | protein_coding_gene | 3R | 15789473 | 15792579 | crossover suppressor on 3 of Gowen | c(3)G | NO |
| FBgn0001235* | CG17117 | protein_coding_gene | 3R | 10507561 | 10639568 | homothorax | hth | NO |
| FBgn0001323* | CG4761 | protein_coding_gene | 3L | 20597141 | 20620659 | knirps-like | knrl | YES |
| FBgn0002522* | CG1264 | protein_coding_gene | 3R | 6661427 | 6678590 | labial | lab | YES |
| FBgn0003042 | CG32443 | protein_coding_gene | 3L | 21306138 | 21318020 | Polycomb | Pc | YES |
| FBgn0004117 | CG4843 | protein_coding_gene | 3R | 15310193 | 15315197 | Tropomyosin 2 | Tm2 | NO |
| FBgn0004577* | CG3477 | protein_coding_gene | 3R | 17014576 | 17027987 | Peroxidase | Pxd | NO |
| FBgn0004629 | CG8050 | protein_coding_gene | 3R | 14569103 | 14569644 | Cystatin-like | Cys | NO |
| FBgn0011474* | CG3307 | protein_coding_gene | 3R | 14316038 | 14320494 | SET domain containing 8 | Set8 | YES |
| FBgn0011582* | CG9652 | protein_coding_gene | 3R | 14165379 | 14214810 | Dopamine 1-like receptor 1 | Dop1R1 | NO |
| FBgn0012034 | CG9390 | protein_coding_gene | 3L | 21270722 | 21280243 | Acetyl Coenzyme A synthase | AcCoAS | YES |
| FBgn0013767* | CG3302 | protein_coding_gene | 3R | 14314854 | 14315716 | Corazonin | Crz | YES |
| FBgn0015946 | CG4345 | protein_coding_gene | 3L | 18302719 | 18304417 | grim | grim | NO |
| FBgn0019925* | CG6202 | protein_coding_gene | 3R | 15343653 | 15346176 | Surfeit 4 | Surf4 | YES |
| FBgn0020299 | CG31317 | protein_coding_gene | 3R | 14577217 | 14608079 | stumps | stumps | NO |
| FBgn0021750* | CG4938 | protein_coding_gene | 3R | 15340667 | 15343218 | Seryl-tRNA synthetase, mitochondrial | SerRS-m | YES |
| FBgn0023495 | CG8823 | protein_coding_gene | 3R | 13370240 | 13371904 | Lipase 3 | Lip3 | YES |
| FBgn0025807 | CG3945 | protein_coding_gene | 3L | 18870045 | 18872553 | Rad9 | Rad9 | YES |
| FBgn0026441 | CG4913 | protein_coding_gene | 3R | 15320756 | 15325279 | ENL/AF9-related | ear | NO |
| FBgn0026616* | CG4606 | protein_coding_gene | 3R | 15804248 | 15824118 | alpha-Mannosidase class IIb | alpha-Man-IIb | YES |
| FBgn0027610 | CG8790 | protein_coding_gene | 3R | 13364938 | 13369278 | Dicarboxylate carrier 1 | Dic1 | YES |
| FBgn0027945 | CG7758 | protein_coding_gene | 3L | 21280491 | 21281506 | pumpless | ppl | YES |
| FBgn0036785 | CG13700 | protein_coding_gene | 3L | 18298742 | 18301260 | - | CG13700 | NO |
| FBgn0036786 | CG13701 | protein_coding_gene | 3L | 18438561 | 18439942 | sickle | skl | NO |
| FBgn0036817* | CG6865 | protein_coding_gene | 3L | 18863647 | 18865514 | - | CG6865 | YES |
| FBgn0036818* | CG14074 | protein_coding_gene | 3L | 18866225 | 18867431 | - | CG14074 | YES |
| FBgn0036819 | CG6856 | protein_coding_gene | 3L | 18867771 | 18869219 | Dysbindin | Dysb | YES |
| FBgn0036820 | CG6852 | protein_coding_gene | 3L | 18869230 | 18870057 | Glutaredoxin 1 | Grx1 | YES |
| FBgn0037015* | CG13253 | protein_coding_gene | 3L | 20712838 | 20719200 | crimpy | cmpy | YES |
| FBgn0037016 | CG13252 | protein_coding_gene | 3L | 20730591 | 20744508 | - | CG13252 | NO |
| FBgn0037065* | CG12974 | protein_coding_gene | 3L | 21281752 | 21285962 | - | CG12974 | YES |
| FBgn0037073 | CG7338 | protein_coding_gene | 3L | 21335215 | 21338232 | Tsr1 ribosome assembly factor | Tsr1 | NO |
| FBgn0037074 | CG7324 | protein_coding_gene | 3L | 21337681 | 21343716 | TBC1 domain family member 8/9 | Tbc1d8-9 | NO |
| FBgn0038067 | CG11598 | protein_coding_gene | 3R | 12634917 | 12636283 | - | CG11598 | NO |
| FBgn0038068* | CG11600 | protein_coding_gene | 3R | 12636540 | 12637976 | - | CG11600 | NO |
| FBgn0038069 | CG11608 | protein_coding_gene | 3R | 12638481 | 12640033 | - | CG11608 | NO |
| FBgn0038070 | CG6753 | protein_coding_gene | 3R | 12640163 | 12642482 | - | CG6753 | NO |
| FBgn0038142* | CG14379 | protein_coding_gene | 3R | 13369648 | 13370346 | Chemosensory protein A 87a | CheA87a | YES |
| FBgn0038143 | CG9813 | protein_coding_gene | 3R | 13373104 | 13379775 | - | CG9813 | YES |
| FBgn0038147 | CG14375 | protein_coding_gene | 3R | 13389059 | 13395197 | CCHamide-2 | CCHa2 | NO |
| FBgn0038148* | CG14377 | protein_coding_gene | 3R | 13397835 | 13398583 | - | CG14377 | NO |
| FBgn0038149 | CG9796 | protein_coding_gene | 3R | 13398526 | 13401862 | Gamma-interferon-inducible lysosomal thiol reductase 1 | GILT1 | NO |
| FBgn0038156* | CG14372 | protein_coding_gene | 3R | 13450590 | 13464184 | sidestep IV | side-IV | YES |
| FBgn0038223 | CG8538 | protein_coding_gene | 3R | 14320191 | 14326689 | Aftiphilin | Afti | YES |
| FBgn0038237* | CG8279 | protein_coding_gene | 3R | 14513901 | 14558304 | Phosphodiesterase 6 | Pde6 | NO |
| FBgn0038238 | CG14854 | protein_coding_gene | 3R | 14559166 | 14560054 | - | CG14854 | NO |
| FBgn0038239 | CG14850 | protein_coding_gene | 3R | 14560935 | 14561521 | Jig | Jig | NO |
| FBgn0038240 | CG14851 | protein_coding_gene | 3R | 14562340 | 14562845 | - | CG14851 | NO |
| FBgn0038241 | CG8087 | protein_coding_gene | 3R | 14564323 | 14564805 | - | CG8087 | NO |
| FBgn0038242 | CG14852 | protein_coding_gene | 3R | 14565927 | 14566513 | - | CG14852 | NO |
| FBgn0038243 | CG8066 | protein_coding_gene | 3R | 14567611 | 14568847 | - | CG8066 | NO |
| FBgn0038246* | CG14853 | protein_coding_gene | 3R | 14609642 | 14618987 | - | CG14853 | NO |
| FBgn0038315 | CG14866 | protein_coding_gene | 3R | 15316471 | 15318285 | - | CG14866 | NO |
| FBgn0038316* | CG6276 | protein_coding_gene | 3R | 15318203 | 15320069 | - | CG6276 | NO |
| FBgn0038320 | CG4931 | protein_coding_gene | 3R | 15333266 | 15338365 | Cytoplasmic FMR1 interacting protein | Cyfip | YES |
| FBgn0038321* | CG6218 | protein_coding_gene | 3R | 15337908 | 15343256 | N-acetylglucosamine kinase | Nagk | YES |
| FBgn0038348* | CG18519 | protein_coding_gene | 3R | 15537713 | 15542414 | Aldehyde oxidase 2 | AOX2 | YES |
| FBgn0038349* | CG6045 | protein_coding_gene | 3R | 15542748 | 15548188 | Aldehyde oxidase 3 | AOX3 | YES |
| FBgn0038350* | CG18516 | protein_coding_gene | 3R | 15548628 | 15552833 | Aldehyde oxidase 4 | AOX4 | YES |
| FBgn0038363 | CG18505 | protein_coding_gene | 3R | 15792688 | 15793487 | Acylphosphatase 2 | Acyp2 | NO |
| FBgn0038365 | CG9593 | protein_coding_gene | 3R | 15802275 | 15803653 | - | CG9593 | YES |
| FBgn0038404 | CG8925 | protein_coding_gene | 3R | 16171991 | 16180475 | - | CG8925 | NO |
| FBgn0038467 | CG3590 | protein_coding_gene | 3R | 17018780 | 17021120 | Adenylosuccinate lyase | Adsl | NO |
| FBgn0038468 | CG5225 | protein_coding_gene | 3R | 17028693 | 17030701 | - | CG5225 | NO |
| FBgn0038469* | CG4009 | protein_coding_gene | 3R | 17033588 | 17036045 | - | CG4009 | NO |
| FBgn0038470 | CG18213 | protein_coding_gene | 3R | 17036394 | 17040203 | - | CG18213 | NO |
| FBgn0038498* | CG14334 | protein_coding_gene | 3R | 17320004 | 17366119 | beaten path IIa | beat-IIa | NO |
| FBgn0040553 | CG14374 | protein_coding_gene | 3R | 13396694 | 13397140 | - | CG14374 | NO |
| FBgn0040554* | CR17025 | antisense_lncRNA_gene | 3R | 13455511 | 13466106 | antisense RNA:CR17025 | asRNA:CR17025 | YES |
| FBgn0042207 | CG18530 | protein_coding_gene | 3R | 12633359 | 12634670 | - | CG18530 | NO |
| FBgn0051268* | CG31268 | protein_coding_gene | 3R | 17031802 | 17033515 | - | CG31268 | NO |
| FBgn0051296 | CG31296 | protein_coding_gene | 3R | 15530072 | 15531473 | - | CG31296 | NO |
| FBgn0051301 | CG31301 | protein_coding_gene | 3R | 15346205 | 15348012 | - | CG31301 | YES |
| FBgn0051313 | CG31313 | protein_coding_gene | 3R | 14566432 | 14567429 | - | CG31313 | NO |
| FBgn0051321 | CG31321 | protein_coding_gene | 3R | 14310362 | 14314412 | - | CG31321 | YES |
| FBgn0051331 | CR31331 | tRNA_gene | 3R | 13798198 | 13798270 | transfer RNA:Phenylalanine-GAA 1-7 | tRNA:Phe-GAA-1-7 | NO |
| FBgn0051481* | CG31481 | protein_coding_gene | 3R | 6707800 | 6742063 | proboscipedia | pb | YES |
| FBgn0051588 | CR31588 | tRNA_gene | 3R | 13797953 | 13798025 | transfer RNA:Phenylalanine-GAA 1-6 | tRNA:Phe-GAA-1-6 | NO |
| FBgn0052204* | CG32204 | protein_coding_gene | 3L | 18922419 | 18946038 | - | CG32204 | NO |
| FBgn0053977 | CG33977 | protein_coding_gene | 3R | 13402559 | 13403051 | Dolichyl-phosphate mannosyltransferase subunit 3 | Dpm3 | NO |
| FBgn0063649* | CG6006 | protein_coding_gene | 3R | 16154982 | 16171766 | - | CG6006 | NO |
| FBgn0085284 | CG34255 | protein_coding_gene | 3L | 18870224 | 18870897 | Biogenesis of lysosome-related organelles complex 1, subunit 3 | Blos3 | YES |
| FBgn0085289 | CG34260 | protein_coding_gene | 3L | 20714100 | 20715158 | - | CG34260 | YES |
| FBgn0086371 | CG9829 | protein_coding_gene | 3R | 13361816 | 13366910 | Elongator complex protein 6 | Elp6 | YES |
| FBgn0086913* | CG34410 | protein_coding_gene | 3L | 21318774 | 21335027 | Rab26 | Rab26 | YES |
| FBgn0259785 | CG7752 | protein_coding_gene | 3L | 21286099 | 21290310 | putzig | pzg | YES |
| FBgn0259831 | CG34309 | protein_coding_gene | 3R | 13372630 | 13373326 | - | CG34309 | YES |
| FBgn0260444 | CG18616 | protein_coding_gene | 3R | 12630647 | 12633188 | CCR4-NOT transcription complex subunit 10 | Not10 | NO |
| FBgn0261679 | CG42726 | protein_coding_gene | 3R | 15348048 | 15350391 | - | CG42726 | YES |
| FBgn0261680 | CG42727 | protein_coding_gene | 3R | 15348048 | 15350391 | - | CG42727 | YES |
| FBgn0262461 | CR43036 | miRNA_gene | 3L | 18860971 | 18861054 | mir-315 stem loop | mir-315 | NO |
| FBgn0262527* | CG4699 | protein_coding_gene | 3R | 15793782 | 15802102 | non-specific lethal 1 | nsl1 | YES |
| FBgn0262969 | CR43280 | lncRNA_gene | 3L | 18440730 | 18441871 | long non-coding RNA:CR43280 | lncRNA:CR43280 | NO |
| FBgn0263449 | CR43472 | antisense_lncRNA_gene | 3R | 16168348 | 16169154 | antisense RNA:CR43472 | asRNA:CR43472 | NO |
| FBgn0263595* | CR43617 | lncRNA_gene | 3R | 16892410 | 16897861 | long non-coding RNA:CR43617 | lncRNA:CR43617 | NO |
| FBgn0264573 | CR43945 | lncRNA_gene | 3L | 20729274 | 20729656 | long non-coding RNA:CR43945 | lncRNA:CR43945 | NO |
| FBgn0264832 | CG44040 | protein_coding_gene | 3R | 15320378 | 15320691 | - | CG44040 | NO |
| FBgn0264857* | CR43261 | lncRNA_gene | 3R | 16831116 | 16923231 | long non-coding RNA:iab8 | lncRNA:iab8 | NO |
| FBgn0264904 | CR44095 | antisense_lncRNA_gene | 3R | 14568962 | 14569871 | antisense RNA:CR44095 | asRNA:CR44095 | NO |
| FBgn0265896* | CR44685 | lncRNA_gene | 3L | 20652906 | 20654126 | long non-coding RNA:CR44685 | lncRNA:CR44685 | YES |
| FBgn0266347* | CG12414 | protein_coding_gene | 3L | 23206669 | 23287448 | nicotinic Acetylcholine Receptor alpha4 | nAChRalpha4 | NO |
| FBgn0266415 | CR45055 | lncRNA_gene | 3R | 11310676 | 11311182 | long non-coding RNA:CR45055 | lncRNA:CR45055 | NO |
| FBgn0266738 | CR45211 | lncRNA_gene | 3R | 15811648 | 15811908 | long non-coding RNA:CR45211 | lncRNA:CR45211 | YES |
| FBgn0266756* | CG44012 | protein_coding_gene | 3R | 14804092 | 14875488 | bitesize | btsz | NO |
| FBgn0266937 | CR45388 | lncRNA_gene | 3L | 18852573 | 18861363 | long non-coding RNA:CR45388 | lncRNA:CR45388 | NO |
| FBgn0267139* | CR45579 | lncRNA_gene | 3R | 11101180 | 11101684 | long non-coding RNA:CR45579 | lncRNA:CR45579 | NO |
| FBgn0267234 | CR45674 | lncRNA_gene | 3L | 21318223 | 21318650 | long non-coding RNA:CR45674 | lncRNA:CR45674 | YES |
| FBgn0267408* | CG18522 | protein_coding_gene | 3R | 15532137 | 15537292 | Aldehyde oxidase 1 | AOX1 | YES |
| FBgn0267551 | CR45891 | antisense_lncRNA_gene | 3R | 15802854 | 15803495 | antisense RNA:CR45891 | asRNA:CR45891 | YES |
| FBgn0267572* | CR45912 | antisense_lncRNA_gene | 3L | 18858690 | 18866107 | antisense RNA:CR45912 | asRNA:CR45912 | YES |
| FBgn0267574 | CR45914 | lncRNA_gene | 3R | 13466495 | 13466722 | long non-coding RNA:CR45914 | lncRNA:CR45914 | NO |
| FBgn0267636 | CR45974 | antisense_lncRNA_gene | 3L | 18303431 | 18304179 | antisense RNA:CR45974 | asRNA:CR45974 | NO |
| FBgn0267637 | CR45975 | lncRNA_gene | 3L | 18302801 | 18303341 | long non-coding RNA:CR45975 | lncRNA:CR45975 | NO |
| FBgn0267682 | CR46018 | antisense_lncRNA_gene | 3R | 12634192 | 12634744 | antisense RNA:CR46018 | asRNA:CR46018 | NO |
| FBgn0267737 | CR46068 | antisense_lncRNA_gene | 3L | 21333010 | 21333511 | antisense RNA:CR46068 | asRNA:CR46068 | NO |
| FBgn0286034 | CR46354 | antisense_lncRNA_gene | 3R | 13459671 | 13466106 | antisense RNA:CR46354 | asRNA:CR46354 | YES |
| FBgn0288835* | CG46527 | protein_coding_gene | 3R | 16831115 | 16896293 | MSA micropeptide | MSAmiP | NO |
| FBti0019373 | - | transposable_element | 3R | 14180867 | 14185556 | - | F{}1358 | NO |
| FBti0019374 | - | transposable_element | 3R | 14194483 | 14203574 | - | roo{}1359 | NO |
| FBti0019389 | - | transposable_element | 3R | 17346744 | 17348235 | - | F{}1374 | NO |
| FBti0020148 | - | transposable_element | 3L | 18506186 | 18509586 | - | F{}1037 | YES |
| FBti0020149 | - | transposable_element | 3L | 18514973 | 18520090 | - | BS{}1038 | YES |
| FBti0020284* | - | transposable_element | 3L | 23221385 | 23224264 | - | X{}1173 | NO |
| FBti0059658 | - | transposable_element | 3L | 23219958 | 23220245 | - | X{}1597 | NO |
| FBti0059667* | - | transposable_element | 3L | 23219852 | 23224264 | - | X{}1606 | NO |
| FBti0061816 | - | transposable_element | 3L | 23216353 | 23216596 | - | INE-1{}3755 | NO |
| FBti0061817 | - | transposable_element | 3L | 23216718 | 23216758 | - | INE-1{}3756 | NO |
| FBti0061819 | - | transposable_element | 3L | 23218788 | 23219072 | - | INE-1{}3758 | NO |
| FBti0061822 | - | transposable_element | 3L | 23225809 | 23226574 | - | INE-1{}3761 | NO |
| FBti0061868 | - | transposable_element | 3L | 23219681 | 23219797 | - | 297{}3807 | NO |
| FBti0062073 | - | transposable_element | 3R | 14565185 | 14565228 | - | INE-1{}4012 | NO |
| FBti0062141 | - | transposable_element | 3R | 14927105 | 14927133 | - | INE-1{}4080 | NO |
| FBti0062746 | - | transposable_element | 3L | 18933429 | 18933497 | - | INE-1{}4685 | NO |
| FBti0063903 | - | transposable_element | 3L | 20659518 | 20659809 | - | INE-1{}5842 | NO |
| FBti0063920 | - | transposable_element | 3R | 6672583 | 6672639 | - | INE-1{}5859 | YES |
| FBtr0304507 | - | - | - | - | - | - | - | NO |
| FBtr0472757 | - | - | - | - | - | - | - | NO |

**Table S11.** Genomic features associated with convergent candidate loci shared between August and November. Convergent candidate loci were identified in both extinct populations in August and rescue populations in November. Outliers include genes containing candidate SNPs as well as genes flanked by significant upstream or downstream intergenic SNPs within 5 kb. Loci were defined using a relaxed threshold (FDR < 0.1) in the ‘unexposed vs. extinct’ (August), ‘unexposed vs. rescue’ (November), and ‘unexposed vs. pre-adapted’ (August and November) contrasts, with unexposed allele frequencies of 0.1–0.9 and an allele frequency difference > 0.1 between unexposed and pre-adapted populations in August. For each locus, the table reports FlyBase identifier, CG annotation symbol, feature type, genomic position, and gene name and symbol. “YES” in the FLYCADD > 0.60 column indicates features containing or flanking at least one candidate SNP with a FlyCADD score > 0.60. FlyBase IDs marked with an asterisk contain candidate SNPs directly, whereas unmarked entries lie within 5 kb upstream or downstream flanking regions.

| **ID** | **SYMBOL** | **TYPE** | **CHROM** | **START** | **STOP** | **NAME** | **GENE SYMBOL** | **FLYCADD > 0.60** |
|---|---|---|---|---|---|---|---|---|
| FBgn0001235* | CG17117 | protein_coding_gene | 3R | 10507561 | 10639568 | homothorax | hth | YES |
| FBgn0001323* | CG4761 | protein_coding_gene | 3L | 20597141 | 20620659 | knirps-like | knrl | YES |
| FBgn0003042 | CG32443 | protein_coding_gene | 3L | 21306138 | 21318020 | Polycomb | Pc | YES |
| FBgn0004049* | CG9764 | protein_coding_gene | 3R | 13428867 | 13434984 | yurt | yrt | NO |
| FBgn0004629 | CG8050 | protein_coding_gene | 3R | 14569103 | 14569644 | Cystatin-like | Cys | NO |
| FBgn0010355* | CG17603 | protein_coding_gene | 3R | 6646889 | 6656048 | TBP-associated factor 1 | Taf1 | NO |
| FBgn0025807 | CG3945 | protein_coding_gene | 3L | 18870045 | 18872553 | Rad9 | Rad9 | YES |
| FBgn0026314 | CG6649 | protein_coding_gene | 3R | 11168503 | 11170246 | UDP-glycosyltransferase family 35 member B1 | Ugt35B1 | NO |
| FBgn0026315 | CG6644 | protein_coding_gene | 3R | 11170817 | 11172664 | UDP-glycosyltransferase family 35 member A1 | Ugt35A1 | NO |
| FBgn0026565 | CG1315 | protein_coding_gene | 3R | 6642262 | 6644434 | Argininosuccinate synthase | Ass | NO |
| FBgn0027584 | CG4757 | protein_coding_gene | 3R | 11161207 | 11163498 | - | CG4757 | NO |
| FBgn0036786 | CG13701 | protein_coding_gene | 3L | 18438561 | 18439942 | sickle | skl | NO |
| FBgn0036817* | CG6865 | protein_coding_gene | 3L | 18863647 | 18865514 | - | CG6865 | YES |
| FBgn0036818* | CG14074 | protein_coding_gene | 3L | 18866225 | 18867431 | - | CG14074 | YES |
| FBgn0036819 | CG6856 | protein_coding_gene | 3L | 18867771 | 18869219 | Dysbindin | Dysb | YES |
| FBgn0036820 | CG6852 | protein_coding_gene | 3L | 18869230 | 18870057 | Glutaredoxin 1 | Grx1 | YES |
| FBgn0037014* | CG13251 | protein_coding_gene | 3L | 20637199 | 20640910 | - | CG13251 | NO |
| FBgn0037073 | CG7338 | protein_coding_gene | 3L | 21335215 | 21338232 | Tsr1 ribosome assembly factor | Tsr1 | YES |
| FBgn0037074* | CG7324 | protein_coding_gene | 3L | 21337681 | 21343716 | TBC1 domain family member 8/9 | Tbc1d8-9 | YES |
| FBgn0037076* | CG12972 | protein_coding_gene | 3L | 21344768 | 21349288 | earthbound 2 | ebd2 | YES |
| FBgn0037081* | CG6049 | protein_coding_gene | 3L | 21420388 | 21433134 | barricade | barc | YES |
| FBgn0037994 | CG4810 | protein_coding_gene | 3R | 12192703 | 12195011 | eukaryotic translation initiation factor 3 subunit d2 | eIF3d2 | NO |
| FBgn0037995 | CG3809 | protein_coding_gene | 3R | 12197941 | 12199473 | Adenosine kinase 1 | Adk1 | NO |
| FBgn0038147 | CG14375 | protein_coding_gene | 3R | 13389059 | 13395197 | CCHamide-2 | CCHa2 | NO |
| FBgn0038148* | CG14377 | protein_coding_gene | 3R | 13397835 | 13398583 | - | CG14377 | NO |
| FBgn0038149 | CG9796 | protein_coding_gene | 3R | 13398526 | 13401862 | Gamma-interferon-inducible lysosomal thiol reductase 1 | GILT1 | NO |
| FBgn0038239 | CG14850 | protein_coding_gene | 3R | 14560935 | 14561521 | Jig | Jig | NO |
| FBgn0038240 | CG14851 | protein_coding_gene | 3R | 14562340 | 14562845 | - | CG14851 | NO |
| FBgn0038241 | CG8087 | protein_coding_gene | 3R | 14564323 | 14564805 | - | CG8087 | NO |
| FBgn0038242 | CG14852 | protein_coding_gene | 3R | 14565927 | 14566513 | - | CG14852 | NO |
| FBgn0038243 | CG8066 | protein_coding_gene | 3R | 14567611 | 14568847 | - | CG8066 | NO |
| FBgn0038244 | CG7987 | protein_coding_gene | 3R | 14570472 | 14575887 | - | CG7987 | NO |
| FBgn0040253* | CG17200 | protein_coding_gene | 3R | 11164423 | 11166074 | UDP-glycosyltransferase family 35 member E1 | Ugt35E1 | NO |
| FBgn0040255* | CG6653 | protein_coding_gene | 3R | 11166177 | 11167981 | UDP-glycosyltransferase family 35 member E2 | Ugt35E2 | NO |
| FBgn0040553 | CG14374 | protein_coding_gene | 3R | 13396694 | 13397140 | - | CG14374 | NO |
| FBgn0051286 | CG31286 | protein_coding_gene | 3R | 6641232 | 6643346 | - | CG31286 | NO |
| FBgn0051313 | CG31313 | protein_coding_gene | 3R | 14566432 | 14567429 | - | CG31313 | NO |
| FBgn0052436* | CG32436 | protein_coding_gene | 3L | 21344033 | 21355049 | - | CG32436 | YES |
| FBgn0053288 | CG33288 | protein_coding_gene | 3L | 21427790 | 21433066 | - | CG33288 | YES |
| FBgn0053977 | CG33977 | protein_coding_gene | 3R | 13402559 | 13403051 | Dolichyl-phosphate mannosyltransferase subunit 3 | Dpm3 | NO |
| FBgn0083950* | CG34114 | protein_coding_gene | 3R | 10972381 | 11050175 | sidestep VI | side-VI | NO |
| FBgn0085284 | CG34255 | protein_coding_gene | 3L | 18870224 | 18870897 | Biogenesis of lysosome-related organelles complex 1, subunit 3 | Blos3 | YES |
| FBgn0086913* | CG34410 | protein_coding_gene | 3L | 21318774 | 21335027 | Rab26 | Rab26 | YES |
| FBgn0261258* | CG6014 | protein_coding_gene | 3L | 21382501 | 21424416 | regeneration | rgn | YES |
| FBgn0261814 | CR42756 | lncRNA_gene | 3R | 13438587 | 13439209 | long non-coding RNA:CR42756 | lncRNA:CR42756 | NO |
| FBgn0262461 | CR43036 | miRNA_gene | 3L | 18860971 | 18861054 | mir-315 stem loop | mir-315 | NO |
| FBgn0262969 | CR43280 | lncRNA_gene | 3L | 18440730 | 18441871 | long non-coding RNA:CR43280 | lncRNA:CR43280 | NO |
| FBgn0263404* | CG43450 | protein_coding_gene | 3R | 12195294 | 12195574 | - | CG43450 | NO |
| FBgn0264904 | CR44095 | antisense_lncRNA_gene | 3R | 14568962 | 14569871 | antisense RNA:CR44095 | asRNA:CR44095 | NO |
| FBgn0265893 | CR44682 | lncRNA_gene | 3L | 20634172 | 20634651 | long non-coding RNA:CR44682 | lncRNA:CR44682 | NO |
| FBgn0265896* | CR44685 | lncRNA_gene | 3L | 20652906 | 20654126 | long non-coding RNA:CR44685 | lncRNA:CR44685 | YES |
| FBgn0266347* | CG12414 | protein_coding_gene | 3L | 23206669 | 23287448 | nicotinic Acetylcholine Receptor alpha4 | nAChRalpha4 | YES |
| FBgn0266717* | CG6303 | protein_coding_gene | 3R | 10312039 | 10331353 | BIR repeat containing ubiquitin-conjugating enzyme | Bruce | NO |
| FBgn0266937* | CR45388 | lncRNA_gene | 3L | 18852573 | 18861363 | long non-coding RNA:CR45388 | lncRNA:CR45388 | NO |
| FBgn0267139* | CR45579 | lncRNA_gene | 3R | 11101180 | 11101684 | long non-coding RNA:CR45579 | lncRNA:CR45579 | YES |
| FBgn0267234 | CR45674 | lncRNA_gene | 3L | 21318223 | 21318650 | long non-coding RNA:CR45674 | lncRNA:CR45674 | YES |
| FBgn0267261 | CR45697 | lncRNA_gene | 2L | 17547185 | 17547849 | long non-coding RNA:CR45697 | lncRNA:CR45697 | NO |
| FBgn0267572* | CR45912 | antisense_lncRNA_gene | 3L | 18858690 | 18866107 | antisense RNA:CR45912 | asRNA:CR45912 | YES |
| FBgn0267737 | CR46068 | antisense_lncRNA_gene | 3L | 21333010 | 21333511 | antisense RNA:CR46068 | asRNA:CR46068 | YES |
| FBti0019369 | - | transposable_element | 3R | 13517303 | 13520034 | - | jockey{}1354 | YES |
| FBti0020284* | - | transposable_element | 3L | 23221385 | 23224264 | - | X{}1173 | YES |
| FBti0059658* | - | transposable_element | 3L | 23219958 | 23220245 | - | X{}1597 | YES |
| FBti0059667* | - | transposable_element | 3L | 23219852 | 23224264 | - | X{}1606 | YES |
| FBti0061816 | - | transposable_element | 3L | 23216353 | 23216596 | - | INE-1{}3755 | YES |
| FBti0061817 | - | transposable_element | 3L | 23216718 | 23216758 | - | INE-1{}3756 | YES |
| FBti0061819 | - | transposable_element | 3L | 23218788 | 23219072 | - | INE-1{}3758 | YES |
| FBti0061822 | - | transposable_element | 3L | 23225809 | 23226574 | - | INE-1{}3761 | NO |
| FBti0061826 | - | transposable_element | 3L | 23226793 | 23226970 | - | INE-1{}3765 | NO |
| FBti0061830 | - | transposable_element | 3L | 23228423 | 23228527 | - | INE-1{}3769 | NO |
| FBti0061859 | - | transposable_element | 3L | 23215676 | 23215900 | - | gypsy{}3798 | YES |
| FBti0061868 | - | transposable_element | 3L | 23219681 | 23219797 | - | 297{}3807 | YES |
| FBti0062073 | - | transposable_element | 3R | 14565185 | 14565228 | - | INE-1{}4012 | NO |
| FBti0063902 | - | transposable_element | 3L | 20641114 | 20641206 | - | INE-1{}5841 | NO |
| FBti0063903 | - | transposable_element | 3L | 20659518 | 20659809 | - | INE-1{}5842 | NO |
| FBti0064026 | - | transposable_element | 3L | 21426786 | 21426855 | - | INE-1{}5965 | YES |
| FBtr0304507 | - | - | - | - | - | - | - | NO |
| FBtr0472757 | - | - | - | - | - | - | - | NO |

**Table S12.** Spearman rank correlations between temporal allele frequency change and initial divergence from unexposed populations. Rank correlation coefficients (ρ) comparing sign-corrected GLM −log_10_(*P*) values from temporal contrasts within unexposed or pre-adapted populations against sign-corrected GLM −log_10_(*P*) values from the ‘unexposed vs. pre-adapted’ contrast in August. Positive correlations indicate that allele frequencies moved toward spinosad-adaptive alleles over time, whereas negative correlations indicate movement toward non-adaptive alleles. Time points are T1 = August 24, T2 = September 30, T3 = October 27, and T4 = November 24, 2023. Asterisks denote significant negative correlations (*P* < 0.05).

| **temporal contrast** | **pre-adapted** | **unexposed** |
|---|---|---|
| T1 vs. T2 | -0.1274* | 0.0493 |
| T1 vs. T3 | -0.3087* | -0.1248* |
| T1 vs. T4 | -0.2765* | -0.1213* |
| T2 vs. T3 | -0.3110* | -0.2036* |
| T2 vs. T4 | -0.2685* | -0.1790* |
| T3 vs. T4 | -0.1447* | -0.0438* |

**Table S13.** Directional change at top GLM outlier SNPs relative to matched background SNPs. One-tailed Welch *t*-tests of whether sign-corrected −log_10_(*P*) values from temporal GLM contrasts are lower for the top 10,000 outlier SNPs, ranked by GLM FDR in focal ‘unexposed vs. exposed’ contrasts, than for matched background SNPs, indicating allele frequency change in the direction of unexposed allelic change over time. Each outlier SNP was matched to a randomly drawn non-outlier SNP on the same chromosome arm with the same predicted variant consequence, a founder allele frequency within 25%, and a position more than 50 kb away; matched sets were drawn 100 times. Top outliers were selected from the ‘unexposed vs. pre-adapted’, ‘unexposed vs. extinct’, and ‘unexposed vs. rescue’ contrasts in August and the ‘unexposed vs. rescue’ contrast in November (SNP sets PAvE.T1, EvSE.T1, EvSP.T1, and EvSP.T4, respectively). Results are shown for temporal contrasts within unexposed and pre-adapted populations. Values are medians across the 100 iterations, except ‘perc. sig’, which gives the percentage of iterations with P < 0.05. Time points are T1 = August 24, T2 = September 30, T3 = October 27, and T4 = November 24, 2023.

| **SNP set** | **treatment** | **temporal contrast** | **focal SNPs: median signed −log10(*P*)** | **matched SNPs: median signed −log10(*P*)** | **median *t*-test statistic** | **median *t*-test P-value** | **perc. sig** |
|---|---|---|---|---|---|---|---|
| PAvE.T1_10K | unexposed | T1 vs. T2 | 0.006 | -0.0188 | 6.9114 | 1.00E+00 | 0 |
| PAvE.T1_10K | unexposed | T1 vs. T3 | -0.1786 | -0.1043 | -17.9306 | 1.49E-71 | 100 |
| PAvE.T1_10K | unexposed | T1 vs. T4 | -0.5626 | -0.2106 | -37.6949 | 2.51E-300 | 100 |
| PAvE.T1_10K | unexposed | T2 vs. T3 | -0.1306 | -0.0412 | -31.8893 | 2.87E-217 | 100 |
| PAvE.T1_10K | unexposed | T2 vs. T4 | -0.4171 | -0.1046 | -45.7383 | 0.00E+00 | 100 |
| PAvE.T1_10K | unexposed | T3 vs. T4 | -0.1107 | -0.0228 | -21.7756 | 3.42E-104 | 100 |
| PAvE.T1_10K | pre-adapted | T1 vs. T2 | 0.2371 | -0.1038 | 38.8365 | 1.00E+00 | 0 |
| PAvE.T1_10K | pre-adapted | T1 vs. T3 | -0.255 | -0.2306 | -2.537 | 5.60E-03 | 85 |
| PAvE.T1_10K | pre-adapted | T1 vs. T4 | -0.5577 | -0.5022 | -2.3152 | 1.03E-02 | 79 |
| PAvE.T1_10K | pre-adapted | T2 vs. T3 | -0.3175 | -0.0892 | -65.4798 | 0.00E+00 | 100 |
| PAvE.T1_10K | pre-adapted | T2 vs. T4 | -0.7363 | -0.2779 | -36.9585 | 1.26E-288 | 100 |
| PAvE.T1_10K | pre-adapted | T3 vs. T4 | -0.0399 | -0.0556 | 3.0615 | 9.99E-01 | 0 |
| EvSE.T1_10K | unexposed | T1 vs. T2 | 0.0014 | -0.0233 | 7.2357 | 1.00E+00 | 0 |
| EvSE.T1_10K | unexposed | T1 vs. T3 | -0.1632 | -0.1401 | -5.3652 | 4.11E-08 | 100 |
| EvSE.T1_10K | unexposed | T1 vs. T4 | -0.5717 | -0.2432 | -36.8776 | 8.74E-288 | 100 |
| EvSE.T1_10K | unexposed | T2 vs. T3 | -0.0738 | -0.0448 | -14.1191 | 2.48E-45 | 100 |
| EvSE.T1_10K | unexposed | T2 vs. T4 | -0.324 | -0.1027 | -42.2599 | 0.00E+00 | 100 |
| EvSE.T1_10K | unexposed | T3 vs. T4 | -0.1061 | -0.019 | -27.2871 | 3.31E-161 | 100 |
| EvSE.T1_10K | pre-adapted | T1 vs. T2 | 0.0801 | -0.0359 | 13.5387 | 1.00E+00 | 0 |
| EvSE.T1_10K | pre-adapted | T1 vs. T3 | -0.3818 | -0.1596 | -22.2996 | 1.55E-108 | 100 |
| EvSE.T1_10K | pre-adapted | T1 vs. T4 | -0.9983 | -0.3061 | -29.3344 | 2.96E-184 | 100 |
| EvSE.T1_10K | pre-adapted | T2 vs. T3 | -0.3949 | -0.0903 | -92.0261 | 0.00E+00 | 100 |
| EvSE.T1_10K | pre-adapted | T2 vs. T4 | -1.0454 | -0.2198 | -67.7194 | 0.00E+00 | 100 |
| EvSE.T1_10K | pre-adapted | T3 vs. T4 | -0.1087 | -0.0354 | -16.1617 | 1.43E-58 | 100 |
| EvSP.T1_10K | unexposed | T1 vs. T2 | 0.18 | 0.1118 | 12.4667 | 1.00E+00 | 0 |
| EvSP.T1_10K | unexposed | T1 vs. T3 | 0.1453 | 0.1004 | 8.5856 | 1.00E+00 | 0 |
| EvSP.T1_10K | unexposed | T1 vs. T4 | 0.1884 | 0.1224 | 7.8264 | 1.00E+00 | 0 |
| EvSP.T1_10K | unexposed | T2 vs. T3 | -0.0361 | -0.0181 | -3.0043 | 1.33E-03 | 96 |
| EvSP.T1_10K | unexposed | T2 vs. T4 | -0.0766 | -0.033 | -4.5361 | 2.89E-06 | 100 |
| EvSP.T1_10K | unexposed | T3 vs. T4 | -0.0144 | -0.0056 | -1.921 | 2.74E-02 | 61 |
| EvSP.T1_10K | pre-adapted | T1 vs. T2 | 0.0997 | -0.0118 | 10.2623 | 1.00E+00 | 0 |
| EvSP.T1_10K | pre-adapted | T1 vs. T3 | 0.1446 | 0.0419 | 8.6594 | 1.00E+00 | 0 |
| EvSP.T1_10K | pre-adapted | T1 vs. T4 | 0.3905 | 0.1191 | 9.2778 | 1.00E+00 | 0 |
| EvSP.T1_10K | pre-adapted | T2 vs. T3 | 0.0281 | 0.0206 | 2.4895 | 9.94E-01 | 0 |
| EvSP.T1_10K | pre-adapted | T2 vs. T4 | 0.1982 | 0.0918 | 7.7201 | 1.00E+00 | 0 |
| EvSP.T1_10K | pre-adapted | T3 vs. T4 | 0.0707 | 0.0196 | 7.2535 | 1.00E+00 | 0 |
| EvSP.T4_10K | unexposed | T1 vs. T2 | -0.0289 | -0.0148 | -3.2265 | 6.30E-04 | 99 |
| EvSP.T4_10K | unexposed | T1 vs. T3 | -0.0472 | -0.0171 | -6.8728 | 3.27E-12 | 100 |
| EvSP.T4_10K | unexposed | T1 vs. T4 | -0.3964 | -0.1024 | -31.6733 | 1.44E-214 | 100 |
| EvSP.T4_10K | unexposed | T2 vs. T3 | -0.0341 | -0.0054 | -8.2574 | 7.97E-17 | 100 |
| EvSP.T4_10K | unexposed | T2 vs. T4 | -0.3082 | -0.066 | -30.7797 | 2.02E-202 | 100 |
| EvSP.T4_10K | unexposed | T3 vs. T4 | -0.1997 | -0.0477 | -31.399 | 5.51E-211 | 100 |
| EvSP.T4_10K | pre-adapted | T1 vs. T2 | 0.4036 | 0.1324 | 22.7852 | 1.00E+00 | 0 |
| EvSP.T4_10K | pre-adapted | T1 vs. T3 | 0.5804 | 0.1348 | 29.5446 | 1.00E+00 | 0 |
| EvSP.T4_10K | pre-adapted | T1 vs. T4 | 1.3297 | 0.3673 | 29.5082 | 1.00E+00 | 0 |
| EvSP.T4_10K | pre-adapted | T2 vs. T3 | 0.0283 | -0.0057 | 11.8289 | 1.00E+00 | 0 |
| EvSP.T4_10K | pre-adapted | T2 vs. T4 | 0.3195 | 0.069 | 20.5495 | 1.00E+00 | 0 |
| EvSP.T4_10K | pre-adapted | T3 vs. T4 | 0.1564 | 0.0612 | 16.0046 | 1.00E+00 | 0 |

## References

1. M. Exposito-Alonso, T. R. Booker, L. Czech, L. Gillespie, S. Hateley, C. C. Kyriazis, P. L. M. Lang, L. Leventhal, D. Nogues-Bravo, V. Pagowski, M. Ruffley, J. P. Spence, S. E. Toro Arana, C. L. Weiß, E. Zess, Genetic diversity loss in the Anthropocene. Science 377, 1431–1435 (2022).

2. R. H. Cowie, P. Bouchet, B. Fontaine, The Sixth Mass Extinction: fact, fiction or speculation? Biol. Rev. Camb. Philos. Soc. 97, 640–663 (2022).

3. M. C. Urban, Climate change. Accelerating extinction risk from climate change. Science 348, 571–573 (2015).

4. G. Bell, A. Gonzalez, Evolutionary rescue can prevent extinction following environmental change. Ecol. Lett. 12, 942–948 (2009).

5. S. M. Carlson, C. J. Cunningham, P. A. H. Westley, Evolutionary rescue in a changing world. Trends Ecol. Evol. 29, 521–530 (2014).

6. G. Bell, Evolutionary rescue. Annu. Rev. Ecol. Evol. Syst. 48, 605–627 (2017).

7. A. Gonzalez, O. Ronce, R. Ferriere, M. E. Hochberg, Evolutionary rescue: an emerging focus at the intersection between ecology and evolution. Philos. Trans. R. Soc. Lond. B Biol. Sci. 368, 20120404 (2013).

8. D. N. Anstett, J. Anstett, S. N. Sheth, D. R. Moxley, H. A. Branch, M. Jahani, K. Huang, M. Todesco, R. Jordan, J. M. Lazaro-Guevara, L. H. Rieseberg, A. L. Angert, Rapid evolution predicts demographic recovery after extreme drought. Science 391, 1172–1176 (2026).

9. L. J. Balstad, M. L. Baskett, S. M. Carlson, C. J. Cunningham, P. A. H. Westley, Revisiting evolutionary rescue in the wild. Trends Ecol. Evol. 41, 762–774 (2026).

10. S. N. Aitken, M. C. Whitlock, Assisted gene flow to facilitate local adaptation to climate change. Annu. Rev. Ecol. Evol. Syst. 44, 367–388 (2013).

11. M. J. H. van Oppen, J. K. Oliver, H. M. Putnam, R. D. Gates, Building coral reef resilience through assisted evolution. Proc. Natl. Acad. Sci. U. S. A. 112, 2307–2313 (2015).

12. R. Gomulkiewicz, R. D. Holt, When does evolution by natural selection prevent extinction? Evolution 49, 201 (1995).

13. H. A. Orr, R. L. Unckless, Population extinction and the genetics of adaptation. Am. Nat. 172, 160–169 (2008).

14. H. Uecker, M. Osmond, C. Alejandre, E. Barrios-Caro, D. Clement, P. Czuppon, L. Dekens, P. Deraje, J. Draghi, A. Hasan, R. D. Holt, T. Kuosmanen, A. Longcamp, L. Marrec, G. Martin, C. Nyhoegen, M. Orive, V. R. Kumar, O. Ronce, R. Stana, L. Wahl, M. Yamamichi, Modeling evolutionary rescue. doi: 10.32942/X25D4T (2026).

15. M. C. Urban, J. Swaegers, R. Stoks, R. R. Snook, S. P. Otto, D. W. A. Noble, M. Moiron, M. H. Hällfors, M. Gómez-Llano, S. Fior, J. Cote, A. Charmantier, E. Bestion, D. Berger, J. Baur, J. M. Alexander, M. Saastamoinen, A. H. Edelsparre, C. Teplitsky, When and how can we predict adaptive responses to climate change? Evolution Letters 8, 172–187 (2024).

16. A. Gonzalez, G. Bell, Evolutionary rescue and adaptation to abrupt environmental change depends upon the history of stress. Philos. Trans. R. Soc. Lond. B Biol. Sci. 368, 20120079 (2013).

17. M. Lagator, N. Colegrave, P. Neve, Selection history and epistatic interactions impact dynamics of adaptation to novel environmental stresses. Proc. Biol. Sci. 281, 20141679 (2014).

18. G. Bell, A. Gonzalez, Adaptation and evolutionary rescue in metapopulations experiencing environmental deterioration. Science 332, 1327–1330 (2011).

19. J. Lachapelle, G. Bell, Evolutionary rescue of sexual and asexual populations in a deteriorating environment: Evolutionary rescue of sexual and asexual populations. Evolution 66, 3508–3518 (2012).

20. G. G. Perron, A. Gonzalez, A. Buckling, Source-sink dynamics shape the evolution of antibiotic resistance and its pleiotropic fitness cost. Proc. Biol. Sci. 274, 2351–2356 (2007).

21. L. Olazcuaga, R. A. Hufbauer, Evolution fails to rescue a population in an increasingly variable environment. Proc. Natl. Acad. Sci. U. S. A. 121, e2414877121 (2024).

22. T. J. Clark-Wolf, P. D. Boersma, F. Plard, G. A. Rebstock, B. Abrahms, Increasing environmental variability inhibits evolutionary rescue in a long-lived vertebrate. Proc. Natl. Acad. Sci. U. S. A. 121, e2406314121 (2024).

23. K. M. Quigley, M. Marzonie, B. Ramsby, D. Abrego, G. Milton, M. J. H. van Oppen, L. K. Bay, Variability in fitness trade-offs amongst coral juveniles with mixed genetic backgrounds held in the wild. Front. Mar. Sci. 8, 636177 (2021).

24. L. M. Bono, L. B. Smith Jr, D. W. Pfennig, C. L. Burch, The emergence of performance trade-offs during local adaptation: insights from experimental evolution. Mol. Ecol. 26, 1720–1733 (2017).

25. J. Ågren, C. G. Oakley, J. K. McKay, J. T. Lovell, D. W. Schemske, Genetic mapping of adaptation reveals fitness tradeoffs in Arabidopsis thaliana. Proc. Natl. Acad. Sci. U. S. A. 110, 21077–21082 (2013).

26. N. Barghi, J. Hermisson, C. Schlötterer, Polygenic adaptation: a unifying framework to understand positive selection. Nat. Rev. Genet. 21, 769–781 (2020).

27. J. M. Kreiner, The genetic architecture and spatiotemporal dynamics of adaptation across human-modified landscapes. New Phytol. 249, 744–750 (2026).

28. H. A. Orr, R. L. Unckless, The population genetics of evolutionary rescue. PLoS Genet. 10, e1004551 (2014).

29. O. Tenaillon, The utility of fisher’s geometric model in evolutionary genetics. Annu. Rev. Ecol. Evol. Syst. 45, 179–201 (2014).

30. G. Lee, B. J. Sanderson, T. J. Ellis, B. P. Dilkes, J. K. McKay, J. Ågren, C. G. Oakley, A large-effect fitness trade-off across environments is explained by a single mutation affecting cold acclimation. Proc. Natl. Acad. Sci. U. S. A. 121, e2317461121 (2024).

31. M. Kimura, On the probability of fixation of mutant genes in a population. Genetics 47, 713–719 (1962).

32. M. Lynch, R. Lande, Evolution and extinction in response to environmental change. Biotic interactions and global change (1993).

33. Y. McDonough, T. Connallon, Effects of population size change on the genetics of adaptation following an abrupt change in environment. Evolution 77, 1852–1863 (2023).

34. S. P. Otto, M. C. Whitlock, The probability of fixation in populations of changing size. Genetics 146, 723–733 (1997).

35. J. S. Andon, C. D. Kocher, K. A. Dill, T. Wang, “Excluded phenotypes” restrict genetic paths toward adaptation in declining populations. Proc. Natl. Acad. Sci. U. S. A. 123, e2507223123 (2026).

36. P. W. Messer, S. P. Ellner, N. G. Hairston Jr, Can Population Genetics Adapt to Rapid Evolution? Trends Genet. 32, 408–418 (2016).

37. J. Hemingway, H. Ranson, Insecticide resistance in insect vectors of human disease. Annu. Rev. Entomol. 45, 371–391 (2000).

38. J. A. Tennessen, R. Brosula, E. Chabanol, S. Bickersmith, A. M. Early, M. Laws, K. A. Kelley, M. E. Grillet, D. Gamboa, E. R. Lucas, J.-B. Duchemin, M. L. Quiñones, M. A. M. Sallum, E. S. Bergo, J. E. Moreno, S. Nagi, N. J. Arisco, M. Sooklall, R. Niles-Robin, M. C. Castro, H. Cox, M. Gendrin, J. E. Conn, D. E. Neafsey, Population genomics of Anopheles darlingi, the principal South American malaria vector mosquito. Science 391, 1373–1378 (2026).

39. J. Mallet, The evolution of insecticide resistance: Have the insects won? Trends Ecol. Evol. 4, 336–340 (1989).

40. J. M. Kreiner, S. M. Latorre, H. A. Burbano, J. R. Stinchcombe, S. P. Otto, D. Weigel, S. I. Wright, Rapid weed adaptation and range expansion in response to agriculture over the past two centuries. Science 378, 1079–1085 (2022).

41. M. Karageorgi, A. S. Lyulina, M. C. Bitter, E. Lappo, S. I. Greenblum, Z. K. Mouza, C. T. Tran, A. V. Huynh, H. Oken, P. Schmidt, D. A. Petrov, Beneficial reversal of dominance maintains a large-effect resistance polymorphism under fluctuating insecticide selection. *Nat*. Ecol. Evol. 9, 2145–2160 (2025).

42. B. E. Gress, F. G. Zalom, Identification and risk assessment of spinosad resistance in a California population of Drosophila suzukii: Spinosad resistance in California Drosophila suzukii. Pest Manag. Sci. 75, 1270–1276 (2019).

43. S. M. Rudman, S. I. Greenblum, S. Rajpurohit, N. J. Betancourt, J. Hanna, S. Tilk, T. Yokoyama, D. A. Petrov, P. Schmidt, Direct observation of adaptive tracking on ecological time scales in Drosophila. Science 375, eabj7484 (2022).

44. S. M. Rudman, S. Greenblum, R. C. Hughes, S. Rajpurohit, O. Kiratli, D. B. Lowder, S. G. Lemmon, D. A. Petrov, J. M. Chaston, P. Schmidt, Microbiome composition shapes rapid genomic adaptation of *Drosophila melanogaster*. Proc. Natl. Acad. Sci. U. S. A. 116, 20025–20032 (2019).

45. T. F. C. Mackay, S. Richards, E. A. Stone, A. Barbadilla, J. F. Ayroles, D. Zhu, S. Casillas, Y. Han, M. M. Magwire, J. M. Cridland, M. F. Richardson, R. R. H. Anholt, M. Barrón, C. Bess, K. P. Blankenburg, M. A. Carbone, D. Castellano, L. Chaboub, L. Duncan, Z. Harris, M. Javaid, J. C. Jayaseelan, S. N. Jhangiani, K. W. Jordan, F. Lara, F. Lawrence, S. L. Lee, P. Librado, R. S. Linheiro, R. F. Lyman, A. J. Mackey, M. Munidasa, D. M. Muzny, L. Nazareth, I. Newsham, L. Perales, L.-L. Pu, C. Qu, M. Ràmia, J. G. Reid, S. M. Rollmann, J. Rozas, N. Saada, L. Turlapati, K. C. Worley, Y.-Q. Wu, A. Yamamoto, Y. Zhu, C. M. Bergman, K. R. Thornton, D. Mittelman, R. A. Gibbs, The Drosophila melanogaster Genetic Reference Panel. Nature 482, 173–178 (2012).

46. A. R. Gabidulin, S. M. Rudman, MLDAAPP: Machine Learning Data Acquisition for Assessing Population Phenotypes. G3, doi: 10.1093/g3journal/jkaf173 (2025).

47. R. S. Shahmohamadloo, A. R. Gabidulin, E. R. Andrews, S. M. Rudman, Microbiome evolution plays a secondary role in host rapid adaptation, bioRxivorg (2025)p. 2025.06. 27.661976.

48. N. M. van Straalen, M. E. Feder, Ecological and evolutionary functional genomics--how can it contribute to the risk assessment of chemicals? Environ. Sci. Technol. 46, 3–9 (2012).

49. V. Mustonen, M. Lässig, From fitness landscapes to seascapes: non-equilibrium dynamics of selection and adaptation. Trends Genet. 25, 111–119 (2009).

50. E. S. King, A. E. Stacy, D. T. Weaver, J. Maltas, R. Barker-Clarke, E. Dolson, J. G. Scott, Fitness seascapes are necessary for realistic modeling of the evolutionary response to drug therapy. Sci. Adv. 11, eadv1268 (2025).

51. J. Cairns, F. Borse, T. Mononen, T. Hiltunen, V. Mustonen, Strong selective environments determine evolutionary outcome in time-dependent fitness seascapes. Evol. Lett. 6, 266–279 (2022).

52. E. Svensson, R. Calsbeek, Eds., The Adaptive Landscape in Evolutionary Biology (Oxford University Press, London, England, 2012).

53. M. C. Bitter, S. Berardi, H. Oken, A. Huynh, E. Lappo, P. Schmidt, D. A. Petrov, Continuously fluctuating selection reveals fine granularity of adaptation. Nature 634, 389–396 (2024).

54. K. Jain, W. Stephan, Modes of rapid polygenic adaptation. Mol. Biol. Evol. 34, 3169–3175 (2017).

55. S. Yeaman, A. C. Gerstein, K. A. Hodgins, M. C. Whitlock, Quantifying how constraints limit the diversity of viable routes to adaptation. PLoS Genet. 14, e1007717 (2018).

56. G. Bell, Fluctuating selection: the perpetual renewal of adaptation in variable environments. Philos. Trans. R. Soc. Lond. B Biol. Sci. 365, 87–97 (2010).

57. T. C. Sparks, F. J. Wessels, T. Perry, M. J. Price, M. Siebert, D. Mann, Spinosyn resistance and cross-resistance–A 25 year review and analysis. Pestic. Biochem. Physiol. 210, 106363 (2025).

58. C. A. Tabuloc, C. R. Carlson, F. Ganjisaffar, C. C. Truong, C.-H. Chen, K. M. Lewald, S. Hidalgo, N. L. Nicola, C. E. Jones, A. A. Sial, F. G. Zalom, J. C. Chiu, Transcriptome analysis of Drosophila suzukii reveals molecular mechanisms conferring pyrethroid and spinosad resistance. Sci. Rep. 14, 19867 (2024).

59. T. Perry, J. A. McKenzie, P. Batterham, A Dalpha6 knockout strain of Drosophila melanogaster confers a high level of resistance to spinosad. Insect Biochem. Mol. Biol. 37, 184–188 (2007).

60. S. M. Rudman, J. J. Faber-Hammond, R. E. Sherman, R. S. Shahmohamadloo, S. I. Greenblum, H. A. MacMillan, P. D. Jeyasingh, P. S. Schmidt, Ecological stoichiometry and life history theory, not the identity of genomic variants, predict rapid adaptation, bioRxivorg (2025)p. 2025.11. 11.687836.

61. B. E. Tabashnik, Y. Carrière, Surge in insect resistance to transgenic crops and prospects for sustainability. Nat. Biotechnol. 35, 926–935 (2017).

62. N. J. Hawkins, C. Bass, A. Dixon, P. Neve, The evolutionary origins of pesticide resistance. Biol. Rev. Camb. Philos. Soc. 94, 135–155 (2019).

63. R. H. Ffrench-Constant, The molecular genetics of insecticide resistance. Genetics 194, 807–815 (2013).

64. J. Overgaard, A. A. Hoffmann, T. N. Kristensen, Assessing population and environmental effects on thermal resistance in Drosophila melanogaster using ecologically relevant assays. J. Therm. Biol. 36, 409–416 (2011).

65. J. R. David, L. O. Araripe, M. Chakir, H. Legout, B. Lemos, G. Pétavy, C. Rohmer, D. Joly, B. Moreteau, Male sterility at extreme temperatures: a significant but neglected phenomenon for understanding Drosophila climatic adaptations. J. Evol. Biol. 18, 838–846 (2005).

66. P. Klepsatel, D. Knoblochová, M. Dharanikota, Ľ. Vidlička, M. Gáliková, Developmental plasticity of thermal performance curve for reproduction in Drosophila melanogaster. Evolution 77, 2606–2618 (2023).

67. A. R. Gunderson, E. J. Armstrong, J. H. Stillman, Multiple stressors in a changing world: The need for an improved perspective on physiological responses to the dynamic marine environment. Ann. Rev. Mar. Sci. 8, 357–378 (2016).

68. R. S. Brennan, J. A. deMayo, H. G. Dam, M. Finiguerra, H. Baumann, V. Buffalo, M. H. Pespeni, Experimental evolution reveals the synergistic genomic mechanisms of adaptation to ocean warming and acidification in a marine copepod. Proc. Natl. Acad. Sci. U. S. A. 119, e2201521119 (2022).

69. M. L. Tissier, R. S. Shahmohamadloo, L. M. Guzman, Pesticide risk assessment in a changing world. Glob. Chang. Biol. 31, e70203 (2025).

70. J. Maynard Smith, “Haldane’s dilemma” and the rate of evolution. Nature 219, 1114–1116 (1968).

71. H. A. Lindsey, J. Gallie, S. Taylor, B. Kerr, Evolutionary rescue from extinction is contingent on a lower rate of environmental change. Nature 494, 463–467 (2013).

72. P. Gienapp, M. Lof, T. E. Reed, J. McNamara, S. Verhulst, M. E. Visser, Predicting demographically sustainable rates of adaptation: can great tit breeding time keep pace with climate change? Philos. Trans. R. Soc. Lond. B Biol. Sci. 368, 20120289 (2013).

73. J. B. S. Haldane, The cost of natural selection. Journal of Genetics 55, 511–524 (1957).

74. J. M. Smith, What determines the rate of evolution? Am. Nat. 110, 331–338 (1976).

75. J. T. Anderson, M. L. DeMarche, D. A. Denney, I. Breckheimer, J. Santangelo, S. M. Wadgymar, Adaptation and gene flow are insufficient to rescue a montane plant under climate change. Science 388, 525–531 (2025).

76. R. A. Martin, C. R. B. da Silva, M. P. Moore, S. E. Diamond, When will a changing climate outpace adaptive evolution? Wiley Interdiscip. Rev. Clim. Change 14 (2023).

77. M. C. Whitlock, Fixation of new alleles and the extinction of small populations: drift load, beneficial alleles, and sexual selection. Evolution 54, 1855–1861 (2000).

78. G. Bell, Evolutionary rescue and the limits of adaptation. Philos. Trans. R. Soc. Lond. B Biol. Sci. 368, 20120080 (2013).

79. M. Kimura, The Neutral Theory of Molecular Evolution (Cambridge University Press, 1983).

80. R. Lande, Genetics and demography in biological conservation. Science 241, 1455–1460 (1988).

81. R. Gargiulo, K. B. Budde, M. Heuertz, Mind the lag: understanding genetic extinction debt for conservation. Trends Ecol. Evol. 40, 228–237 (2025).

82. Y. E. Stuart, T. Veen, J. N. Weber, D. Hanson, M. Ravinet, B. K. Lohman, C. J. Thompson, T. Tasneem, A. Doggett, R. Izen, N. Ahmed, R. D. H. Barrett, A. P. Hendry, C. L. Peichel, D. I. Bolnick, Contrasting effects of environment and genetics generate a continuum of parallel evolution. Nature Ecology & Evolution 1, 0158 (2017).

83. S. J. Arnold, R. Bürger, P. A. Hohenlohe, B. C. Ajie, A. G. Jones, Understanding the evolution and stability of the G-matrix. Evolution 62, 2451–2461 (2008).

84. C. Bank, Epistasis and adaptation on fitness landscapes. Annu. Rev. Ecol. Evol. Syst. 53, 457–479 (2022).

85. J. Beets, J. Höglund, B. Y. Kim, J. Ellers, K. M. Hoedjes, M. Bosse, Predicting the functional impact of single nucleotide variants in *Drosophila melanogaster* with FlyCADD. Genetics 232, iyaf250 (2026).

86. P. J. Daborn, J. L. Yen, M. R. Bogwitz, G. Le Goff, E. Feil, S. Jeffers, N. Tijet, T. Perry, D. Heckel, P. Batterham, R. Feyereisen, T. G. Wilson, R. H. ffrench-Constant, A single p450 allele associated with insecticide resistance in Drosophila. Science 297, 2253–2256 (2002).

87. J. D. Gaitán-Espitia, A. J. Hobday, Evolutionary principles and genetic considerations for guiding conservation interventions under climate change. Glob. Chang. Biol. 27, 475–488 (2021).

88. M. M. Osmond, C. A. Klausmeier, An evolutionary tipping point in a changing environment. Evolution 71, 2930–2941 (2017).

89. M. Hartfield, S. P. Otto, Recombination and hitchhiking of deleterious alleles: Recombination and the undesirable hitchhiker. Evolution 65, 2421–2434 (2011).

90. P. C. Phillips, Epistasis--the essential role of gene interactions in the structure and evolution of genetic systems. Nat. Rev. Genet. 9, 855–867 (2008).

91. J. A. deMayo, R. S. Brennan, M. Pespeni, C. Jaspers, Ø. Varpe, C. E. Lee, H. G. Dam, Recognizing adaptation costs in the Anthropocene. Trends Ecol. Evol. 40, 1225–1233 (2025).

92. F. Courchamp, T. Clutton-Brock, B. Grenfell, Inverse density dependence and the Allee effect. Trends Ecol. Evol. 14, 405–410 (1999).

93. S. Ellner, N. G. Hairston Jr, Role of overlapping generations in maintaining genetic variation in a fluctuating environment. Am. Nat. 143, 403–417 (1994).

94. R. Lande, Risks of population extinction from demographic and environmental stochasticity and random catastrophes. Am. Nat. 142, 911–927 (1993).

95. S. W. Nordstrom, R. A. Hufbauer, L. Olazcuaga, L. F. Durkee, B. A. Melbourne, How density dependence, genetic erosion and the extinction vortex impact evolutionary rescue. Proc. Biol. Sci. 290, 20231228 (2023).

96. K. Xu, M. M. Osmond, When does the probability of evolutionary rescue increase with the strength of selection despite a potential demographic cost? Am. Nat. 207, 615–626 (2026).

97. V. Radchuk, T. Reed, C. Teplitsky, M. van de Pol, A. Charmantier, C. Hassall, P. Adamík, F. Adriaensen, M. P. Ahola, P. Arcese, J. Miguel Avilés, J. Balbontin, K. S. Berg, A. Borras, S. Burthe, J. Clobert, N. Dehnhard, F. de Lope, A. A. Dhondt, N. J. Dingemanse, H. Doi, T. Eeva, J. Fickel, I. Filella, F. Fossøy, A. E. Goodenough, S. J. G. Hall, B. Hansson, M. Harris, D. Hasselquist, T. Hickler, J. Joshi, H. Kharouba, J. G. Martínez, J.-B. Mihoub, J. A. Mills, M. Molina-Morales, A. Moksnes, A. Ozgul, D. Parejo, P. Pilard, M. Poisbleau, F. Rousset, M.-O. Rödel, D. Scott, J. C. Senar, C. Stefanescu, B. G. Stokke, T. Kusano, M. Tarka, C. E. Tarwater, K. Thonicke, J. Thorley, A. Wilting, P. Tryjanowski, J. Merilä, B. C. Sheldon, A. Pape Møller, E. Matthysen, F. Janzen, F. S. Dobson, M. E. Visser, S. R. Beissinger, A. Courtiol, S. Kramer-Schadt, Adaptive responses of animals to climate change are most likely insufficient. Nat. Commun. 10, 3109 (2019).

98. X. Wu, T. Bellagio, Y. Peng, L. Czech, M. Lin, P. Lang, R. Epstein, M. Abdelaziz, J. Alexander, C. Alonso-Blanco, H. L. Andersen, M. Berbel, J. Bergelson, O. Bossdorf, L. Burghardt, M. Caton-Darby, R. Colautti, C. Delker, P. G. Dimitrakopoulos, K. Donohue, W. Durka, G. Escribano-Avila, S. J. Franks, F. B. Fritschi, A. Galanidis, A. Garcia-Fernández, A. García-Muñoz, E. Hamann, A. Hutt, J. M. Iriondo, T. E. Juenger, S. R. Keller, K. Koehl, A. Korte, P. Korte, A. Kutschera, C. Lara-Romero, L. Leventhal, D. Maag, A. Marcer, M. March-Salas, J. de Meaux, B. Méndez-Vigo, J. Morente-López, T. C. Morton, Z. Münzbergova, A. Muola, H. A. Nomoto, M. Pärtel, F. X. Picó, B. Quarles-Chidyagwai, M. Quint, N. Reichelt, A. Rudak, J. Schmitt, G. Schmitz, M. Seifan, B. L. Snoek, R. Stam, M. Stift, J. R. Stinchcombe, M. A. Taylor, P. Tiffin, I. Till-Bottraud, A. Traveset, J.-G. Valay, M. Van Zanten, V. Vandvik, C. Violle, D. Weigel, M. Wódkiewicz, F. Vasseur, J. F. Scheepens, M. Exposito-Alonso, Rapid adaptation and extinction in synchronized outdoor evolution experiments of Arabidopsis. Science 391, eadz0777 (2026).

99. M. Exposito-Alonso, 500 Genomes Field Experiment Team, H. A. Burbano, O. Bossdorf, R. Nielsen, D. Weigel, Publisher Correction: Natural selection on the Arabidopsis thaliana genome in present and future climates. Nature 574, E16 (2019).

100. T. Capblancq, M. C. Fitzpatrick, R. A. Bay, M. Exposito-Alonso, S. R. Keller, Genomic Prediction of (Mal)Adaptation Across Current and Future Climatic Landscapes. doi: 10.1146/annurev-ecolsys-020720-042553 (2020).

101. S. M. Rudman, M. A. Barbour, K. Csilléry, P. Gienapp, F. Guillaume, N. G. Hairston Jr, A. P. Hendry, J. R. Lasky, M. Rafajlović, K. Räsänen, P. S. Schmidt, O. Seehausen, N. O. Therkildsen, M. M. Turcotte, J. M. Levine, What genomic data can reveal about eco-evolutionary dynamics. Nat Ecol Evol 2, 9–15 (2018).

102. R. A. Bay, S. R. Palumbi, Multilocus adaptation associated with heat resistance in reef-building corals. Curr. Biol. 24, 2952–2956 (2014).

103. S. N. Aitken, J. B. Bemmels, Time to get moving: assisted gene flow of forest trees. Evol. Appl. 9, 271–290 (2016).

104. S. W. Fitzpatrick, J. C. Gerberich, L. M. Angeloni, L. L. Bailey, E. D. Broder, J. Torres-Dowdall, C. A. Handelsman, A. López-Sepulcre, D. N. Reznick, C. K. Ghalambor, W. Chris Funk, Gene flow from an adaptively divergent source causes rescue through genetic and demographic factors in two wild populations of Trinidadian guppies. Evol. Appl. 9, 879–891 (2016).

105. R. A. Knapp, M. Q. Wilber, M. B. Joseph, T. C. Smith, R. L. Grasso, Reintroduction of resistant frogs facilitates landscape-scale recovery in the presence of a lethal fungal disease. Nat. Commun. 15, 9436 (2024).

106. A. Q. Byrne, A. P. Rothstein, R. A. Knapp, E. B. Rosenblum, Genomic correlates of disease recovery in natural populations of the Sierra Nevada yellow-legged frog (Rana sierrae). Mol. Ecol. 35, e70348 (2026).

107. T. J. Kawecki, D. Ebert, Conceptual issues in local adaptation. Ecol. Lett. 7, 1225–1241 (2004).

108. A. A. Hoffmann, A. Anderson, R. Hallas, Opposing clines for high and low temperature resistance in *Drosophila melanogaster*. Ecol. Lett. 5, 614–618 (2002).

109. L. Czech, M. Exposito-Alonso, Grenepipe: A flexible, scalable and reproducible pipeline to automate variant calling from sequence reads. Bioinformatics (2022).

110. S. Chen, Y. Zhou, Y. Chen, J. Gu, fastp: an ultra-fast all-in-one FASTQ preprocessor. Bioinformatics 34, i884–i890 (2018).

111. Vasimuddin, S. Misra, H. Li, S. Aluru, Efficient architecture-aware acceleration of BWA-MEM for multicore systems. IPDPS, 314–324 (2019).

112. Broad Institute, Picard: A Set of Command Line Tools (in Java) for Manipulating High-Throughput Sequencing (HTS) Data and Formats Such as SAM/BAM/CRAM and VCF (GitHub Repository, 2019; https://github.com/broadinstitute/picard).

113. P. Danecek, J. K. Bonfield, J. Liddle, J. Marshall, V. Ohan, M. O. Pollard, A. Whitwham, T. Keane, S. A. McCarthy, R. M. Davies, H. Li, Twelve years of SAMtools and BCFtools. Gigascience 10, giab008 (2021).

114. S. Tilk, A. Bergland, A. Goodman, P. Schmidt, D. Petrov, S. Greenblum, Accurate allele frequencies from ultra-low coverage pool-seq samples in evolve-and-resequence experiments. G3: Genes, Genomes, Genetics 9, 4159–4168 (2019).

115. A. D. Long, K. M. Hanson, S. J. Macdonald, The illusion of polygenicity in pool-seq genetic mapping studies: Insufficient power can mask simple genetic architectures. Genetics, iyag068 (2026).

116. P. Ewels, M. Magnusson, S. Lundin, M. Käller, MultiQC: summarize analysis results for multiple tools and samples in a single report. Bioinformatics 32, 3047–3048 (2016).

117. D. Bates, M. Machler, B. Bolker, S. C. Walker, Fitting linear mixed-effects models using lme4. Journal of Statistical Software 67, 1–48 (2015).

118. A. Kuznetsova, P. B. Brockhoff, R. H. B. Christensen. R package version (2015).

119. R. Lenth, H. Singmann, J. Love, P. Buerkner, M. Herve, Emmeans: Estimated marginal means, aka least-squares means (R package version 1.5. 1.)[Computer software]. (2021).

120. Y. Benjamini, Y. Hochberg, Controlling the False Discovery Rate: A Practical and Powerful Approach to Multiple Testing. J. R. Stat. Soc. Series B Stat. Methodol. 57, 289–300 (1995).

121. D. Vaughan, slider: Sliding window functions. R package version 0.3. 1. (2023).

122. L. Czech, J. P. Spence, M. Expósito-Alonso, Grenedalf: Population genetic statistics for the next generation of pool sequencing, arXiv [q-bio.PE] (2023). http://arxiv.org/abs/2306.11622.

123. C. Burny, V. Nolte, M. Dolezal, C. Schlötterer, Highly parallel genomic selection response in replicated *Drosophila melanogaster* populations with reduced genetic variation. Genome Biol. Evol. 13, evab239 (2021).

124. S. B. Plaisier, R. Taschereau, J. A. Wong, T. G. Graeber, Rank–rank hypergeometric overlap: identification of statistically significant overlap between gene-expression signatures. Nucleic acids research 38, e169–e169 (2010).

125. K. M. Cahill, Z. Huo, G. C. Tseng, R. W. Logan, M. L. Seney, Improved identification of concordant and discordant gene expression signatures using an updated rank-rank hypergeometric overlap approach. Sci. Rep. 8, 9588 (2018).

126. S. Maere, K. Heymans, M. Kuiper, BiNGO: a Cytoscape plugin to assess overrepresentation of gene ontology categories in biological networks. Bioinformatics 21, 3448–3449 (2005).

